# Secreted TAL effectors protect symbiotic bacteria from entrapment within fungal hyphae

**DOI:** 10.1101/2020.03.28.013177

**Authors:** Ingrid Richter, Zerrin Uzum, Claire E. Stanley, Nadine Moebius, Timothy P. Stinear, Sacha J. Pidot, Iuliia Ferling, Falk Hillmann, Christian Hertweck

**Affiliations:** Department of Biomolecular Chemistry, Leibniz Institute for Natural Product Research and Infection Biology (HKI), Jena, Germany; Plant – Soil Interactions, Agroecology and Environment Research Division, Agroscope, Zurich, Switzerland; Department of Microbiology and Immunology at the Doherty Institute for Infection and Immunity, University of Melbourne, Melbourne, Australia; Junior Research Group Evolution of Microbial Interactions, HKI, Jena, Germany; Faculty of Biological Sciences, Friedrich Schiller University Jena, Jena, Germany

## Abstract

The association of the agriculturally significant phytopathogenic fungus *Rhizopus microsporus* with the bacterial endosymbiont *Burkholderia rhizoxinica* is a remarkable example of bacteria controlling host physiology and reproduction. Here, we show that a group of transcription activator-like effectors (TALEs) called *Burkholderia* TALE-like proteins (BATs) from *B. rhizoxinica* are essential for the establishment of the symbiosis. Mutants lacking BAT proteins are unable to induce host sporulation. Utilising novel microfluidic devices in combination with fluorescence microscopy we observed the accumulation of BAT-deficient mutants in specific fungal side-hyphae with accompanying increased fungal re-infection. High-resolution live imaging revealed septa biogenesis at the base of infected hyphae leading to compartmental trapping of BAT-deficient endobacteria. Trapped endosymbionts showed reduced intracellular survival, suggesting a protective response from the fungal host against bacteria lacking specific effectors. These findings underscore the involvement of BAT proteins in maintaining a balance between mutualism and antagonism in bacterial-fungal interactions and provide deeper insights into the dynamic interactions between bacteria and eukaryotes.

---

Bacteria living in close association with eukaryotic hosts may control and exploit their host via pathogenic or mutualistic interactions^1^. For example, amplification of a genomic region in a normally beneficial *Wolbachia* symbiont leads to over-proliferation at the hosts’ expense^2^. Various bacteria control their eukaryotic hosts by translocating protein effector molecules into host cells. Therefore, they may use dedicated protein secretion systems (Type 1 to 7SS, Fig. 1a)^3,4^. For example, the type 3 secretion system (T3SS)-associated *Xanthomonas* AvrBs3 effector family (termed transcription activator-like effectors, TALEs) are highly orthologous proteins from bacterial plant pathogens that aid colonisation by mimicking plant transcription factors^5^. In humans, the gut pathogen *Helicobacter pylori* directly injects virulence factors into host cells. These T4SS-secreted proteins impair intracellular signalling systems, leading to morphological changes and increased inflammation in the host^6^.

**Fig. 1.**
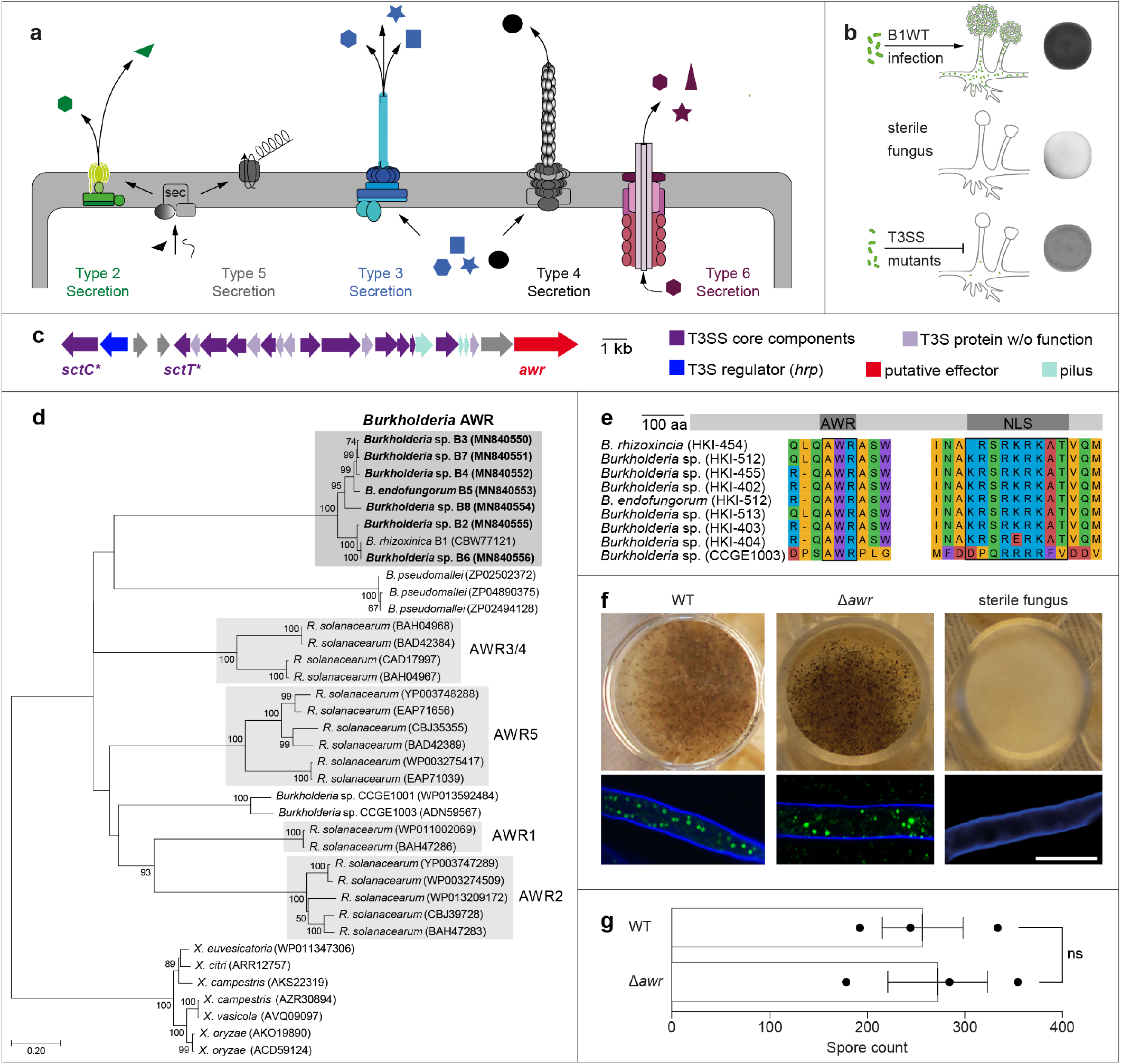
Characterisation and functional analysis of an effector protein associated with the type 3 secretion system (T3SS) from endofungal *Burkholderia* species. **a**, Structural organisation of bacterial (Gram-negative) secretion systems^3^. **b**, Schematic representation of *Rhizopus microsporus* containing *Burkholderia rhizoxinica*. Co-cultivation of isolated wild-type *B. rhizoxinica* (B1WT) with cured (endosymbiont-free) fungi leads to fungal reinfection, and the hosts’ ability to form sporangiospores is restored. Mature sporangia are absent in cured fungi. T3SS mutants can partially re-infect hyphae, but fail to cause sporulation^21^. **c**, Schematic representation of the T3SS gene cluster (*sct*) located on plasmid pBR01 that encodes for the secretion apparatus^21^. The putative T3 effector-encoding gene (*awr*) is indicated in red. T3SS mutant strains (Δ*sctC*::Kan^r^ and Δ*sctT*::Kan^r^), used in this study, were generated previously^21^ (*). **d**, Phylogenetic tree of AWR homologs from plant pathogenic *R. solanaceraum* and *Burkholderia* species. The phylogenetic analyses were performed using MEGA7 (see Methods for details). Sequences obtained in this study are in bold. GenBank accession numbers are given in brackets (Supplementary Table 6). **e**, Illustration and multiple sequence alignment of the functional domains of AWR containing nuclear localisation sequences (NLS) from *Burkholderia* species. **f**, Photographs and fluorescence microscopy images of sporulation assay. After inoculation of *R. microsporus* hyphae with B1WT (left) or Δ*awr*::Kan^r^ mutant strain (middle), fungal cultures sporulate, indicating successful reinfection. Fluorescence microscopy confirmed the localisation of bacteria (green) inside the fungal hyphae (blue). The control (no bacteria added) showed no sporulation and no presence of endohyphal bacteria (right). Scale bar: 20 μm. **g**, Spore count after one week of cocultivation. N = 3 biological replicates (3 technical replicates) ± one SEM. One-way ANOVA with Tukey’s multiple comparison test (ns: not significant, *p>0.05*, Supplementary Table 2).

An intriguing case of bacteria controlling host physiology and reproduction is the endosymbiosis between the plant-pathogenic zygomycete *Rhizopus microsporus* and its toxin-producing bacterial endosymbiont *Burkholderia rhizoxinica^7,8^* (which was recently suggested to be placed in the novel genus *Mycetohabitans^9^*). In this agriculturally relevant symbiosis^10^, host reproduction through spores relies exclusively on the presence of endobacteria (Fig. 1b). If *R. microsporus* is cured of its endosymbionts, the host is unable to reproduce vegetatively^8^. This strict control of sporulation ensures that the symbiosis persists over time, because the endosymbionts are translocated into the fungal spores during host reproduction. Multiple investigations have shed light on various symbiotic factors. For example, bacteria can invade the fungal host by secreting effector proteins via the T2SS^11^, while a linear lipopeptide aids the reinfection process by reducing surface tension^12^. Once inside the fungal hyphae, *B. rhizoxinica* lives in “stealth mode” due to the presence of a specialised bacterial lipopeptide O-antigen^13^. In addition, a functional T3SS is required for the formation of a stable symbiosis^14^. While these findings imply that secreted effector molecules produced by bacterial endosymbionts control fungal physiology and development, there is a considerable lack of knowledge about the molecular basis of this interaction and the types of effectors involved.

Here we show that TALEs produced by the symbionts play a key role in controlling fungal reproduction in this phytopathogenic bacterial-fungal interaction (BFI). Furthermore, we present the first real-time snapshot of septa biogenesis in a zygomycete fungus leading to hyphal trapping of endosymbionts incapable of secreting TALEs.

## Results

### Investigation of a T3SS-associated effector

To identify genes coding for T3 effector proteins we sequenced the genomes of seven endofungal *Burkholderia* species (Supplementary Table 1). These seven strains form the *B. rhizoxinica* complex together with *B. rhizoxinica* HKI-454, whose genome sequence has been published previously (Supplementary Table 2)^15,16^. Within each draft genome sequence, we found a gene encoding for a potential T3SS effector, with an alanine-tryptophan-arginine triad (AWR) peptide (Fig. 1c). The *awr* gene family, present in a number of bacterial pathogens including phytopathogenic *Burkholderia* and *Ralstonia* strains, collectively contributes to bacterial virulence^17,18^. For example, deletion of all five *awr* genes in *Ralstonia solanacearum* severely impairs its capacity to multiply in the natural host plant^17^.

To confirm the identified sequences as genuine *awr* orthologues, a phylogeny was inferred from alignment of the predicted endofungal *Burkholderia* AWR peptide sequences (Fig. 1d). These sequences were aligned with homologous AWR sequences from Gram-negative plant and animal pathogens and other *Burkholderia* symbionts (Supplementary Fig. 1)^17^. The tree was rooted in the *Xanthomonas* (γ-proteobacteria) sequences, which are more distantly related to the *Ralstonia* and *Burkholderia* (β-proteobacteria) sequences. According to the inferred phylogeny, the nearest relative of the endofungal AWR proteins was found in the mammalian pathogen *Burkholderia pseudomallei* (GenBank acc. no. ZP02502372)^19^. In addition, all eight endofungal *Burkholderia* AWR peptide sequences contained monopartite nuclear localisation sequences (NLS) at the C-terminal end of the protein (Fig. 1e and Supplementary Table 3). These sequences are consistent with the previously formulated NLS consensus sequence K(K/R)X(K/R)^20^.

To test whether endobacterial AWRs are involved in fungal reproduction, we generated targeted deletion mutants (Δ*awr*::Kan^r^, Supplementary Fig. 2) using a double-crossover strategy^21^. AWR-deficient mutants were examined for their reinfection ability, using a previously described sporulation bioassay^21^. When wild-type *B. rhizoxinica* and the endosymbiont-free (cured) fungal host are cocultured, mature sporangiophores can be seen after four days (Fig. 1f). The appearance of mature sporangiophores indicates the successful establishment of the symbiosis. Similar numbers of spores were observed upon co-cultivation with *B. rhizoxinica* Δ*awr*::Kan^r^ leading to complete restoration of the wild-type phenotype (Fig. 1g and Supplementary Table 2). In addition, the intracellular localisation of the Δ*awr*::Kan^r^ mutant strain was comparable to wild-type *B. rhizoxinica* (Fig. 1f).

### Wide distribution of TALEs in endofungal symbionts

Since we were unable to observe an effect of AWR on fungal reproduction, we continued to search for putative T3 effectors by investigating the genomes of eight endofungal *Burkholderia* species associated with *R. microsporus* (Supplementary Table 1) using T3SS prediction tools (see Methods for details). Three putative effectors, encoded by genes with the locus tags RBRH_01844 (2,316 bp), RBRH_01777 (936 bp), and RBRH_01776 (2,994 bp) were previously identified in *B. rhizoxinica* HKI-454 (Fig. 2a).

**Fig. 2.**
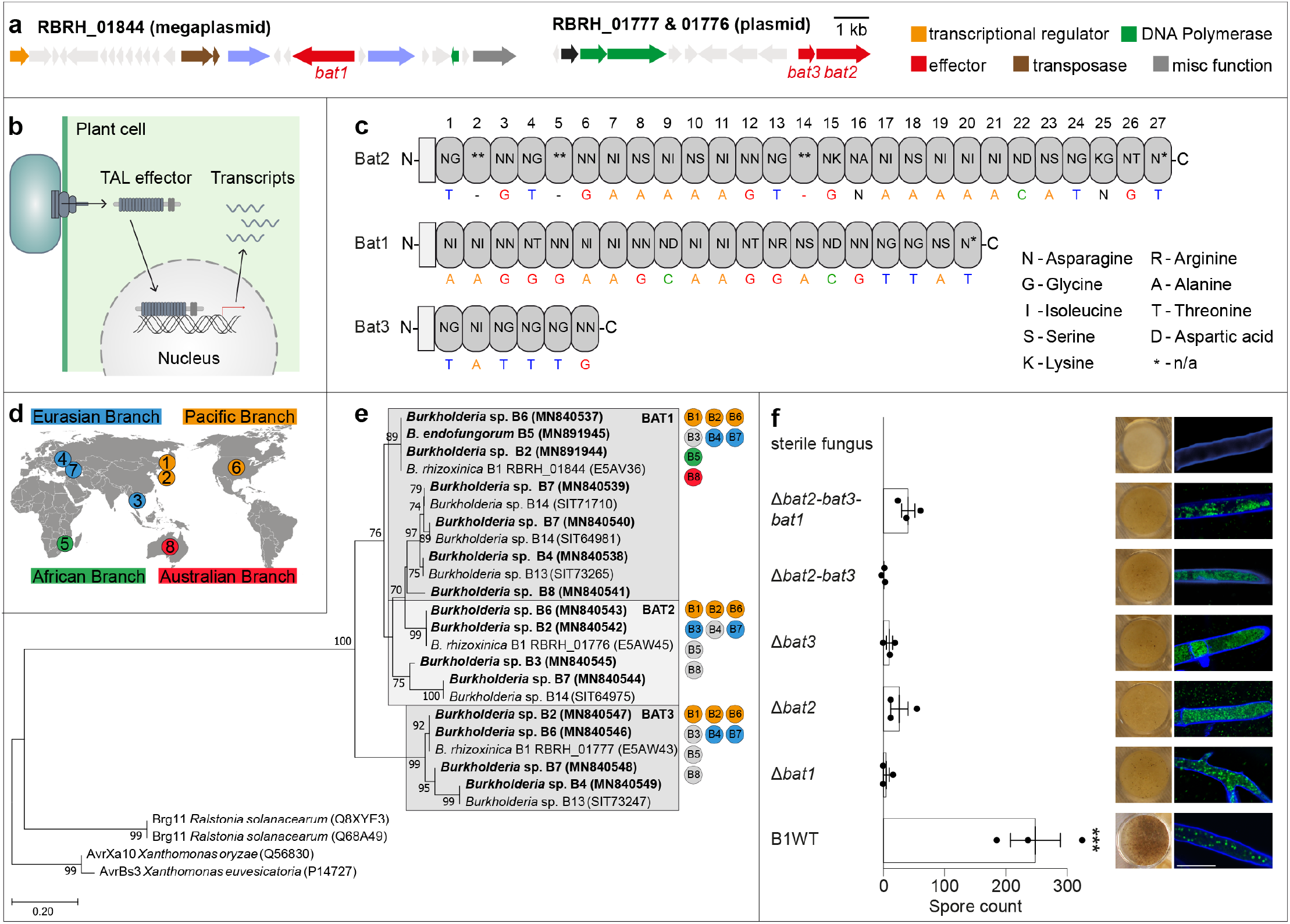
Identification and functional analysis of predicted transcription-activator like effectors (TALEs) from endofungal *Burkholderia* species (BATs). **a**, Schematic illustration of the gene clusters encoding BAT1 (RBRH_01844), BAT2 (RBRH_01776), and BAT3 (RBRH_01777) indicated in red. **b**, Mode of action of TALEs from *Xanthomonas* sp. TALEs are secreted into plant cells via the T3SS, translocate to the nucleus, and induce expression of target genes^22^. **c**, Schematic representation of the overall domain structure of BATs and the amino acid tandem repeats responsible for specification of the target nucleotide sequence^23^. **d**, Map depicting global distribution of endofungal *Burkholderia* strains and their classification into four branches^16^. **e**, Phylogenetic tree of TALE-like proteins from eight endofungal *Burkholderia* species, and plant pathogenic *Ralstonia solanaceraum* and *Xanthomonas* sp (Supplementary Fig. 3). Phylogenetic analysis was performed using MEGA7 (see Methods for details). BAT sequences obtained in this study are highlighted in bold and GenBank accession numbers are given in brackets (Supplementary Table 6). The distribution of BATs across the four *Burkholderia* branches is indicted as follows: orange: Pacific branch; blue: Eurasian branch; green: African branch; red: Australian branch; grey: not detected. **f**, Deletion of BATs decreases the sporulation ability of *R. microsporus* after re-infection. Photographs and spore count of *R. microsporus* re-infected with *B. rhizoxinica* wild-type (B1WT) or BAT mutant strains (Δ*bat1*::Apra^r^, Δ*bat2*::Kan^r^, Δ*bat3*::Kan^r^, Δ*bat2_bat3::Kan^r^*, or Δ*bat1::Apra^r^-Δbat2_bat3::Kan^r^*) after one week of co-cultivation. N = 3 biological replicates (3 technical replicates) ± one SEM. One-way ANOVA with Tukey’s multiple comparison test (*\*\*\*p<0.05*, Supplementary Table 4). Localisation of bacteria (green) inside the fungal hyphae (blue) was confirmed by fluorescence microscopy. Scale bar: 20 μm. The sterile fungus (no bacteria added) showed no sporulation and no presence of endohyphal bacteria.

As these genes show striking similarity to TAL effectors, they were named *Burkholderia* TALE-like effectors (BAT1: RBRH_01844; BAT2: RBRH_01776; BAT3: RBRH_01777, Fig. 2a)^23,25,26^. First identified in phytopathogenic bacteria of the genera *Xanthomonas* and *Ralstonia*, TALEs directly modify the expression of host plant genes to help bacterial colonisation (Fig. 2b)^27–31^. Similar to the *Xanthomonas* and *R. solanacearum* TALEs, BATs consist of numerous amino acid repeat units (BAT1: 22 repeats; BAT2: 27 repeats; BAT3: 6 repeats; Fig. 2c). Although slightly shorter than the *Xanthomonas* and *Ralstonia* repeats (34 repeats), BAT repeats contain a sequence-specific DNA binding motif following the canonical TALE code^23,25,27^.

The endofungal *bat* sequences identified in the genomes were confirmed as genuine TALE orthologues using: (i) PCR amplification and Sanger sequencing (Supplementary Table 5) and; (ii) phylogenetic analyses of predicted protein sequences. BAT1 was found in each of the five *Burkholderia* branches, while BAT2 and BAT3 were present in species belonging to the Eurasian and Pacific branch (Fig. 2d,e). It is conceivable that missing *bat* sequences evaded detection due to their highly repetitive nature. Using phylogenetic analysis, *R. solanacearum* TALEs (RipTALs) were identified as the closest relatives of endofungal BAT proteins (Fig. 2e and Supplementary Fig. 3).

TALEs usually harbour a T3S signal, a NLS, and an activation domain, all of which are required for the transcriptional activation of host plant genes^22^. We searched for NLS motifs within endofungal BAT peptide sequences using the NucPred and cNLS prediction software (see Methods for details). A fraction of BAT1 and BAT2 proteins were predicted to localise to the nucleus (52%) while the remaining fraction localises to the cytoplasm (48%). Localisation of BAT3 could not be predicted (Supplementary Table 7). Although a transcriptional activation domain (AD) seems to be absent in BAT proteins, a BAT1 fusion protein containing a viral NLS and AD was shown to transcriptionally activate a promoter with its target sequence in both human and plant cells^23^. However, their native function in *B. rhizoxinica*, or in the interaction with *R. microsporus*, is still unknown.

### Impact of BAT mutants on fungal sporulation

In order to identify the role of BAT proteins in the *Burkholderia* – *Rhizopus* symbiosis, we performed targeted gene deletion using a double-crossover strategy^21^. Although *B. rhizoxinica* can be maintained in axenic cultures under laboratory conditions, genetic manipulations are challenging. The long doubling time and the aggregation of cells^32^ significantly impair the selection process. Despite these challenges, we succeeded in deleting individual *bat* genes to generate Δ*bat1::Apra^r^*, Δ*bat2::Kan^r^*, and Δ*bat3::Kan^r^* mutant strains. Additionally, multi-deletions of *bat* genes were performed to generate double (Δ*bat2_bat3::Kan^r^*) and triple (Δ*bat1::Apra^r^-bat2_bat3::Kan^r^*) mutant strains (Supplementary Fig. 4).

Next, the reinfection potential of the knock-out strains was evaluated using a sporulation assay. The sporulation efficiency was significantly (*p<0.05*) reduced in all *B. rhizoxinica* BAT mutant strains as only a limited number of mature sporangia were formed (Fig. 2f and Supplementary Table 4). In contrast, the wild-type *B. rhizoxinica* strain readily triggered sporulation after three days of co-incubation. Fluorescence microscopy was used to monitor bacteria (stained with SYTO9) inside the fungal hyphae (counter-stained with calcofluor white) following re-infection. BAT mutants reached an extremely dense cell density within the host cytosol, while wild-type bacteria spread equally among the fungal mycelium (Fig. 2f).

### BAT-dependent changes in fungal morphology

To investigate the accumulation of BAT mutant cells in the fungal hyphae, we utilised a tailor-made microfluidic platform that allows live monitoring of BFIs over time using fluorescence microscopy (Fig. 3a)^24^. First, cured *R. microsporus* was grown in BFI devices filled with PDB (Fig. 3b). After two days of incubation, the fungus was coincubated with SYTO9-stained wild-type *B. rhizoxinica* or BAT mutant strains (Δ*bat1*::Apra^r^, Δ*bat2*::Kan_r_, and Δ*bat3*::Kan_r_). As a control, T3SS-deficient mutants (Δ*sctC*::Kan^r^ and Δ*sctT*::Kan^r^)^21^ or a rhizoxin-deficient mutant (Δ*rhiG*::Kan^r^) were also co-incubated. We observed four morphologically distinct types of hyphae (Fig. 3c and Supplementary Fig. 5), which were classified as follows: (1) vegetative side hyphae; (2) vegetative main hyphae; (3) vegetative empty side hyphae; and (4) abortive sporangiophore. Vegetative main hyphae (morphotype 2) are characterised by their relatively large diameter (app. 10–15 μm). They appear to be tightly packed with organelles (e.g. nuclei, vacuoles, mitochondria, etc.), which are transported through the hyphae at relatively high speed (Video 1). This hyphal classification has been described for members of the Basidiomycetes and Ascomycetes, but rarely in Zygomycetes^33,34^. We also observed smaller side hyphae, which were either filled with cellular components (morphotype 1) or were completely empty (morphotype 3). Morphotype 4 showed strong green fluorescence indicative of high bacterial cell numbers (Fig. 3c and Supplementary Fig. 5). Since *B. rhizoxinica* accumulates in the hyphal tip prior to the formation of sporangia^8^, this morphotype was termed abortive sporangiophore.

**Fig. 3.**
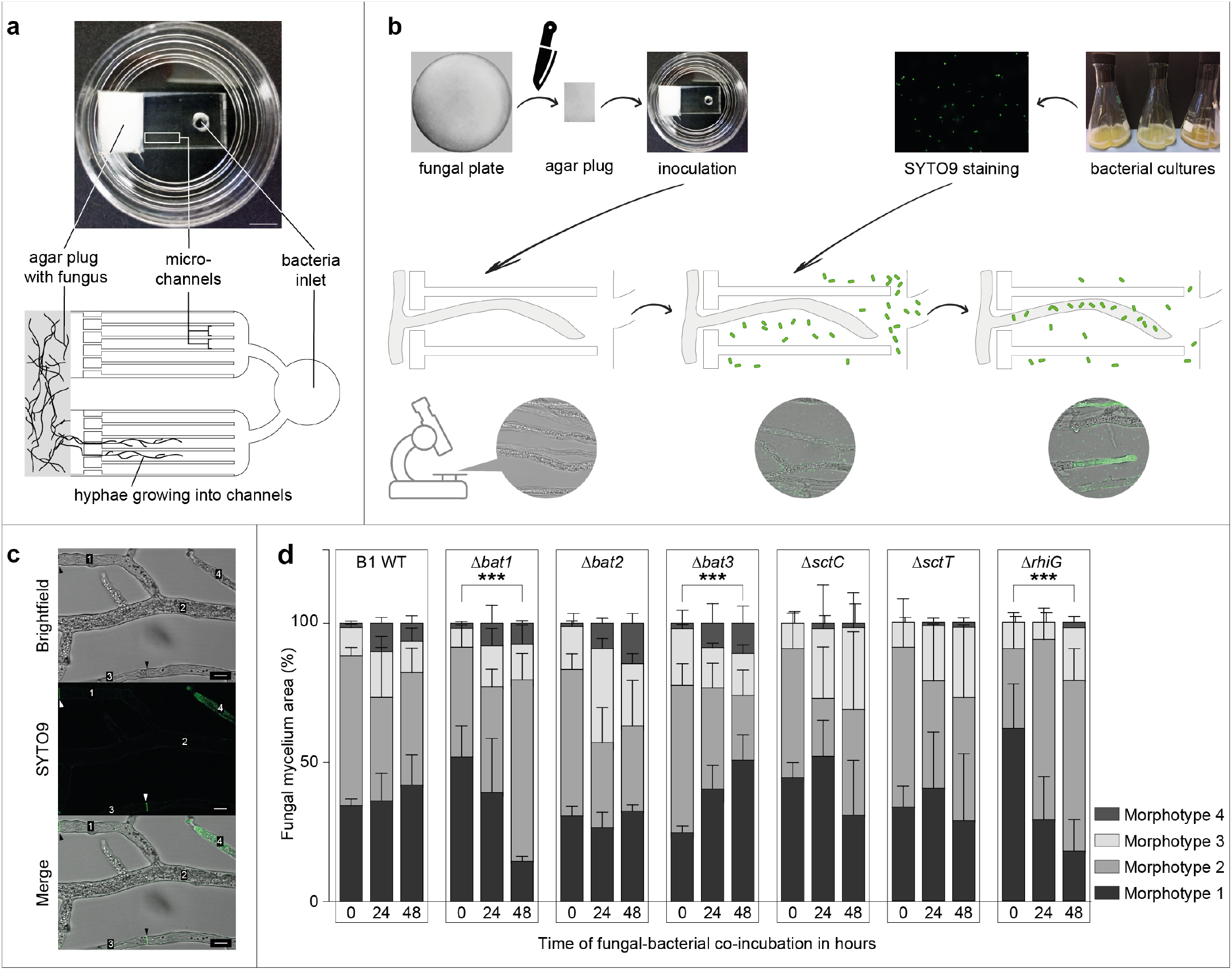
Phenotypic observations of *Rhizopus microsporus* co-cultivated with *Burkholderia rhizoxinica* wild-type or mutant strains in bacterial-fungal interaction (BFI) devices. **a**, Photograph and simplified illustration showing the microfluidic device in a glass petri dish. The BFI device is made of a patterned poly(dimethylsiloxane) layer bonded to glass-bottomed dish to form microchannels. The microchannels were filled with potato dextrose broth. Scale bar: 5 mm. A simplified two-dimensional representation of the design showing the narrow entry points into the microchannels which limits the number of hyphae that can enter the device^24^. **b**, Illustration showing the workflow for observation of BFIs. An agar plug containing endosymbiont-free *R. microsporus* is placed in direct contact with the microchannels. After two days of incubation, hyphae are growing inside the microchannels and SYTO9-stained bacterial strains are introduced into the microchannels via the ‘bacteria inlet’. Fungal reinfection is monitored over 48 hours and microscopic images are taken every 24 hours. **c**, Microscopic images of *R. microsporus* co-cultivated with wild-type *B. rhizoxinica* (stained with SYTO9) depicting four morphologically distinct types of hyphae (1: vegetative side hyphae; 2: vegetative main hyphae; 3: vegetative empty side hyphae; 4: abortive sporangiophores). Arrowheads indicate the presence of septa. Images were taken 48 hours post infection (hpi). Scale bar: 10 μm. Microscopic images of *R. microsporus* co-cultivated with *B. rhizoxinica* mutant strains are shown in Supplementary Fig. 5. **d**, The fungal mycelium area (in percent) of each morphotype was measured over a 48 hour time period of co-incubation in BFI devices. At time point 0, bacterial cells (B1WT: wild-type *B. rhizoxinica; Δbat1: Δbat1::Apra^r^*, Δ*bat2:* Δ*bat2*::Kan^r^, Δ*bat3:* Δ*bat3*::Kan^r^, Δ*sctC:* Δ*sctC*::Kan^r^, Δ*sctT:* Δ*sctT*::Kan^r^, and Δ*rhiG:* Δ*rhiG*::Kan^r^) were stained with SYTO9, added to the inlet, and co-incubated with endosymbiont-free *R. microsporus*. Images were taken at the time of infection (0 hpi), as well as 24 and 48 hpi. N = 3 biological replicates (16 technical replicates) ± one SEM. One-way ANOVA with Tukey’s multiple comparison test (*\*\*\*p<0.05*, significant differences are indicated for morphotypes 1 and 2, Supplementary Table 8).

Next, we calculated the mycelium area (as a percentage) of all four hyphal morphotypes over time (0, 24, and 48 hours post infection) using brightfield images (Fig. 3d). We monitored changes in the fungal morphological composition by: (i) comparing individual morphotypes at specific time points across *B. rhizoxinica* wild-type and mutant strains and; (ii) by comparing morphotypes of individual *B. rhizoxinica* strains between time points. In general, we observed no significant changes in any of the *B. rhizoxinica* strains, with two exceptions (Supplementary Table 8a,b). After 48 hours, morphotype 1 significantly decreased (*p<0.05*) and morphotype 2 significantly increased (*p<0.05*) in fungi co-cultured with BAT1 mutants, while the reverse effect was observed for fungi co-cultured with BAT3 mutants (Fig. 3d and Supplementary Table 8c). Interestingly, fungal reinfection with wild-type *B. rhizoxinica* and BAT-deficient mutants resulted in a noticeable increase in morphotype 4 after 48 hours. Although this higher percentage of morphotype 4 was not significant, this effect was virtually absent in the control strains (Fig. 3d).

### Monitoring fungal re-infection in real-time

The reinfection process was investigated in more detail by calculating the number of endohyphal bacteria from the integrated density at 485/498 nm (Fig. 4a). One day after infection, BAT1 and BAT3 mutants showed a 30% increase in fungal reinfection, compared to the positive control (sterile *R. microsporus* ATCC62417/S re-infected with wild-type *B. rhizoxinica*), while 25% less BAT2 mutant cells were detected inside the fungal hyphae. In a control experiment, endobacteria were virtually absent when T3SS defective mutants were co-cultured with the cured fungus (Supplementary Fig. 6). In addition, rhizoxin-deficient mutants were able to enter the fungal cells similarly to the positive control (Supplementary Fig. 7).

**Fig. 4.**
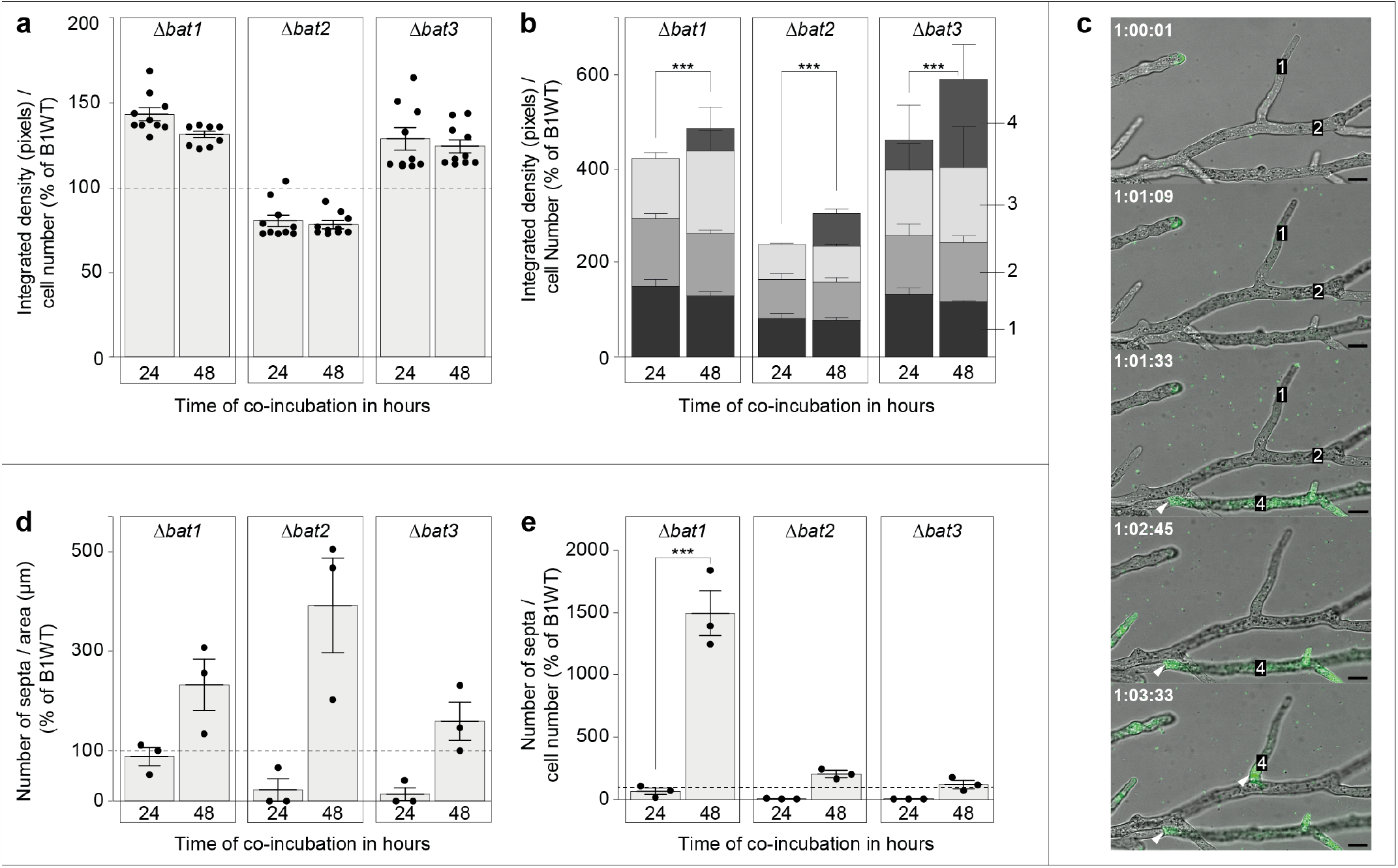
Course of reinfection of *Burkholderia rhizoxinica* transcription-activator like effector (BAT) mutant strain into *Rhizopus microsporus*. **a**, Endosymbiont-free *R. microsporus* ATCC62417/S was co-incubated with *B. rhizoxinica* BAT mutant strains (Δ*bat1*::Apra^r^, Δ*bat2*::Kan^r^, and Δ*bat3*::Kan^r^) for 48 hours. Bacterial cells were stained with SYTO9 prior to co-incubation. Following fluorescence microscopy at 485/498 nm (SYTO9), the integrated density (product of area and mean grey value) per bacterial cell number was calculated for both measurements using Fiji^36^, and then plotted as percent of the positive control (*R. microsporus* ATCC62417/S co-incubated with wild-type *B. rhizoxinica;* % of B1WT). N = 10 biological replicates ± one SEM. **b**, Integrated density (in % of B1WT) was measured for each individual morphotype (1 – 4) following reinfection with BAT mutant strains. N = 3 biological replicates (16 technical replicates) ± one SEM. One-way ANOVA with Tukey’s multiple comparison test (\*\*\**p<0.05*, significant differences are indicated for morphotype 4, Supplementary Table 9). **c**, Course of reinfection of *B. rhizoxinica Δbat1::Apra^r^* into *R. microsporus*. Reinfection was monitored over time using fluorescence microscopy (Video 2). Arrowheads indicate the formation of septa. Scale bar, 10 μm. **d,** The number of septa per area (in μm) was plotted as % of B1WT. N = 3 biological replicates (16 technical replicates) ± one SEM. **e,** The number of septa per bacterial cell number was plotted as % of B1WT. N = 3 biological replicates (16 technical replicates) ± one SEM. One-way ANOVA with Tukey’s multiple comparison test (\*\*\**p<0.05*, Supplementary Table 10).

The number of fluorescent BAT mutants inside the fungal hyphae after two days of co-incubation was similar to that seen one day after co-incubation (Fig. 4a and Supplementary Table 9), suggesting that reinfection with BAT mutants takes place within 24 hours of their introduction (Supplementary Fig. 5). It is possible that a potential increase in bacterial cells after 48 hours was not detected due to bleaching of SYTO9^35^. To minimise the effects of bleaching, light exposure was kept to a minimum by using a low exposure time (100 ms). Surprisingly, the four fungal morphotypes differed across the 48 hour infection period. The number of bacteria observed in morphotype 4 (abortive sporangiophore) after two days of coincubation was significantly higher (*p<0.05*) compared to one day after infection for all BAT mutants (Fig. 4b and Supplementary Table 9). Bacterial localisation in morphotype 4 was also observed in microscopic images (Fig. 4c and Supplementary Fig. 5).

**Fig. 5.**
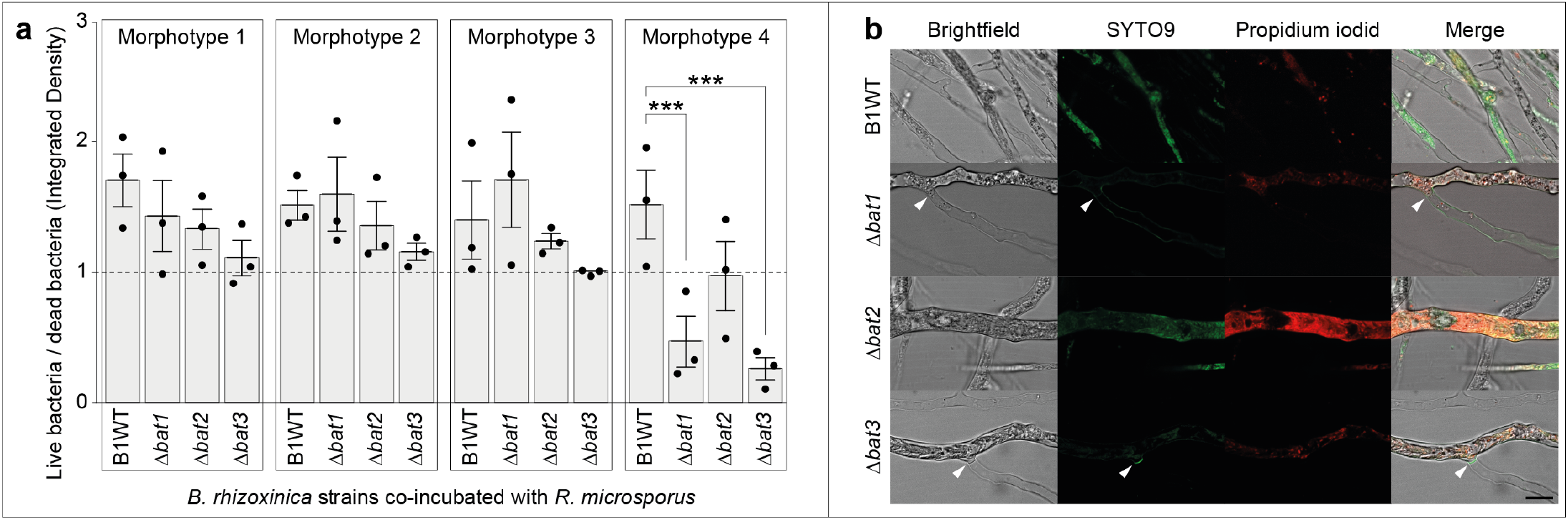
Viability test of *Rhizopus microsporus* re-infected with *Burkholderia rhizoxinica* transcription-activator like effector (BAT) mutant strains. **a**, Endosymbiont-free *R. microsporus* ATCC62417/S was co-incubated with *B. rhizoxinica* BAT mutant strains (Δ*bat1*::Apra^r^, Δ*bat2*::Kan_r_, and Δ*bat3*::Kan_r_) for 72 hours. Co-cultures were stained with LIVE/DEAD BacLight fluorescent dyes inside the microfluidic device. Following fluorescence microscopy, the integrated density was calculated for both live (SYTO9) and dead bacteria (propidium iodide) using Fiji and the ratio (live/dead) was plotted for each morphotype (see Methods for details). N = 3 biological replicates (16 technical replicates) ± one SEM. One-way ANOVA with Tukey’s multiple comparison test (\*\*\**p<0.05*, Supplementary Tables 10 and 11). b, Microscopic images of *R. microsporus* re-infected with *B. rhizoxinica* wild-type (B1WT) or BAT mutant strains stained with LIVE/DEAD BacLight fluorescent dyes. Arrowheads indicate the presence of septa. Scale bar, 20 μm.

The fungal reinfection and subsequent formation of morphotype 4 was captured in real-time (Fig. 4c and Video 2). First, BAT mutants penetrate and re-infect selected hyphae of *R. microsporus*, especially side-hyphae. Following infection, the bacterial cells reach very high densities (strong green-fluorescent signal) and septa are formed at the base of these hyphae.

As a result, the cytoplasmic flow between infected hyphae and the remaining fungal mycelium is abolished, leading to the physical containment of bacteria (Video 2). These observations were surprising, considering that *R. microsporus* generally does not form septa. In the rare cases where septa can be found, they are formed to either wall off old or injured hyphae or as separators between mycelia and spore-forming structures^33,37^.

### BAT mutants induce septation in fungal host

To further investigate the formation of septa, we analysed the total number of septa formed following fungal reinfection with *B. rhizoxinica* BAT mutants (Δ*bat1*::Apra^r^, Δ*bat2*::Kan^r^, and Δ*bat3*::Kan^r^) in a BFI device. The total number of septa was smaller or equal to the positive control (≤100% of sterile *R. microsporus* ATCC62417/S re-infected with wild-type *B. rhizoxinica*) 24 hours after reinfection, while septa formation increased markedly after 48 hours of co-incubation (Fig. 4d,e and Supplementary Table 10a). Compared to the reinfection process where most bacteria are inside the fungal hyphae after 24 hours (Fig. 4a), the formation of septa appears to take place at a later stage of the reinfection process (48 hours). In addition, septa formation and accumulation of bacteria in side-hyphae correlated with morphotype 4 (Fig. 4c). Using a LIVE/DEAD stain we observed that the majority of BAT-deficient mutants were dead in morphotype 4, while all other morphotypes contained more live bacteria (*p<0.05*). In contrast, wild-type *B. rhizoxinica* showed similar ratios of live versus dead bacteria in all four hyphal morphotypes (Fig. 5a,b and Supplementary Table 11).

It has to be taken into consideration that LIVE/DEAD staining, utilising SYTO9 and propidium iodide (PI), may underestimate bacterial viability. SYTO9 can stain live and dead cells with different efficiencies^35^, while PI has a strong binding affinity to extracellular nucleic acids, a major component of bacterial biofilms^38^. Since *B. rhizoxinica* is a known biofilm producer^32^, we performed LIVE/DEAD stains of axenic cultures prior to co-incubation in BFI devices. The majority of the cells stained green (Supplementary Fig. 8), indicating that underestimation of bacterial viability due to extracellular nucleic acids is unlikely.

## Discussion

Bacteria that live in close association with eukaryotes can control their host by employing specialised tools^39,40^. One prominent example are the TALEs, a T3SS-associated group of highly homologous proteins that allow plant pathogens to modify and exploit their hosts^27,30,41^. In recent years, TALEs have become popular tools in biotechnological applications due to their ability to manipulate DNA in a site-directed manner^42^. As a consequence, TALE-related proteins have been identified in a wide range of bacterial species, including *B. rhizoxinica* (BAT1, BAT2, and BAT3)^26^. In this study we combined genomic and functional studies to characterise T3SS-associated proteins in the fungal endosymbiont *B. rhizoxinica*. While fungal sporulation is independent of the T3SS-associated alanine-tryptophan-arginine (AWR) protein, targeted gene knockouts resulted in BAT-deficient mutants that are unable to induce fungal sporulation in the host, suggesting that BAT proteins are an important factor for the *Rhizopus* – *Burkholderia* interaction.

By utilising a tailor-made microfluidic device^24^, we show that the fungal mycelium is composed of four morphologically distinct types of hyphae. These fungal morphotypes are rare in the Zygomycetes^33^, and this is the first time that different cell types have been described in *R. microsporus*. The vegetative main hyphae of *R. microsporus* (morphotype 2), is one of the most important type of hyphae. These so called trunk hyphae^43^ transport intracellular material across the *R. microsporus* mycelium at relative high speeds. This allows *R. microsporus*, and fungi in general, to cope well with heterogeneous distributions of soil nutrients in their natural habitat^34^. In addition, fast cytoplasmic flow also allows for free migration of *B. rhizoxinica* across the host mycelium^44^. Hyphae serving as dispersal vectors for motile bacteria are an important feature of BFI ecology^45^ and may promote colonisation of new ecological niches^46^.

Microfluidics was used to gain a deeper understanding of the interaction between BAT-deficient endosymbionts and their fungal hosts. We have demonstrated (i) efficient fungal re-infection of BAT mutant strains within one day, and (ii) accumulation and trapping of these strains in specific fungal side-hyphae after two days. Accumulation of mutant endosymbionts is unique to BAT-deficient *B. rhizoxinica*, as T2SS- and T3SS-deficient mutants are unable to reinfect the sterile fungus efficiently^11,21^. Since total bacterial reinfection reaches its maximum after one day, it is conceivable that BAT-deficient bacteria are redistributed inside the fungal mycelium following reinfection and subsequently accumulate in side-hyphae. These infected side-hyphae are then walled-off from the remaining mycelium via the formation of septa, which leads to trapping of BAT-deficient bacteria. The occurrence of septa is one of the most surprising results, as Zygomycetes generally lack septate hyphae^33^. Septation causes the cytosolic exchange to stop and bacteria, which normally move freely within the mycelium^8^, become physically restricted.

Considering that the majority of trapped bacteria are dead, our observations are in agreement with a model where the lack of BAT proteins marks *B. rhizoxinica* as a pathogen or potentially harmful, thereby eliciting a protective response from the fungus. It should be kept in mind that these hyphae could represent abortive sporangia due to the set-up of the microfluidics devices. *R. microsporus* is grown submerged in liquid medium under which conditions the fungus struggles to produce sporangia. However, considering that the persistence of the symbiosis is dependent on spores containing endobacteria^8^, we would expect the majority of the trapped endobacteria to be alive (as seen for wild-type bacteria). Therefore, we argue that BAT-deficient bacteria induce a controlled suicide of the host, a mechanism that is well described in plantpathogen interactions. For example, TALEs from *Xanthomonas campestris* induce a suicide gene in resistant plants, causing programmed cell death of the host and thereby preventing the pathogen from spreading^47^.

Notably, BAT proteins described in this study, are the first TALE-related proteins whose absence causes an increase in host colonisation. In comparison, TALE proteins from plant pathogens aid host colonisation through the induction of susceptibility genes, which leads to a dramatic increase in bacterial cell numbers in the host^31^. It is remarkable that the absence of BAT proteins causes a similar phenotype in fungi, i.e. high numbers of bacterial cells. Thus, BATs might represent DNA-binding proteins with a function different to TALEs^42^. For example, multiple unrelated proteins showing nuclear localisation and interaction with DNA have been identified as mechanisms to aid host colonisation for various pathogens and symbionts^39^. The fact that parasites and mutualists commonly share the same molecular tools (e.g. secretion systems) to control their hosts^1^ supports the hypothesis that *B. rhizoxinica* becomes pathogenic ‘in the eyes of the fungus’ if BAT proteins are absent. Indeed, multiple investigations have suggested that *B. rhizoxinica* endosymbionts were initially parasites before switching to mutualists^48^. For example, phylogenetic analyses indicated that *B. rhizoxinica* has switched its host multiple times during evolution^16^, while *R. microsporus* has acquired resistance against a potent phytotoxin (rhizoxin) produced by the endosymbiont^49^. Thus, BAT proteins might represent an important factor for the maintenance of the mutualistic relationship in this BFI and their absence may facilitate a switch from mutualists back to parasites. This challenges the traditional views of symbiotic relationships, which were defined as a mutually beneficial for both partners. Instead, the BFI presented in this paper may be better described as a sliding scale, where symbionts can act as mutualist or pathogen depending on the genetic context^50^.

## Conclusion

In summary, we show that BAT-deficient endosymbionts successfully re-colonise the host but become trapped in specific hyphae due to the formation of septa. This represents an unprecedented case where mutant bacteria induce biogenesis of septa in a Zygomycete. In contrast to TALE proteins from phytopathogenic bacteria, a lack of BAT proteins facilitates bacterial colonisation of the host. Finally, microscopic investigations revealed that TALE-deficient mutants could be regarded as pathogens in the fungal cytosol, which would be in line with a plausible antagonism-mutualism shift in the evolution of this microbial symbiosis. The possibility of endobacteria controlling the physiology of a fungal host offers a broader view on the dynamic interactions between bacteria and fungi.

## Methods

### Strains and growth conditions

Eight *Rhizopus microsporus* strains harbouring endobacteria *Burkholderia* sp. were used in this study (Supplementary Table 1)^16^. Endobacteria from *R. microsporus* ATCC62417 were eliminated by continuous antibiotic treatment^8^ and the cured fungal strain was named ATCC62417/S. All *R. microsporus* strains (ATCC62417 and ATCC62417/S) were cultivated on Potato Dextrose Agar (PDA; Becton, Dickinson & Company, Sparks, MD, USA) at 30 °C. Bacterial endosymbionts were isolated from the mycelium of eight fungal strains as previously reported ^51^. Pure cultures of *B. rhizoxinica* were grown at 30 °C in MGY M9 medium (10 g/L glycerol, 1.25 g/L yeast extract, M9 salts) or Standard I Nutrient Agar (Merck, Darmstadt, Germany) supplemented with 1% glycerol.

### Genome sequencing, assembly, and annotation

Genomic DNA (gDNA) was isolated from seven axenic *Burkholderia* sp. cultures (Supplementary Table 1) using the Epicentre MasterPure DNA Purification kit (Illumina Biotechnology, San Diego, CA, USA) following the manufacturer’s protocol. gDNA was quantified using a NanoDrop™ (Thermo Fisher Scientific, Waltham, MA, USA) and used for shotgun sequencing. Sequencing was performed at the University of Melbourne using the Nextera XT Library preparation kit in combination with the Illumina™ NextSeq System^52^ producing paired-end reads with a mean length of approximately 500 base pairs.

Raw sequence reads were trimmed with trimmomatic^53^ and reference assemblies were performed using Spades v3.7.0, with the previously published *B. rhizoxinica* HKI-0454 genome^14^ as a reference. Draft genome sequences were error corrected with Lighter^54^ and annotated using Prokka version 1.12^55^. The RNA genes were determined by using the software Barnap implemented in Prokka. The key attributes for the genome sequences are summarized in Supplementary Table 2.

### *In silico* predictions and characterisation of Type 3 effectors

Potential type 3 secreted effector proteins were predicted using the T3SS PREDICTION server^56^ and the EFFECTIVE T3 prediction tool^57^. Nuclear localisation signals (NLS) were predicted using the cNLS domain prediction tool^58^ and NucPred^59^. The nucleotide sequences of potential *Burkholderia* T3 effector genes have been deposited in GenBank under the accession numbers provided in Supplementary Table 3.

### Amplification and Sanger Sequencing of *Burkholderia* sp. *bat* genes

gDNA was isolated from eight axenic *Burkholderia* spp cultures (Supplementary Table 1) and quantified as described above. Polymerase chain reaction (PCR) primers were designed to amplify partial coding sequences of *bat1* (GenBank acc. no.: RBRH_01844, Supplementary Table 2a). PCRs were performed in 25.0 μL reactions containing: 12.5 μL of high-fidelity Taq DNA polymerase (Phusion^®^ Master Mix, New England Biolabs, Ipswich, MA, USA), forward and reverse primers (both 0.4 μM), and 100 ng of template gDNA. The following thermocycling conditions were used for amplification: 98 °C / 30 s, 1 cycle; 98 °C / 10 s, 65 °C / 30 s, 72 °C / 3 min, 30 cycles; 72 °C / 7 min, 1 cycle; 16 °C / hold.

The PCR products were checked on an agarose gel stained with ethidium bromide before gel extraction (Zymoclean™ Gel DNA Recovery Kit, Zymo Research, Irvine, CA, USA). The resulting pure amplicons were ligated into pCR™-Blunt II-TOPO^®^ (Invitrogen, Carlsbad, CA, USA), followed by transformation into TOP10 *Escherichia coli*. The plasmids were purified (Monarch Plasmid Miniprep Kit, New England Biolabs) and plasmid inserts were bidirectionally sequenced by an external contractor (Eurofins Genomics, Ebersberg, Germany). Sequences were deposited in GenBank under the accession numbers provided in Supplementary Table 1.

### Phylogenetic analysis

For phylogenetic analysis, predicted T3 effector protein sequences were aligned using ClustalW^60^. Alignments were generated using a gap open penalty of 10 and a gap extension penalty of 0.1 as implemented in the MEGA7 package (Molecular Evolutionary Genetics Analysis software, version 5.0)^61^. All positions containing gaps and missing data were eliminated. The evolutionary history was inferred using the Neighbour-Joining method with maximum composite likelihood distances and 10,000 bootstrap repetitions^62,63^. The alignments of sequences used in this study are shown in Supplementary Figs. 1 and 3.

### Generation of *B. rhizoxinica* mutant strains

To investigate the role of both AWR and BAT proteins in the symbiosis, four genes (*awr*: RBRH_03012; *bat1:* RBRH_01844; *bat2:* RBRH_01776; *bat3*: RBRH_01777) were deleted using a double crossover strategy as previously described^21^.

Using a proof-reading polymerase, the upstream and downstream regions of the genes of interest were amplified. Primers were designed to contain 20 bp overlap with the gene of interest and as well as a 20 bp overlap with an antibiotic resistance cassette (*bat2*, *bat3:* kanamycin, *bat1:* apramycin). The kanamycin and apramycin cassettes were amplified from pK19 and pJI773, respectively, using primers carrying the same 20 bp.

The knockout vector pGL42a was used to generate *B. rhizoxinica Δawr::Kan^r^ Δbat1::Apra^r^* Δ*bat2*::Kan^r^, and Δ*bat3*::Kan^r^ deletion mutants. Additionally, a double mutant (Δ*bat2_bat3::Kan^r^*) was generated by using the upstream region of *bat3* and the downstream region of *bat2*. The plasmid pZU52 (Δ*bat2_bat3*::Kan^r^) was introduced into *B. rhizoxinica* together with plasmid pZU17 (Δ*bat1*::Apra^r^) to generate a triple mutant (Δ*bat1::*Apra^r^*-* Δ*bat2_bat3:*:Kan^r^). pGL42a was double-digested with the restriction enzymes SpeI and KpnI (New England Bioloabs). The linear vector was gel-purified (Monarch^®^ DNA Gel Extraction Kit, New England Bioloabs) and quantified on a NanoDrop™ (Thermo Fisher Scientific).

For each target gene, equimolar amounts of three PCR products and linear pGL42a were mixed with NEBuilder^®^ 2X Master Mix (NEBuilder^®^ HiFi DNA Assembly Cloning Kit, New England Bioloabs) and incubated at 60 °C for 1 hr following the manufacturer’s recommendations. The new plasmids pZU52 (Δ*AWR*), pZU17 (Δ*bat1*), pZU21 (Δ*bat2*), pZU19 (Δ*bat3*), pZU23 (Δ*bat2_bat3*), and pZU17/pZU23 (Δ*bat1-Δbat2_bat3*) were introduced into *E. coli* by chemical transformation. Transformants were selected on standard nutrient agar supplemented with either 50 μg/mL kanamycin (pZU52, pZU21, pZU19, pZU23) or 50 μg/mL apramycin (pZU17).

Competent *B. rhizoxinica* HKI-454 (B1 WT) cells were transformed with vectors pZU52, pZU17, pZU21, and pZU19 via electroporation. Transformants were grown on standard nutrient agar containing 50 μg/mL of either apramycin or kanamycin. Colonies were subsequently passaged onto agar plates containing double selection medium^21^ until the correct knockout constructs were observed using colony PCR. Colony PCRs were carried out in 12.0 μL final volumes containing: 5 μL of high-fidelity OneTaq^®^ QuickLoad^®^ 2X Master Mix (New England Biolabs), appropriate forward and reverse control primers (both 0.4 μM, Supplementary Table 14), and 5 μL colony suspension. The following thermocycling conditions were used for amplification: 96 °C / 3 min, 1 cycle; 96 °C / 10 s, 58 °C / 15 s, 68 °C / 1 min, 30 cycles; 68 °C / 5 min, 1 cycle; 16 °C / hold. The resulting PCR products were visualised on an agarose gel. Primers were designed to span the two recombination sites, yielding amplicons A and B in mutant strains and amplicons C and D in *B. rhizoxinica* wild-type strains.

### Sporulation assay

In a liquid sporulation assay, sterile *R. microsporus* aerial hypha (app. 0.1 cm^3^) was grown in 24-well plates containing 750 μL VK medium (5 g/L glycerol, 10 g/L yeast extract, 10 g/L corn starch, 10 g/L corn step solids, 10 g/L CaCO3; pH 6,5). After 15 hours of incubation, 100 μL of overnight wild-type or mutant *B. rhizoxinica* cultures were added to individual wells in three biological replicates. Co-culture plates were inspected daily for sporulation. The spores were harvested from each well using 500 μL NaCl (0.15 M) and counted using a Thoma Chamber.

Experiments were performed three times independently (N = 3) with three technical replicates on each plate. Data are presented as means with SD. Raw data from sporulation experiments were processed with MS Excel. GraphPad Prism 5.03 (GraphPad Software, La Jolla, California, USA, www.graphpad.com) was used for statistical analysis and graphing. Data from spore counts were compared for *B. rhizoxinica* wild-type and *B. rhizoxinica* mutant strains using one-way analysis of variance (ANOVA) and Tukey HSD test function in GraphPad. P-values with *p<0.05* were considered statistically significant. The Brown-Forsythe test was used to test for equal variance and a *p* value with < *0.05* was considered significant.

### Fluorescence microscopy of co-cultures

One-week old co-culture fungal hyphae were used to visualise the localisation of the *B. rhizoxinica* mutant strains. The bacterial cells were stained with 5 μM Syto 9 (Invitrogen) and fungal cells were counter-stained with 2 μg/mL calcofluor white (Fluka, Germany) for 5-10 min. Fluorescent microscopy was carried out using a Zeiss LSM 710 confocal laser-scanning microscope, and images were captured using the Zeiss-Zen software.

### Inoculation of bacterial-fungal interaction devices with fungus

Bacterial-fungal interaction (BFI) devices were prepared as previously described^24^. Before the BFI devices were inoculated, cured (endosymbiont-free) *R. microsporus* ATCC62417/S was subcultured on PDA plates for two days at 30 °C. A piece of young, growing mycelium (app. 1 cm^3^) was cut from the agar plate and placed upside down in front of the microchannels of a BFI device filled with Potato Dextrose Broth (PDB; Becton, Dickinson & Company). The agar plug was positioned with the growth direction of the fungus facing the microchannels. We paid special attention to the size of the agar plug to minimise biases between experiments. After two days of incubation at 30 °C, the hyphae had grown far enough into the microchannels to perform subsequent reinfection experiments.

### Bacterial inoculation into bacterial-fungal interaction devices

Bacterial over-night cultures (500 μL) were harvested in an Eppendorf tube and resuspended in 0.5 mL NaCl (0.85%) containing SYTO9 green-fluorescent nucleic acid stain (5 nM final concentration, Invitrogen). Following incubation in the dark for 5 min, stained cells were washed and resuspended in 0.5 mL PDB medium. Cells were counted using a CASY Cell Counter (OMNI Life Science, Bremen, Germany) and then added to the bacteria inlet of the BFI device (app. 50–100 μL), which is connected with the microchannels containing the cured fungus (Fig. 3a,b).

### Live-cell imaging of bacterial-fungal interactions

A fluorescence spinning disc microscope (Axio Observer microscope-platform equipped with Cell Observer SD, Zeiss) was used to capture BFIs at 0 h, 24 h, and 48 h after bacterial inoculation. For each time point, 16 images were taken at random position of the microchannels, including both ends of the BFI device (N = 16 biological replicates). Brightfield images were captured using a laser intensity of 7.1 V and an exposure time of 100 ms. Bacterial cells (wild-type *B. rhizoxinica*, Δ*bat1*, Δ*bat2*, Δ*bat3*, Δ*sctT*, Δ*sctC, and* Δ*rhiG*), stained with SYTO9, were captured at 485/498 nm with an exposure time of 100 ms. As a reference, wild-type *R. microsporus* ATCC62417, naturally containing endosymbionts, was also analysed, although no additional bacterial cultures were added through the inlet. Each reinfection experiment was performed at least three times independently (N ≥ 3). Images were generated using custom software (Zeiss Zen Blue Software) and analysed using Fiji^36^. Images, containing spatial calibration meta data, were imported to Fiji and the same spatial scale was applied to all images using the global scale function. Prior to image analyses, all images were converted to 16-bit greyscale, which was propagated to both channels (brightfield and 485/498 nm). The hyphal area and integrated density (ID; product of area and mean grey value) of both channels were measured using the free hand tool and measuring tool implemented in Fiji. According to the brightfield images, four morphologically distinct types of hyphae were defined (Fig. 3d) and the number of septa observed was counted. Subsequent calculations and data visualisation was carried out using MS Excel and GraphPad Prism 5.03. To correct for autofluorescence and variations between experiments, raw ID values at 0 hours post infection (hpi) were averaged from three independent experiments and then subtracted from raw ID values at 24 hpi and 48 hpi. Corrected ID values were divided by the number of bacterial cells (cells/mL) inoculated in the bacterial inlet and then plotted as percent of the positive control (cured *R. microsporus* ATCC62417/S co-incubated with wild-type *B. rhizoxinica* HKI-0454). The same approach was applied to correct for the number of septa observed. The corrected number of septa was divided by either the inoculated bacterial cell number (cells/mL) or by the mycelium area (in μm) and then plotted as percent of the positive control.

### Imaging of bacterial survival

To test the bacterial survival rate in BFI devices, cells were stained with LIVE/DEAD BacLight fluorescent dyes (Invitrogen) after 72 h of fungal-bacterial co-incubation. To stain cells inside the BFI device, the PDB medium was removed through the bacterial inlet and replaced with 100 μL staining solution (10 nM SYTO9 and 60 nM propidium iodide). Fluid replacement was repeated three times, before the devices were incubated in the dark for 15 min. Following staining, fluorescent dyes were replaced with PDB medium by flushing the devices four times. Fluorescence microscopy and subsequent image analysis was performed as described above.

### Statistical analysis

Raw data from re-infection experiments and bacterial survival experiments were processed with MS Excel. GraphPad Prism 5.03 was used for statistical analysis and graphing. One-way analysis of variance (ANOVA) was used to study the relationship between different *B. rhizoxinica* strains (wild-type and mutant strains) and fungal physiology (e.g. mycelium area, reinfection efficiency, septation rate, bacterial survival) following fungal reinfection using the Tukey HSD test function. P-values with *p<0.05* were considered statistically significant. The Brown-Forsythe test was used to test for equal variance and a *p* value with < *0.05* was considered significant.

## Data availability

All data generated or analysed during this study are included in the manuscript and in the supporting files. Source data files have been provided for Supplemental videos.

## Acknowledgements

We thank S. Lindner for assistance with image analyses. I.R. is grateful for financial support from the European Research Council (ERC) for a Marie Sklodowska-Curie Individual Fellowship (MSCA-IF-EF-RI) Project reference 794343. Financial support by the Deutsche Forschungsgemeinschaft (SFB 1127 ChemBioSys, and Leibniz Award to C.H.), the JSMC to Z.U., and the Swiss National Science Foundation in the form of an Ambizione Career Grant (PZ00P2_168005) to C.E.S. is gratefully acknowledged.

## Author contributions

IR, ZU, NM, TPS, SJP, IF, Acquisition of data, Analysis and interpretation of data, Drafting or revising the article; CES, Preparation of microfluidic devices, Drafting or revising the article; FH, Analysis and interpretation of data, revising the article; CH, Conception and design, drafting and revising the article.

## Competing interests

The authors declare no competing interests.

## Note

During the submission process we have noted the publication of a pre-print manuscript with complementary results to our study^64^.

