## Supplemental Material for "Secreted TAL effectors protect symbiotic bacteria from entrapment within fungal hyphae"

**Supplementary Information**

**Supplementary Table 1.** Fungal strains harbouring bacterial endosymbionts[1].

| **Taxon** | **Strain designation** | **Origin** | **Bacterial endosymbiont (isolate)** |
|---|---|---|---|
| *Rhizopus microsporus* van Tieghem | ATCC 62417 | Rice seedlings, Japan | *Burkholderia rhizoxinica* HKI-0454 (B1) |
| *Rhizopus* sp. strain F-1360 | ATCC 20577 | Soil, Japan | *Burkholderia* sp. strain HKI-0512 (B2) |
| *Rhizopus microsporus* Tieghem var. *microsporus* | CBS 111563 | Sufu starter culture, rice wine tablet, Vietnam | *Burkholderia* sp. strain HKI-0455 (B3) |
| *Rhizopus microsporus* Tieghem var. *microsporus* | CBS 699.68 | Soil, Ukraine | *Burkholderia* sp. strain HKI-0402 (B4) |
| *Rhizopus microsporus* Tieghem | CBS 112285 | Ground nuts, Mozambique | *Burkholderia endofungorum* HKI-0456 (B5) |
| *Rhizopus microsporus* var. *chinensis* (Saito) Schipper & Stalpers | CBS 261.28 | Not specified, USA | *Burkholderia* sp. strain HKI-0513 (B6) |
| *Rhizopus microsporus* Tieghem var. *microsporus* | CBS 700.68 | Forest soil, Georgia | *Burkholderia* sp. strain HKI-0403 (B7) |
| *Rhizopus microsporus* Tieghem var. *microsporus* | CBS 308.87 | Man, from deep necrotic tissue within the hand following a spider bite, Australia | *Burkholderia* sp. strain HKI-0404 (B8) |

**Supplementary Table 2.** Genome statistics of the seven *Burkholderia* strains sequenced in this study and the previously sequenced *B. rhizoxinica* HKI-0454 (GenBank acc. no.: GCA_000198775.1).

| **Strain** | **Genome size (bp)** | **GC-content (%)** | **Protein coding genes** | **Percent coding (%)** | **Hypothetical protein coding genes** | **tRNA genes** |
|---|---|---|---|---|---|---|
| *Burkholderia rhizoxinica* HKI-0454 (B1) | 3,750,140 | 60.69 | 2970 | 84.1 | 833 | 47 |
| *Burkholderia* sp. strain HKI-0512 (B2) | 3,795,941 | 60.62 | 3369 | 81.1 | 1146 | 45 |
| *Burkholderia* sp. strain HKI-0455 (B3) | 3,548,451 | 60.69 | 3103 | 81.5 | 965 | 47 |
| *Burkholderia* sp. strain HKI-0402 (B4) | 3,500,892 | 60.64 | 3035 | 82.1 | 907 | 46 |
| *Burkholderia endofungorum* HKI-0456 (B5) | 3,355,426 | 61.25 | 2967 | 82.3 | 920 | 49 |
| *Burkholderia* sp. strain HKI-0513 (B6) | 3,720,001 | 60.80 | 3254 | 79.6 | 1085 | 45 |
| *Burkholderia* sp. strain HKI-0403 (B7) | 3,591,347 | 60.55 | 3247 | 82.1 | 1051 | 47 |
| *Burkholderia* sp. strain HKI-0404 (B8) | 3,281,183 | 61.16 | 2884 | 83.7 | 837 | 48 |

**Supplementary Table 3.** Predicted nuclear localisation sequences (NLS) within AWR proteins from endofungal and plant-associated *Burkholderia* species, and plant pathogenic *Ralstonia solanaceraum* using **a,** NucPred prediction software[2] and **b,** cNLS Mapper[3]. GenBank accession numbers are given in brackets.

| a | Species (Gen. Bank Acc. No) | NLS motif (monopartite) | Position in protein[1] | NucPred Score[2] | Specificity[3] | Sensitivity (coverage)[4] |
|---|---|---|---|---|---|---|
| | *Ralstonia solanacearum* (BAH47286.1) AWR1 | GATRRRRNSG | 38 – 45 | 0.75 | 0.81 | 0.44 |
| | *Paraburkholderia graminis* (WP_006047106.1) | TRRRM | 36 – 40 | 0.62 | 0.71 | 0.53 |
| | *Burkholderia glumae* (WP_012733093.1) | QARRRQG | 78 – 84 | 0.55 | 0.70 | 0.62 |
| | *Burkholderia* sp. CCGE1003 (ADN59567.1) | DPQRRRRFVDD | 693 – 693 | 0.62 | 0.71 | 0.53 |
| | *Burkholderia rhizoxinica* HKI-0454 (B1) | KRSRKRKATV | 577 – 586 | 0.68 | 0.71 | 0.53 |
| | *Burkholderia* sp. strain HKI-0512 (B2) | KRSRKRKATV | 577 – 586 | 0.68 | 0.71 | 0.53 |
| | *Burkholderia* sp. strain HKI-0455 (B3) | KRSRKRKATV | 572 – 581 | 0.75 | 0.81 | 0.44 |
| | *Burkholderia* sp. strain HKI-0402 (B4) | KRSRKRKATV | 572 – 581 | 0.75 | 0.81 | 0.44 |
| | *Burkholderia endofungorum* HKI-0456 (B5) | KRSRKRKATV | 572 – 581 | 0.68 | 0.71 | 0.53 |
| | *Burkholderia* sp. strain HKI-0513 (B6) | KRSRKRKATV | 577 – 586 | 0.69 | 0.71 | 0.53 |
| | *Burkholderia* sp. strain HKI-0403 (B7) | KRSRKRKATV | 572 – 581 | 0.73 | 0.81 | 0.44 |
| | *Burkholderia* sp. strain HKI-0404 (B8) | KRSRERKATV | 559 – 568 | 0.62 | 0.71 | 0.53 |

| b | Species (Gen. Bank Acc. No) | NLS motif (monopartite) | Position in protein[1] | cNLS Score[5] | NLS motif (bipartite) | Position in protein[1] | cNLS Score[5] |
|---|---|---|---|---|---|---|---|
| | *Ralstonia solanacearum* (BAH47286.1) AWR1 | N/A | N/A | N/A | N/A | N/A | N/A |
| | *Paraburkholderia graminis* (WP_006047106.1) | N/A | N/A | N/A | N/A | N/A | N/A |
| | *Burkholderia glumae* (WP_012733093.1) | N/A | N/A | N/A | N/A | N/A | N/A |
| | *Burkholderia* sp. CCGE1003 (ADN59567.1) | N/A | N/A | N/A | N/A | N/A | N/A |
| | *Burkholderia rhizoxinica* HKI-0454 (B1) | RSRKRKATVQ | 578 – 587 | 13 | KATKATKATKLPKLPKLPKLPKLPKLP | 1083 – 1096 | 5.5 |
| | | | | | KATKATKATKLPKLPKLPKLPKLPKLPKLP | 1083 – 1112 | 6.1 |
| | | | | | KATKATKLPKLPKLPKLPKLPKLPKLPKG | 1086 – 1114 | 5.6 |
| | *Burkholderia* sp. strain HKI-0512 (B2) | RSRKRKATVQ | 578 – 587 | 13 | FMGKPVAATKATKLPKLPKLPKLPKLP | 1061 – 1090 | 5.5 |
| | | | | | KATKLPKLPKLPKLPKLPKLPKLPKG | 1070 – 1095 | 5.5 |
| | *Burkholderia* sp. strain HKI-0455 (B3) | RSRKRKATVQ | 573 – 582 | 13 | N/A | N/A | N/A |
| | *Burkholderia* sp. strain HKI-0402 (B4) | RSRKRKATVQ | 573 – 582 | 13 | N/A | N/A | N/A |
| | *Burkholderia endofungorum* HKI-0456 (B5) | RSRKRKATVQ | 573 – 582 | 13 | N/A | N/A | N/A |
| | *Burkholderia* sp. strain HKI-0513 (B6) | RSRKRKATVQ | 578 – 587 | 13 | KATKATKATKLPKLPKLPKLPKLPKLP | 1070 – 1096 | 5.5 |
| | | | | | KATKATKATKLPKLPKLPKLPKLPKLPKG | 1070 – 1098 | 5.5 |
| | *Burkholderia* sp. strain HKI-0403 (B7) | RSRKRKATVQ | 573 – 582 | 13 | N/A | N/A | N/A |
| | *Burkholderia* sp. strain HKI-0404 (B8) | N/A | N/A | | N/A | N/A | N/A |

[1] Amino acid position of predicted NLS within AWR peptide sequence.

[2] Fraction of proteins predicted to be nuclear that actually are nuclear.

[3] Fraction of true nuclear proteins that are predicted (coverage).

[4] cNLS score: > 10: protein exclusively localised to the nucleus; 7 – 8: partially localised to the nucleus; 3 – 5: localised to both the nucleus and the cytoplasm; 1 – 2: localised to the cytoplasm.

N/A: not available.

**Supplementary Table 4. a,** Approximate probabilities (p) of Brown-Forsythe test, **b,** one-way analysis of variance (ANOVA), **c,** and Tukey HSD Post Hoc Test for fungal spore counts following co-cultivation of *Rhizopus microsporus* with *Burkholderia rhizoxinica* wild-type (B1 WT), *B. rhizoxinica* transcription-activator like effector (BAT) mutant strains (Δ*bat1*::Apra$^r$, Δ*bat2*::Kan$^r$, Δ*bat3*::Kan$^r$, Δ*bat2_bat3*::Kan$^r$, and Δ*bat1*::Apra$^r$-Δ*bat2_bat3*::Kan$^r$), type 2 secretion system mutant strains (Δ*sctC*::Kan$^r$ and Δ*sctT*::Kan$^r$) or T3SS-associated alanine-tryptophan-arginine (AWR) mutant strains (Δ*awr*::Kan$^r$). Homogeneous data (non-significant Brown-Forsythe) is shown in black numbers and non-homogeneous data (significant Brown-Forsythe) is highlighted in red numbers. P-values with *p°<°0.05* were considered statistically significant (highlighted in grey).

**a**

| **Brown-Forsythe Test** | |
|---|---|
| F (DFn, DFd) | 2.170 (8, 18) |
| P value | 0.0821 |
| P value summary | ns |
| Are SDs significantly different (P < 0.05)? | No |

**b**

| **ANOVA Table** | **SS** | **DF** | **MS** | **F (DFn, DFd)** | **P value** |
|---|---|---|---|---|---|
| Treatment (between columns) | 283210 | 8 | 35401 | F (8, 18) = 23.30 | P<0.0001 |
| Residual (within columns) | 27347 | 18 | 1519 | | |
| Total | 310556 | 26 | | | |

**c**

| **Strain Comparison** | | **Mean Diff.** | **95% CI** | **P<0.05?** | **Summary** |
|---|---|---|---|---|---|
| B1WT vs. | Δ*bat1*::Apra$^r$ | 243 | 131.5 to 354.5 | Yes | *** |
| | Δ*bat2*::Kan$^r$ | 222 | 110.5 to 333.5 | Yes | *** |
| | Δ*bat3*::Kan$^r$ | 238.3 | 126.8 to 349.9 | Yes | *** |
| | Δ*bat2_bat3*::Kan$^r$ | 246.3 | 134.8 to 357.9 | Yes | *** |
| | *Δ*bat1*::Apra$^r$-* Δbat2_bat3*::Kan$^r$* | 207.3 | 95.80 to 318.9 | Yes | *** |
| | Δ*awr*::Kan$^r$ | −16 | −127.5 to 95.53 | No | ns |
| | Δ*sctC*::Kan$^r$ | 248.3 | 136.8 to 359.9 | Yes | *** |
| | Δ*sctT*::Kan$^r$ | 248.3 | 136.8 to 359.9 | Yes | *** |
| Δ*bat1*::Apra$^r$ vs | Δ*bat2*::Kan$^r$ | −21 | −132.5 to 90.53 | No | ns |
| | Δ*bat3*::Kan$^r$ | −4.667 | −116.2 to 106.9 | No | ns |
| | Δ*bat2_bat3*::Kan$^r$ | 3.333 | −108.2 to 114.9 | No | ns |
| | *Δ*bat1*::Apra$^r$-* Δbat2_bat3*::Kan$^r$* | −35.67 | −147.2 to 75.86 | No | ns |
| | Δ*awr*::Kan$^r$ | −259 | −370.5 to −147.5 | Yes | *** |
| | Δ*sctC*::Kan$^r$ | 5.333 | −106.2 to 116.9 | No | ns |
| | Δ*sctT*::Kan$^r$ | 5.333 | −106.2 to 116.9 | No | ns |
| Δ*bat2*::Kan$^r$ vs | Δ*bat3*::Kan$^r$ | 16.33 | −95.20 to 127.9 | No | ns |
| | Δ*bat2_bat3*::Kan$^r$ | 24.33 | −87.20 to 135.9 | No | ns |
| | *Δ*bat1*::Apra$^r$-* Δbat2_bat3*::Kan$^r$* | −14.67 | −126.2 to 96.86 | No | ns |

| | | | | | |
|---|---|---|---|---|---|
| | Δ*awr*::Kan$^r$ | −238 | −349.5 to −126.5 | Yes | *** |
| | Δ*sctC*::Kan$^r$ | 26.33 | −85.20 to 137.9 | No | ns |
| | Δ*sctT*::Kan$^r$ | 26.33 | −85.20 to 137.9 | No | ns |
| Δ*bat3*::Kan$^r$ vs | Δ*bat2_bat3*::Kan$^r$ | 8 | −103.5 to 119.5 | No | ns |
| | *Δ*bat1*::Apra*$^r$- *Δ*bat2_bat3*::Kan*$^r$ | −31 | −142.5 to 80.53 | No | ns |
| | Δ*awr*::Kan$^r$ | −254.3 | −365.9 to −142.8 | Yes | *** |
| | Δ*sctC*::Kan$^r$ | 10 | −101.5 to 121.5 | No | ns |
| | Δ*sctT*::Kan$^r$ | 10 | −101.5 to 121.5 | No | ns |
| Δ*bat2_bat3*::Kan$^r$ vs | *Δ*bat1*::Apra*$^r$- *Δ*bat2_bat3*::Kan*$^r$ | −39 | −150.5 to 72.53 | No | ns |
| | Δ*awr*::Kan$^r$ | −262.3 | −373.9 to −150.8 | Yes | *** |
| | Δ*sctC*::Kan$^r$ | 2 | −109.5 to 113.5 | No | ns |
| | Δ*sctT*::Kan$^r$ | 2 | −109.5 to 113.5 | No | ns |
| *Δ*bat1*::Apra*$^r$- *Δ*bat2_bat3*::Kan*$^r$ vs | Δ*awr*::Kan$^r$ | −223.3 | −334.9 to −111.8 | Yes | *** |
| | Δ*sctC*::Kan$^r$ | 41 | −70.53 to 152.5 | No | ns |
| | Δ*sctT*::Kan$^r$ | 41 | −70.53 to 152.5 | No | ns |
| Δ*awr*::Kan$^r$ vs | Δ*sctC*::Kan$^r$ | 264.3 | 152.8 to 375.9 | Yes | *** |
| | Δ*sctT*::Kan$^r$ | 264.3 | 152.8 to 375.9 | Yes | *** |
| Δ*sctC*::Kan$^r$ | Δ*sctT*::Kan$^r$ | 0 | −111.5 to 111.5 | No | ns |

Abbreviations: SS: sum of squares, df: degrees of freedom, MS: mean square, mean Diff.: mean difference, 95% CI: 95% confidence interval, ns: not significant, * $p°<°0.0332$, ** $p°<°0.0021$, *** $p°<°0.0002$.

**Supplementary Table 5.** Primers used for the amplification of *Burkholderia rhizoxinica* transcription activator-like effector 1 (*bat1*), 2 (*bat2*), and 3 (*bat3*) partial coding sequences in polymerase chain reactions (PCRs).

| Gene | GenBank acc. no. | Primer Sequence (5' - 3') | | Predicted amplicon size (bp) |
|---|---|---|---|---|
| *bat1* | RBRH_01844 | GCTACCACCTTGGCGTTTAT | Forward | 3100 bp |
| | | TACTTTGCGCGTAACAGCAC | Reverse | |

**Supplementary Table 6.** GenBank accession numbers of genes identified and used in this study.

| Gene | Locus Tag | Strain | GenBank Acc. No. |
|---|---|---|---|
| *awr* | RBRH_02721 | *Burkholderia rhizoxinica* HKI-0512 (B2) | MN840555 |
| *awr* | RBRH_00074 | *Burkholderia rhizoxinica* HKI-0455 (B3) | MN840550 |
| *awr* | RBRH_00457 | *Burkholderia rhizoxinica* HKI-0402 (B4) | MN840552 |
| *awr* | RBRH_01365 | *Burkholderia endofungorum* HKI-0456 (B5) | MN840553 |
| *awr* | RBRH_02790 | *Burkholderia rhizoxinica* HKI-0513 (B6) | MN840556 |
| *awr* | RBRH_03309 | *Burkholderia rhizoxinica* HKI-0403 (B7) | MN840551 |
| *awr* | RBRH_02340 | *Burkholderia rhizoxinica* HKI-0404 (B8) | MN840554 |
| *bat1* | RBRH_01844* | *Burkholderia* sp. HKI-0512 (B2) | MN891944 |
| *bat1* | RBRH_01585 | *Burkholderia rhizoxinica* HKI-0402 (B4) | MN840538 |
| *bat1* | RBRH_01844* | *Burkholderia endofungorum* HKI-0456 (B5) | MN891945 |
| *bat1* | RBRH_01580 | *Burkholderia rhizoxinica* HKI-0513 (B6) | MN840537 |
| *bat1* | RBRH_01934 | *Burkholderia rhizoxinica* HKI-0403 (B7) | MN840539 |
| *bat1* | RBRH_02907 | *Burkholderia rhizoxinica* HKI-0403 (B7) | MN840540 |
| *bat1* | RBRH_00422 | *Burkholderia rhizoxinica* HKI-0404 (B8) | MN840541 |
| *bat2* | RBRH_00895 | *Burkholderia* sp. HKI-0512 (B2) | MN840542 |
| *bat2* | RBRH_01901 | *Burkholderia rhizoxinica* HKI-0455 (B3) | MN840545 |
| *bat2* | RBRH_01694 | *Burkholderia rhizoxinica* HKI-0513 (B6) | MN840543 |
| *bat2* | RBRH_02271 | *Burkholderia rhizoxinica* HKI-0403 (B7) | MN840544 |
| *bat3* | RBRH_00894 | *Burkholderia* sp. HKI-0512 (B2) | MN840547 |
| *bat3* | RBRH_01581 | *Burkholderia rhizoxinica* HKI-0402 (B4) | MN840549 |
| *bat3* | RBRH_01695 | *Burkholderia rhizoxinica* HKI-0513 (B6) | MN840546 |
| *bat3* | RBRH_01772 | *Burkholderia rhizoxinica* HKI-0403 (B7) | MN840548 |

* Sequences obtained via amplification and Sanger sequencing.

**Supplementary Table 7.** Predicted nuclear localisation sequences (NLS) within transcription activator-like effector proteins from endofungal *Burkholderia* species (BATs) using the NucPred prediction software[2] and the cNLS Mapper[3].

| Species | BAT | NucPred Predictions[1] | | | cNLS |
|---|---|---|---|---|---|
| | protein | Score | Specificity[2] | Sensitivity[3] | Score[4] |
| *Burkholderia rhizoxinica* HKI-0454 (B1) | BAT1 | 0.23 | 0.52 | 0.83 | N/A |
| | BAT2 | 0.27 | 0.52 | 0.83 | N/A |
| | BAT3 | 0.08 | N/A | N/A | N/A |
| *Burkholderia* sp. strain HKI-0512 (B2) | BAT1 | 0.22 | 0.52 | 0.83 | N/A |
| | BAT2 | 0.27 | 0.52 | 0.83 | N/A |
| | BAT3 | 0.08 | N/A | N/A | N/A |
| *Burkholderia* sp. strain HKI-0455 (B3) | BAT2 | 0.14 | 0.45 | 0.88 | N/A |
| *Burkholderia* sp. strain HKI-0402 (B4) | BAT1 | 0.23 | 0.52 | 0.83 | N/A |
| | BAT3 | 0.09 | N/A | N/A | N/A |
| *Burkholderia endofungorum* HKI-0456 (B5) | BAT1 | 0.22 | 0.52 | 0.83 | N/A |
| *Burkholderia* sp. strain HKI-0513 (B6) | BAT1 | 0.23 | 0.52 | 0.83 | N/A |
| | BAT2 | 0.27 | 0.52 | 0.83 | N/A |
| | BAT3 | 0.08 | N/A | N/A | N/A |
| *Burkholderia* sp. strain HKI-0403 (B7) | BAT1 | 0.17 | 0.45 | 0.88 | N/A |
| | BAT2 | 0.16 | 0.45 | 0.88 | N/A |
| | BAT3 | 0.05 | N/A | N/A | N/A |
| *Burkholderia* sp. strain HKI-0404 (B8) | BAT1 | 0.12 | 0.45 | 0.88 | N/A |

[1] Nuclear localisation predicted with NucPred [2].
[2] Fraction of proteins predicted to be nuclear that actually are nuclear.
[3] Fraction of true nuclear proteins that are predicted (coverage).
[4] cNLS score: > 10: protein exclusively localised to the nucleus; 7 – 8: partially localised to the nucleus; 3 – 5: localised to both the nucleus and the cytoplasm; 1 – 2: localised to the cytoplasm.
N/A: not available.

**Supplementary Table 8. a,** Approximate probabilities (p) of Brown-Forsythe test and **b,** one-way analysis of variance (ANOVA) for the mycelium area of each morphotype following co-cultivation of *Rhizopus microsporus* with *Burkholderia rhizoxinica* wild-type (B1 WT), *B. rhizoxinica* transcription-activator like effector (BAT) mutant strains (Δ*bat1*::Apra$^r$, Δ*bat2*::Kan$^r$, Δ*bat3*::Kan$^r$), type 2 secretion system mutant strains (Δ*sctC*::Kan$^r$ and Δ*sctT*::Kan$^r$) or rhizoxin-deficient mutant strains (Δ*rhiG*::Kan$^r$). Homogeneous data (non-significant Brown-Forsythe) is shown in black numbers and non-homogeneous data (significant Brown-Forsythe) is highlighted in red numbers. **c,** Comparison of individual morphotypes at specific time points (0, 24, or 48 hours post infection; hpi) across all seven *B. rhizoxinica* strains using the Tukey HSD Post Hoc Test. **d,** Comparison of morphotypes from individual *B. rhizoxinica* strains over time (0 hpi vs 24 hpi and 0 hpi vs 48 hpi) using the Tukey HSD Post Hoc Test. P-values with *p<0.05* were considered statistically significant (highlighted in grey).

**a**

| **Brown-Forsythe Test** | |
|---|---|
| F (DFn, DFd) | 0.9933 (83, 168) |
| P value | 0.5056 |
| P value summary | ns |
| Are SDs significantly different (P < 0.05)? | No |

**b**

| **ANOVA Table** | **SS** | **DF** | **MS** | **F (DFn, DFd)** | **P value** |
|---|---|---|---|---|---|
| Treatment (between columns) | 88406 | 83 | 1065 | F (83, 168) = 8.763 | P<0.0001 |
| Residual (within columns) | 20421 | 168 | 121.6 | | |
| Total | 108827 | 251 | | | |

**c**

| | **Time** | **Strain Comparison** | | | **Mean Diff.** | **95% CI** | **P<0.05?** | **Summary** |
|---|---|---|---|---|---|---|---|---|
| **Morphotype 1** | **0 hpi** | B1 WT | vs | Δ*bat1*::Apra$^r$ | –19.67 | –58.84 to 19.51 | No | ns |
| | | B1 WT | vs | Δ*bat2*::Kan$^r$ | 4 | –35.18 to 43.18 | No | ns |
| | | B1 WT | vs | Δ*bat3*::Kan$^r$ | 9.667 | –29.51 to 48.84 | No | ns |
| | | B1 WT | vs | Δ*rhzG*::Kan$^r$ | –28.33 | –67.51 to 10.84 | No | ns |
| | | B1 WT | vs | Δ*sctC*::Kan$^r$ | –10.33 | –49.51 to 28.84 | No | ns |
| | | B1 WT | vs | Δ*sctT*::Kan$^r$ | 0.6667 | –38.51 to 39.84 | No | ns |
| | | Δ*bat1*::Apra$^r$ | vs | Δ*bat2*::Kan$^r$ | 23.67 | –15.51 to 62.84 | No | ns |
| | | Δ*bat1*::Apra$^r$ | vs | Δ*bat3*::Kan$^r$ | 29.33 | –9.842 to 68.51 | No | ns |
| | | Δ*bat1*::Apra$^r$ | vs | Δ*rhzG*::Kan$^r$ | –8.667 | –47.84 to 30.51 | No | ns |
| | | Δ*bat1*::Apra$^r$ | vs | Δ*sctC*::Kan$^r$ | 9.333 | –29.84 to 48.51 | No | ns |
| | | Δ*bat1*::Apra$^r$ | vs | Δ*sctT*::Kan$^r$ | 20.33 | –18.84 to 59.51 | No | ns |
| | | Δ*bat2*::Kan$^r$ | vs | Δ*bat3*::Kan$^r$ | 5.667 | –33.51 to 44.84 | No | ns |
| | | Δ*bat2*::Kan$^r$ | vs | Δ*rhzG*::Kan$^r$ | –32.33 | –71.51 to 6.842 | No | ns |
| | | Δ*bat2*::Kan$^r$ | vs | Δ*sctC*::Kan$^r$ | –14.33 | –53.51 to 24.84 | No | ns |
| | | Δ*bat2*::Kan$^r$ | vs | Δ*sctT*::Kan$^r$ | –3.333 | –42.51 to 35.84 | No | ns |
| | | Δ*bat3*::Kan$^r$ | vs | Δ*rhzG*::Kan$^r$ | –38 | –77.18 to 1.176 | No | ns |
| | | Δ*bat3*::Kan$^r$ | vs | Δ*sctC*::Kan$^r$ | –20 | –59.18 to 19.18 | No | ns |
| | | Δ*bat3*::Kan$^r$ | vs | Δ*sctT*::Kan$^r$ | –9 | –48.18 to 30.18 | No | ns |
| | | Δ*sctC*::Kan$^r$ | vs | Δ*rhzG*::Kan$^r$ | –18 | –57.18 to 21.18 | No | ns |
| | | Δ*sctC*::Kan$^r$ | vs | Δ*sctT*::Kan$^r$ | 11 | –28.18 to 50.18 | No | ns |
| | | Δ*sctT*::Kan$^r$ | vs | Δ*rhzG*::Kan$^r$ | –29 | –68.18 to 10.18 | No | ns |
| | **24 hpi** | B1 WT | vs | Δ*bat1*::Apra$^r$ | –3 | –42.18 to 36.18 | No | ns |

| | | | | | | | | |
|---|---|---|---|---|---|---|---|---|
| | | B1 WT | vs | Δ*bat2*::Kan$^r$ | 10 | −29.18 to 49.18 | No | ns |
| | | B1 WT | vs | Δ*bat3*::Kan$^r$ | −4.333 | −43.51 to 34.84 | No | ns |
| | | B1 WT | vs | Δ*rhzG*::Kan$^r$ | 7.333 | −31.84 to 46.51 | No | ns |
| | | B1 WT | vs | Δ*sctC*::Kan$^r$ | −18.33 | −57.51 to 20.84 | No | ns |
| | | B1 WT | vs | Δ*sctT*::Kan$^r$ | −4.333 | −43.51 to 34.84 | No | ns |
| | | Δ*bat1*::Apra$^r$ | vs | Δ*bat2*::Kan$^r$ | 13 | −26.18 to 52.18 | No | ns |
| | | Δ*bat1*::Apra$^r$ | vs | Δ*bat3*::Kan$^r$ | −1.333 | −40.51 to 37.84 | No | ns |
| | | Δ*bat1*::Apra$^r$ | vs | Δ*rhzG*::Kan$^r$ | 10.33 | −28.84 to 49.51 | No | ns |
| | | Δ*bat1*::Apra$^r$ | vs | Δ*sctC*::Kan$^r$ | −15.33 | −54.51 to 23.84 | No | ns |
| | | Δ*bat1*::Apra$^r$ | vs | Δ*sctT*::Kan$^r$ | −1.333 | −40.51 to 37.84 | No | ns |
| | | Δ*bat2*::Kan$^r$ | vs | Δ*bat3*::Kan$^r$ | −14.33 | −53.51 to 24.84 | No | ns |
| | | Δ*bat2*::Kan$^r$ | vs | Δ*rhzG*::Kan$^r$ | −2.667 | −41.84 to 36.51 | No | ns |
| | | Δ*bat2*::Kan$^r$ | vs | Δ*sctC*::Kan$^r$ | −28.33 | −67.51 to 10.84 | No | ns |
| | | Δ*bat2*::Kan$^r$ | vs | Δ*sctT*::Kan$^r$ | −14.33 | −53.51 to 24.84 | No | ns |
| | | Δ*bat3*::Kan$^r$ | vs | Δ*rhzG*::Kan$^r$ | 11.67 | −27.51 to 50.84 | No | ns |
| | | Δ*bat3*::Kan$^r$ | vs | Δ*sctC*::Kan$^r$ | −14 | −53.18 to 25.18 | No | ns |
| | | Δ*bat3*::Kan$^r$ | vs | Δ*sctT*::Kan$^r$ | 0 | −39.18 to 39.18 | No | ns |
| | | Δ*sctC*::Kan$^r$ | vs | Δ*rhzG*::Kan$^r$ | 25.67 | −13.51 to 64.84 | No | ns |
| | | Δ*sctC*::Kan$^r$ | vs | Δ*sctT*::Kan$^r$ | 14 | −25.18 to 53.18 | No | ns |
| | | Δ*sctT*::Kan$^r$ | vs | Δ*rhzG*::Kan$^r$ | 11.67 | −27.51 to 50.84 | No | ns |
| | **48 hpi** | B1 WT | vs | Δ*bat1*::Apra$^r$ | 27.67 | −11.51 to 66.84 | No | ns |
| | | B1 WT | vs | Δ*bat2*::Kan$^r$ | 9.667 | −29.51 to 48.84 | No | ns |
| | | B1 WT | vs | Δ*bat3*::Kan$^r$ | −9 | −48.18 to 30.18 | No | ns |
| | | B1 WT | vs | Δ*rhzG*::Kan$^r$ | 24.33 | −14.84 to 63.51 | No | ns |
| | | B1 WT | vs | Δ*sctC*::Kan$^r$ | 11 | −28.18 to 50.18 | No | ns |
| | | B1 WT | vs | Δ*sctT*::Kan$^r$ | 13.33 | −25.84 to 52.51 | No | ns |
| | | Δ*bat1*::Apra$^r$ | vs | Δ*bat2*::Kan$^r$ | −18 | −57.18 to 21.18 | No | ns |
| | | Δ*bat1*::Apra$^r$ | vs | Δ*bat3*::Kan$^r$ | −36.67 | −75.84 to 2.509 | No | ns |
| | | Δ*bat1*::Apra$^r$ | vs | Δ*rhzG*::Kan$^r$ | −3.333 | −42.51 to 35.84 | No | ns |
| | | Δ*bat1*::Apra$^r$ | vs | Δ*sctC*::Kan$^r$ | −16.67 | −55.84 to 22.51 | No | ns |
| | | Δ*bat1*::Apra$^r$ | vs | Δ*sctT*::Kan$^r$ | −14.33 | −53.51 to 24.84 | No | ns |
| | | Δ*bat2*::Kan$^r$ | vs | Δ*bat3*::Kan$^r$ | −18.67 | −57.84 to 20.51 | No | ns |
| | | Δ*bat2*::Kan$^r$ | vs | Δ*rhzG*::Kan$^r$ | 14.67 | −24.51 to 53.84 | No | ns |
| | | Δ*bat2*::Kan$^r$ | vs | Δ*sctC*::Kan$^r$ | 1.333 | −37.84 to 40.51 | No | ns |
| | | Δ*bat2*::Kan$^r$ | vs | Δ*sctT*::Kan$^r$ | 3.667 | −35.51 to 42.84 | No | ns |
| | | Δ*bat3*::Kan$^r$ | vs | Δ*rhzG*::Kan$^r$ | 33.33 | −5.842 to 72.51 | No | ns |
| | | Δ*bat3*::Kan$^r$ | vs | Δ*sctC*::Kan$^r$ | 20 | −19.18 to 59.18 | No | ns |
| | | Δ*bat3*::Kan$^r$ | vs | Δ*sctT*::Kan$^r$ | 22.33 | −16.84 to 61.51 | No | ns |
| | | Δ*sctC*::Kan$^r$ | vs | Δ*rhzG*::Kan$^r$ | 13.33 | −25.84 to 52.51 | No | ns |
| | | Δ*sctC*::Kan$^r$ | vs | Δ*sctT*::Kan$^r$ | 2.333 | −36.84 to 41.51 | No | ns |
| | | Δ*sctT*::Kan$^r$ | vs | Δ*rhzG*::Kan$^r$ | 11 | −28.18 to 50.18 | No | ns |
| **Morphotype 2** | **0 hpi** | B1 WT | vs | Δ*bat1*::Apra$^r$ | 18 | −21.18 to 57.18 | No | ns |
| | | B1 WT | vs | Δ*bat2*::Kan$^r$ | 3.333 | −35.84 to 42.51 | No | ns |
| | | B1 WT | vs | Δ*bat3*::Kan$^r$ | 2.667 | −36.51 to 41.84 | No | ns |
| | | B1 WT | vs | Δ*rhzG*::Kan$^r$ | 28.33 | −10.84 to 67.51 | No | ns |
| | | B1 WT | vs | Δ*sctC*::Kan$^r$ | 9.667 | −29.51 to 48.84 | No | ns |

| | | | | | | | |
|---|---|---|---|---|---|---|---|
| | B1 WT | vs | Δ*sctT*::Kan$^r$ | −1 | −40.18 to 38.18 | No | ns |
| | Δ*bat1*::Apra$^r$ | vs | Δ*bat2*::Kan$^r$ | −14.67 | −53.84 to 24.51 | No | ns |
| | Δ*bat1*::Apra$^r$ | vs | Δ*bat3*::Kan$^r$ | −15.33 | −54.51 to 23.84 | No | ns |
| | Δ*bat1*::Apra$^r$ | vs | Δ*rhzG*::Kan$^r$ | 10.33 | −28.84 to 49.51 | No | ns |
| | Δ*bat1*::Apra$^r$ | vs | Δ*sctC*::Kan$^r$ | −8.333 | −47.51 to 30.84 | No | ns |
| | Δ*bat1*::Apra$^r$ | vs | Δ*sctT*::Kan$^r$ | −19 | −58.18 to 20.18 | No | ns |
| | Δ*bat2*::Kan$^r$ | vs | Δ*bat3*::Kan$^r$ | −0.6667 | −39.84 to 38.51 | No | ns |
| | Δ*bat2*::Kan$^r$ | vs | Δ*rhzG*::Kan$^r$ | 25 | −14.18 to 64.18 | No | ns |
| | Δ*bat2*::Kan$^r$ | vs | Δ*sctC*::Kan$^r$ | 6.333 | −32.84 to 45.51 | No | ns |
| | Δ*bat2*::Kan$^r$ | vs | Δ*sctT*::Kan$^r$ | −4.333 | −43.51 to 34.84 | No | ns |
| | Δ*bat3*::Kan$^r$ | vs | Δ*rhzG*::Kan$^r$ | 25.67 | −13.51 to 64.84 | No | ns |
| | Δ*bat3*::Kan$^r$ | vs | Δ*sctC*::Kan$^r$ | 7 | −32.18 to 46.18 | No | ns |
| | Δ*bat3*::Kan$^r$ | vs | Δ*sctT*::Kan$^r$ | −3.667 | −42.84 to 35.51 | No | ns |
| | Δ*sctC*::Kan$^r$ | vs | Δ*rhzG*::Kan$^r$ | 18.67 | −20.51 to 57.84 | No | ns |
| | Δ*sctC*::Kan$^r$ | vs | Δ*sctT*::Kan$^r$ | −10.67 | −49.84 to 28.51 | No | ns |
| | Δ*sctT*::Kan$^r$ | vs | Δ*rhzG*::Kan$^r$ | 29.33 | −9.842 to 68.51 | No | ns |
| **24 hpi** | B1 WT | vs | Δ*bat1*::Apra$^r$ | 8.667 | −30.51 to 47.84 | No | ns |
| | B1 WT | vs | Δ*bat2*::Kan$^r$ | 16.67 | −22.51 to 55.84 | No | ns |
| | B1 WT | vs | Δ*bat3*::Kan$^r$ | 7 | −32.18 to 46.18 | No | ns |
| | B1 WT | vs | Δ*rhzG*::Kan$^r$ | −18 | −57.18 to 21.18 | No | ns |
| | B1 WT | vs | Δ*sctC*::Kan$^r$ | 27.33 | −11.84 to 66.51 | No | ns |
| | B1 WT | vs | Δ*sctT*::Kan$^r$ | 7.667 | −31.51 to 46.84 | No | ns |
| | Δ*bat1*::Apra$^r$ | vs | Δ*bat2*::Kan$^r$ | 8 | −31.18 to 47.18 | No | ns |
| | Δ*bat1*::Apra$^r$ | vs | Δ*bat3*::Kan$^r$ | −1.667 | −40.84 to 37.51 | No | ns |
| | Δ*bat1*::Apra$^r$ | vs | Δ*rhzG*::Kan$^r$ | −26.67 | −65.84 to 12.51 | No | ns |
| | Δ*bat1*::Apra$^r$ | vs | Δ*sctC*::Kan$^r$ | 18.67 | −20.51 to 57.84 | No | ns |
| | Δ*bat1*::Apra$^r$ | vs | Δ*sctT*::Kan$^r$ | −1 | −40.18 to 38.18 | No | ns |
| | Δ*bat2*::Kan$^r$ | vs | Δ*bat3*::Kan$^r$ | −9.667 | −48.84 to 29.51 | No | ns |
| | Δ*bat2*::Kan$^r$ | vs | Δ*rhzG*::Kan$^r$ | −34.67 | −73.84 to 4.509 | No | ns |
| | Δ*bat2*::Kan$^r$ | vs | Δ*sctC*::Kan$^r$ | 10.67 | −28.51 to 49.84 | No | ns |
| | Δ*bat2*::Kan$^r$ | vs | Δ*sctT*::Kan$^r$ | −9 | −48.18 to 30.18 | No | ns |
| | Δ*bat3*::Kan$^r$ | vs | Δ*rhzG*::Kan$^r$ | −25 | −64.18 to 14.18 | No | ns |
| | Δ*bat3*::Kan$^r$ | vs | Δ*sctC*::Kan$^r$ | 20.33 | −18.84 to 59.51 | No | ns |
| | Δ*bat3*::Kan$^r$ | vs | Δ*sctT*::Kan$^r$ | 0.6667 | −38.51 to 39.84 | No | ns |
| | Δ*sctC*::Kan$^r$ | vs | Δ*rhzG*::Kan$^r$ | −45.33 | −84.51 to −6.158 | Yes | ** |
| | Δ*sctC*::Kan$^r$ | vs | Δ*sctT*::Kan$^r$ | −19.67 | −58.84 to 19.51 | No | ns |
| | Δ*sctT*::Kan$^r$ | vs | Δ*rhzG*::Kan$^r$ | −25.67 | −64.84 to 13.51 | No | ns |
| **48 hpi** | B1 WT | vs | Δ*bat1*::Apra$^r$ | −25 | −64.18 to 14.18 | No | ns |
| | B1 WT | vs | Δ*bat2*::Kan$^r$ | 9.667 | −29.51 to 48.84 | No | ns |
| | B1 WT | vs | Δ*bat3*::Kan$^r$ | 18.33 | −20.84 to 57.51 | No | ns |
| | B1 WT | vs | Δ*rhzG*::Kan$^r$ | −21 | −60.18 to 18.18 | No | ns |
| | B1 WT | vs | Δ*sctC*::Kan$^r$ | 1.333 | −37.84 to 40.51 | No | ns |
| | B1 WT | vs | Δ*sctT*::Kan$^r$ | −4.667 | −43.84 to 34.51 | No | ns |
| | Δ*bat1*::Apra$^r$ | vs | Δ*bat2*::Kan$^r$ | 34.67 | −4.509 to 73.84 | No | ns |
| | Δ*bat1*::Apra$^r$ | vs | Δ*bat3*::Kan$^r$ | 43.33 | 4.158 to 82.51 | Yes | ** |
| | Δ*bat1*::Apra$^r$ | vs | Δ*rhzG*::Kan$^r$ | 4 | −35.18 to 43.18 | No | ns |

| | | | | | | | | |
|---|---|---|---|---|---|---|---|---|
| | | Δ*bat1*::Apra$^r$ | vs | Δ*sctC*::Kan$^r$ | 26.33 | −12.84 to 65.51 | No | ns |
| | | Δ*bat1*::Apra$^r$ | vs | Δ*sctT*::Kan$^r$ | 20.33 | −18.84 to 59.51 | No | ns |
| | | Δ*bat2*::Kan$^r$ | vs | Δ*bat3*::Kan$^r$ | 8.667 | −30.51 to 47.84 | No | ns |
| | | Δ*bat2*::Kan$^r$ | vs | Δ*rhzG*::Kan$^r$ | −30.67 | −69.84 to 8.509 | No | ns |
| | | Δ*bat2*::Kan$^r$ | vs | Δ*sctC*::Kan$^r$ | −8.333 | −47.51 to 30.84 | No | ns |
| | | Δ*bat2*::Kan$^r$ | vs | Δ*sctT*::Kan$^r$ | −14.33 | −53.51 to 24.84 | No | ns |
| | | Δ*bat3*::Kan$^r$ | vs | Δ*rhzG*::Kan$^r$ | −39.33 | −78.51 to −0.1575 | Yes | * |
| | | Δ*bat3*::Kan$^r$ | vs | Δ*sctC*::Kan$^r$ | −17 | −56.18 to 22.18 | No | ns |
| | | Δ*bat3*::Kan$^r$ | vs | Δ*sctT*::Kan$^r$ | −23 | −62.18 to 16.18 | No | ns |
| | | Δ*sctC*::Kan$^r$ | vs | Δ*rhzG*::Kan$^r$ | −22.33 | −61.51 to 16.84 | No | ns |
| | | Δ*sctC*::Kan$^r$ | vs | Δ*sctT*::Kan$^r$ | −6 | −45.18 to 33.18 | No | ns |
| | | Δ*sctT*::Kan$^r$ | vs | Δ*rhzG*::Kan$^r$ | −16.33 | −55.51 to 22.84 | No | ns |
| **Morphotype 3** | **0 hpi** | B1 WT | vs | Δ*bat1*::Apra$^r$ | 3.333 | −35.84 to 42.51 | No | ns |
| | | B1 WT | vs | Δ*bat2*::Kan$^r$ | −5.333 | −44.51 to 33.84 | No | ns |
| | | B1 WT | vs | Δ*bat3*::Kan$^r$ | −10.33 | −49.51 to 28.84 | No | ns |
| | | B1 WT | vs | Δ*rhzG*::Kan$^r$ | 0.6667 | −38.51 to 39.84 | No | ns |
| | | B1 WT | vs | Δ*sctC*::Kan$^r$ | 1 | −38.18 to 40.18 | No | ns |
| | | B1 WT | vs | Δ*sctT*::Kan$^r$ | 1 | −38.18 to 40.18 | No | ns |
| | | Δ*bat1*::Apra$^r$ | vs | Δ*bat2*::Kan$^r$ | −8.667 | −47.84 to 30.51 | No | ns |
| | | Δ*bat1*::Apra$^r$ | vs | Δ*bat3*::Kan$^r$ | −13.67 | −52.84 to 25.51 | No | ns |
| | | Δ*bat1*::Apra$^r$ | vs | Δ*rhzG*::Kan$^r$ | −2.667 | −41.84 to 36.51 | No | ns |
| | | Δ*bat1*::Apra$^r$ | vs | Δ*sctC*::Kan$^r$ | −2.333 | −41.51 to 36.84 | No | ns |
| | | Δ*bat1*::Apra$^r$ | vs | Δ*sctT*::Kan$^r$ | −2.333 | −41.51 to 36.84 | No | ns |
| | | Δ*bat2*::Kan$^r$ | vs | Δ*bat3*::Kan$^r$ | −5 | −44.18 to 34.18 | No | ns |
| | | Δ*bat2*::Kan$^r$ | vs | Δ*rhzG*::Kan$^r$ | 6 | −33.18 to 45.18 | No | ns |
| | | Δ*bat2*::Kan$^r$ | vs | Δ*sctC*::Kan$^r$ | 6.333 | −32.84 to 45.51 | No | ns |
| | | Δ*bat2*::Kan$^r$ | vs | Δ*sctT*::Kan$^r$ | 6.333 | −32.84 to 45.51 | No | ns |
| | | Δ*bat3*::Kan$^r$ | vs | Δ*rhzG*::Kan$^r$ | 11 | −28.18 to 50.18 | No | ns |
| | | Δ*bat3*::Kan$^r$ | vs | Δ*sctC*::Kan$^r$ | 11.33 | −27.84 to 50.51 | No | ns |
| | | Δ*bat3*::Kan$^r$ | vs | Δ*sctT*::Kan$^r$ | 11.33 | −27.84 to 50.51 | No | ns |
| | | Δ*sctC*::Kan$^r$ | vs | Δ*rhzG*::Kan$^r$ | −0.3333 | −39.51 to 38.84 | No | ns |
| | | Δ*sctC*::Kan$^r$ | vs | Δ*sctT*::Kan$^r$ | 0 | −39.18 to 39.18 | No | ns |
| | | Δ*sctT*::Kan$^r$ | vs | Δ*rhzG*::Kan$^r$ | −0.3333 | −39.51 to 38.84 | No | ns |
| | **24 hpi** | B1 WT | vs | Δ*bat1*::Apra$^r$ | 1 | −38.18 to 40.18 | No | ns |
| | | B1 WT | vs | Δ*bat2*::Kan$^r$ | −20 | −59.18 to 19.18 | No | ns |
| | | B1 WT | vs | Δ*bat3*::Kan$^r$ | 2 | −37.18 to 41.18 | No | ns |
| | | B1 WT | vs | Δ*rhzG*::Kan$^r$ | 10.33 | −28.84 to 49.51 | No | ns |
| | | B1 WT | vs | Δ*sctC*::Kan$^r$ | −8.667 | −47.84 to 30.51 | No | ns |
| | | B1 WT | vs | Δ*sctT*::Kan$^r$ | −3.333 | −42.51 to 35.84 | No | ns |
| | | Δ*bat1*::Apra$^r$ | vs | Δ*bat2*::Kan$^r$ | −21 | −60.18 to 18.18 | No | ns |
| | | Δ*bat1*::Apra$^r$ | vs | Δ*bat3*::Kan$^r$ | 1 | −38.18 to 40.18 | No | ns |
| | | Δ*bat1*::Apra$^r$ | vs | Δ*rhzG*::Kan$^r$ | 9.333 | −29.84 to 48.51 | No | ns |
| | | Δ*bat1*::Apra$^r$ | vs | Δ*sctC*::Kan$^r$ | −9.667 | −48.84 to 29.51 | No | ns |
| | | Δ*bat1*::Apra$^r$ | vs | Δ*sctT*::Kan$^r$ | −4.333 | −43.51 to 34.84 | No | ns |
| | | Δ*bat2*::Kan$^r$ | vs | Δ*bat3*::Kan$^r$ | 22 | −17.18 to 61.18 | No | ns |
| | | Δ*bat2*::Kan$^r$ | vs | Δ*rhzG*::Kan$^r$ | 30.33 | −8.842 to 69.51 | No | ns |

| | | | | | | | | |
|---|---|---|---|---|---|---|---|---|
| | | Δ*bat2*::Kan$^r$ | vs | Δ*sctC*::Kan$^r$ | 11.33 | −27.84 to 50.51 | No | ns |
| | | Δ*bat2*::Kan$^r$ | vs | Δ*sctT*::Kan$^r$ | 16.67 | −22.51 to 55.84 | No | ns |
| | | Δ*bat3*::Kan$^r$ | vs | Δ*rhzG*::Kan$^r$ | 8.333 | −30.84 to 47.51 | No | ns |
| | | Δ*bat3*::Kan$^r$ | vs | Δ*sctC*::Kan$^r$ | −10.67 | −49.84 to 28.51 | No | ns |
| | | Δ*bat3*::Kan$^r$ | vs | Δ*sctT*::Kan$^r$ | −5.333 | −44.51 to 33.84 | No | ns |
| | | Δ*sctC*::Kan$^r$ | vs | Δ*rhzG*::Kan$^r$ | 19 | −20.18 to 58.18 | No | ns |
| | | Δ*sctC*::Kan$^r$ | vs | Δ*sctT*::Kan$^r$ | 5.333 | −33.84 to 44.51 | No | ns |
| | | Δ*sctT*::Kan$^r$ | vs | Δ*rhzG*::Kan$^r$ | 13.67 | −25.51 to 52.84 | No | ns |
| | **48 hpi** | B1 WT | vs | Δ*bat1*::Apra$^r$ | −4 | −43.18 to 35.18 | No | ns |
| | | B1 WT | vs | Δ*bat2*::Kan$^r$ | −12.67 | −51.84 to 26.51 | No | ns |
| | | B1 WT | vs | Δ*bat3*::Kan$^r$ | −4 | −43.18 to 35.18 | No | ns |
| | | B1 WT | vs | Δ*rhzG*::Kan$^r$ | −8.667 | −47.84 to 30.51 | No | ns |
| | | B1 WT | vs | Δ*sctC*::Kan$^r$ | −18.33 | −57.51 to 20.84 | No | ns |
| | | B1 WT | vs | Δ*sctT*::Kan$^r$ | −4.33 | −53.51 to 24.84 | No | ns |
| | | Δ*bat1*::Apra$^r$ | vs | Δ*bat2*::Kan$^r$ | −8.667 | −47.84 to 30.51 | No | ns |
| | | Δ*bat1*::Apra$^r$ | vs | Δ*bat3*::Kan$^r$ | 0 | −39.18 to 39.18 | No | ns |
| | | Δ*bat1*::Apra$^r$ | vs | Δ*rhzG*::Kan$^r$ | −4.667 | −43.84 to 34.51 | No | ns |
| | | Δ*bat1*::Apra$^r$ | vs | Δ*sctC*::Kan$^r$ | −14.33 | −53.51 to 24.84 | No | ns |
| | | Δ*bat1*::Apra$^r$ | vs | Δ*sctT*::Kan$^r$ | −10.33 | −49.51 to 28.84 | No | ns |
| | | Δ*bat2*::Kan$^r$ | vs | Δ*bat3*::Kan$^r$ | 8.667 | −30.51 to 47.84 | No | ns |
| | | Δ*bat2*::Kan$^r$ | vs | Δ*rhzG*::Kan$^r$ | 4 | −35.18 to 43.18 | No | ns |
| | | Δ*bat2*::Kan$^r$ | vs | Δ*sctC*::Kan$^r$ | −5.667 | −44.84 to 33.51 | No | ns |
| | | Δ*bat2*::Kan$^r$ | vs | Δ*sctT*::Kan$^r$ | −1.667 | −40.84 to 37.51 | No | ns |
| | | Δ*bat3*::Kan$^r$ | vs | Δ*rhzG*::Kan$^r$ | −4.667 | −43.84 to 34.51 | No | ns |
| | | Δ*bat3*::Kan$^r$ | vs | Δ*sctC*::Kan$^r$ | −14.33 | −53.51 to 24.84 | No | ns |
| | | Δ*bat3*::Kan$^r$ | vs | Δ*sctT*::Kan$^r$ | −10.33 | −49.51 to 28.84 | No | ns |
| | | Δ*sctC*::Kan$^r$ | vs | Δ*rhzG*::Kan$^r$ | 9.667 | −29.51 to 48.84 | No | ns |
| | | Δ*sctC*::Kan$^r$ | vs | Δ*sctT*::Kan$^r$ | 4 | −35.18 to 43.18 | No | ns |
| | | Δ*sctT*::Kan$^r$ | vs | Δ*rhzG*::Kan$^r$ | 5.667 | −33.51 to 44.84 | No | ns |
| **Morphotype 4** | **0 hpi** | B1 WT | vs | Δ*bat1*::Apra$^r$ | −0.3333 | −39.51 to 38.84 | No | ns |
| | | B1 WT | vs | Δ*bat2*::Kan$^r$ | 0.3333 | −38.84 to 39.51 | No | ns |
| | | B1 WT | vs | Δ*bat3*::Kan$^r$ | −0.3333 | −39.51 to 38.84 | No | ns |
| | | B1 WT | vs | Δ*rhzG*::Kan$^r$ | 1 | −38.18 to 40.18 | No | ns |
| | | B1 WT | vs | Δ*sctC*::Kan$^r$ | 1 | −38.18 to 40.18 | No | ns |
| | | B1 WT | vs | Δ*sctT*::Kan$^r$ | 1 | −38.18 to 40.18 | No | ns |
| | | Δ*bat1*::Apra$^r$ | vs | Δ*bat2*::Kan$^r$ | 0.6667 | −38.51 to 39.84 | No | ns |
| | | Δ*bat1*::Apra$^r$ | vs | Δ*bat3*::Kan$^r$ | 0 | −39.18 to 39.18 | No | ns |
| | | Δ*bat1*::Apra$^r$ | vs | Δ*rhzG*::Kan$^r$ | 1.333 | −37.84 to 40.51 | No | ns |
| | | Δ*bat1*::Apra$^r$ | vs | Δ*sctC*::Kan$^r$ | 1.333 | −37.84 to 40.51 | No | ns |
| | | Δ*bat1*::Apra$^r$ | vs | Δ*sctT*::Kan$^r$ | 1.333 | −37.84 to 40.51 | No | ns |
| | | Δ*bat2*::Kan$^r$ | vs | Δ*bat3*::Kan$^r$ | −0.6667 | −39.84 to 38.51 | No | ns |
| | | Δ*bat2*::Kan$^r$ | vs | Δ*rhzG*::Kan$^r$ | 0.6667 | −38.51 to 39.84 | No | ns |
| | | Δ*bat2*::Kan$^r$ | vs | Δ*sctC*::Kan$^r$ | 0.6667 | −38.51 to 39.84 | No | ns |
| | | Δ*bat2*::Kan$^r$ | vs | Δ*sctT*::Kan$^r$ | 0.6667 | −38.51 to 39.84 | No | ns |
| | | Δ*bat3*::Kan$^r$ | vs | Δ*rhzG*::Kan$^r$ | 1.333 | −37.84 to 40.51 | No | ns |
| | | Δ*bat3*::Kan$^r$ | vs | Δ*sctC*::Kan$^r$ | 1.333 | −37.84 to 40.51 | No | ns |

| | | | | | | | |
|---|---|---|---|---|---|---|---|
| | Δ*bat3*::Kan$^r$ | vs | Δ*sctT*::Kan$^r$ | 1.333 | −37.84 to 40.51 | No | ns |
| | Δ*sctC*::Kan$^r$ | vs | Δ*rhzG*::Kan$^r$ | 0 | −39.18 to 39.18 | No | ns |
| | Δ*sctC*::Kan$^r$ | vs | Δ*sctT*::Kan$^r$ | 0 | −39.18 to 39.18 | No | ns |
| | Δ*sctT*::Kan$^r$ | vs | Δ*rhzG*::Kan$^r$ | 0 | −39.18 to 39.18 | No | ns |
| **24 hpi** | B1 WT | vs | Δ*bat1*::Apra$^r$ | 2 | −37.18 to 41.18 | No | ns |
| | B1 WT | vs | Δ*bat2*::Kan$^r$ | 1 | −38.18 to 40.18 | No | ns |
| | B1 WT | vs | Δ*bat3*::Kan$^r$ | 1.333 | −37.84 to 40.51 | No | ns |
| | B1 WT | vs | Δ*rhzG*::Kan$^r$ | 9.667 | −29.51 to 48.84 | No | ns |
| | B1 WT | vs | Δ*sctC*::Kan$^r$ | 8.333 | −30.84 to 47.51 | No | ns |
| | B1 WT | vs | Δ*sctT*::Kan$^r$ | 9 | −30.18 to 48.18 | No | ns |
| | Δ*bat1*::Apra$^r$ | vs | Δ*bat2*::Kan$^r$ | −1 | −40.18 to 38.18 | No | ns |
| | Δ*bat1*::Apra$^r$ | vs | Δ*bat3*::Kan$^r$ | −0.6667 | −39.84 to 38.51 | No | ns |
| | Δ*bat1*::Apra$^r$ | vs | Δ*rhzG*::Kan$^r$ | 7.667 | −31.51 to 46.84 | No | ns |
| | Δ*bat1*::Apra$^r$ | vs | Δ*sctC*::Kan$^r$ | 6.333 | −32.84 to 45.51 | No | ns |
| | Δ*bat1*::Apra$^r$ | vs | Δ*sctT*::Kan$^r$ | 7 | −32.18 to 46.18 | No | ns |
| | Δ*bat2*::Kan$^r$ | vs | Δ*bat3*::Kan$^r$ | 0.3333 | −38.84 to 39.51 | No | ns |
| | Δ*bat2*::Kan$^r$ | vs | Δ*rhzG*::Kan$^r$ | 8.667 | −30.51 to 47.84 | No | ns |
| | Δ*bat2*::Kan$^r$ | vs | Δ*sctC*::Kan$^r$ | 7.333 | −31.84 to 46.51 | No | ns |
| | Δ*bat2*::Kan$^r$ | vs | Δ*sctT*::Kan$^r$ | 8 | −31.18 to 47.18 | No | ns |
| | Δ*bat3*::Kan$^r$ | vs | Δ*rhzG*::Kan$^r$ | 8.333 | −30.84 to 47.51 | No | ns |
| | Δ*bat3*::Kan$^r$ | vs | Δ*sctC*::Kan$^r$ | 7 | −32.18 to 46.18 | No | ns |
| | Δ*bat3*::Kan$^r$ | vs | Δ*sctT*::Kan$^r$ | 7 | −32.18 to 46.18 | No | ns |
| | Δ*sctC*::Kan$^r$ | vs | Δ*rhzG*::Kan$^r$ | 1.333 | −37.84 to 40.51 | No | ns |
| | Δ*sctC*::Kan$^r$ | vs | Δ*sctT*::Kan$^r$ | 0.6667 | −38.51 to 39.84 | No | ns |
| | Δ*sctT*::Kan$^r$ | vs | Δ*rhzG*::Kan$^r$ | 0.6667 | −38.51 to 39.84 | No | ns |
| **48 hpi** | B1 WT | vs | Δ*bat1*::Apra$^r$ | −1 | −40.18 to 38.18 | No | ns |
| | B1 WT | vs | Δ*bat2*::Kan$^r$ | −8 | −47.18 to 31.18 | No | ns |
| | B1 WT | vs | Δ*bat3*::Kan$^r$ | −4.333 | −43.51 to 34.84 | No | ns |
| | B1 WT | vs | Δ*rhzG*::Kan$^r$ | 4.667 | −34.51 to 43.84 | No | ns |
| | B1 WT | vs | Δ*sctC*::Kan$^r$ | 5 | −34.18 to 44.18 | No | ns |
| | B1 WT | vs | Δ*sctT*::Kan$^r$ | 4.667 | −34.51 to 43.84 | No | ns |
| | Δ*bat1*::Apra$^r$ | vs | Δ*bat2*::Kan$^r$ | −7 | −46.18 to 32.18 | No | ns |
| | Δ*bat1*::Apra$^r$ | vs | Δ*bat3*::Kan$^r$ | −3.333 | −42.51 to 35.84 | No | ns |
| | Δ*bat1*::Apra$^r$ | vs | Δ*rhzG*::Kan$^r$ | 5.667 | −33.51 to 44.84 | No | ns |
| | Δ*bat1*::Apra$^r$ | vs | Δ*sctC*::Kan$^r$ | 6 | −33.18 to 45.18 | No | ns |
| | Δ*bat1*::Apra$^r$ | vs | Δ*sctT*::Kan$^r$ | 5.667 | −33.51 to 44.84 | No | ns |
| | Δ*bat2*::Kan$^r$ | vs | Δ*bat3*::Kan$^r$ | 3.667 | −35.51 to 42.84 | No | ns |
| | Δ*bat2*::Kan$^r$ | vs | Δ*rhzG*::Kan$^r$ | 12.67 | −26.51 to 51.84 | No | ns |
| | Δ*bat2*::Kan$^r$ | vs | Δ*sctC*::Kan$^r$ | 13 | −26.18 to 52.18 | No | ns |
| | Δ*bat2*::Kan$^r$ | vs | Δ*sctT*::Kan$^r$ | 12.67 | −26.51 to 51.84 | No | ns |
| | Δ*bat3*::Kan$^r$ | vs | Δ*rhzG*::Kan$^r$ | 9 | −30.18 to 48.18 | No | ns |
| | Δ*bat3*::Kan$^r$ | vs | Δ*sctC*::Kan$^r$ | 9.333 | −29.84 to 48.51 | No | ns |
| | Δ*bat3*::Kan$^r$ | vs | Δ*sctT*::Kan$^r$ | 9 | −30.18 to 48.18 | No | ns |
| | Δ*sctC*::Kan$^r$ | vs | Δ*rhzG*::Kan$^r$ | −0.3333 | −39.51 to 38.84 | No | ns |
| | Δ*sctC*::Kan$^r$ | vs | Δ*sctT*::Kan$^r$ | −0.3333 | −39.51 to 38.84 | No | ns |

| | | | | | | | |
|---|---|---|---|---|---|---|---|
| | Δ*sctT*::Kan$^{r}$ | vs | Δ*rhzG*::Kan$^{r}$ | 0 | −39.18 to 39.18 | No | ns |

**d**

| Strain | Morphotype | Time Comparison | Mean Diff. | 95% CI | P<0.05? | Summary |
|---|---|---|---|---|---|---|
| **B1 WT** | Morphotype 1 | 0 hpi vs 24 hpi | −2 | −27,93 to 23,93 | No | ns |
| | | 0 hpi vs 48 hpi | −7,667 | −33,59 to 18,26 | No | ns |
| | Morphotype 2 | 0 hpi vs 24 hpi | 9,333 | −16,59 to 35,26 | No | ns |
| | | 0 hpi vs 48 hpi | 16,33 | −9,592 to 42,26 | No | ns |
| | Morphotype 3 | 0 hpi vs 24 hpi | −6,333 | −32,26 to 19,59 | No | ns |
| | | 0 hpi vs 48 hpi | −1 | −26,93 to 24,93 | No | ns |
| | Morphotype 4 | 0 hpi vs 24 hpi | −8,667 | −34,59 to 17,26 | No | ns |
| | | 0 hpi vs 48 hpi | −5 | −30,93 to 20,93 | No | ns |
| **Δ*bat1*::Apra$^{r}$** | Morphotype 1 | 0 hpi vs 24 hpi | 14,67 | −11,26 to 40,59 | No | ns |
| | | 0 hpi vs 48 hpi | 39,67 | 13,74 to 65,59 | Yes | *** |
| | Morphotype 2 | 0 hpi vs 24 hpi | 0 | −25,93 to 25,93 | No | ns |
| | | 0 hpi vs 48 hpi | −26,67 | −52,59 to −0,7415 | Yes | * |
| | Morphotype 3 | 0 hpi vs 24 hpi | −8,667 | −34,59 to 17,26 | No | ns |
| | | 0 hpi vs 48 hpi | −8,333 | −34,26 to 17,59 | No | ns |
| | Morphotype 4 | 0 hpi vs 24 hpi | −6,333 | −32,26 to 19,59 | No | ns |
| | | 0 hpi vs 48 hpi | −5,667 | −31,59 to 20,26 | No | ns |
| **Δ*bat2*::Kan$^{r}$** | Morphotype 1 | 0 hpi vs 24 hpi | 4 | −21,93 to 29,93 | No | ns |
| | | 0 hpi vs 48 hpi | −2 | −27,93 to 23,93 | No | ns |
| | Morphotype 2 | 0 hpi vs 24 hpi | 22,67 | −3,259 to 48,59 | No | ns |
| | | 0 hpi vs 48 hpi | 22,67 | −3,259 to 48,59 | No | ns |
| | Morphotype 3 | 0 hpi vs 24 hpi | −21 | −46,93 to 4,925 | No | ns |
| | | 0 hpi vs 48 hpi | −8,333 | −34,26 to 17,59 | No | ns |
| | Morphotype 4 | 0 hpi vs 24 hpi | −8 | −33,93 to 17,93 | No | ns |
| | | 0 hpi vs 48 hpi | −13,33 | −39,26 to 12,59 | No | ns |
| **Δ*bat3*::Kan$^{r}$** | Morphotype 1 | 0 hpi vs 24 hpi | −16 | −41,93 to 9,925 | No | ns |
| | | 0 hpi vs 48 hpi | −26,33 | −52,26 to −0,4081 | Yes | * |
| | Morphotype 2 | 0 hpi vs 24 hpi | 13,67 | −12,26 to 39,59 | No | ns |
| | | 0 hpi vs 48 hpi | 32 | 6,075 to 57,93 | Yes | ** |
| | Morphotype 3 | 0 hpi vs 24 hpi | 6 | −19,93 to 31,93 | No | ns |
| | | 0 hpi vs 48 hpi | 5,333 | −20,59 to 31,26 | No | ns |
| | Morphotype 4 | 0 hpi vs 24 hpi | −7 | −32,93 to 18,93 | No | ns |
| | | 0 hpi vs 48 hpi | −9 | −34,93 to 16,93 | No | ns |
| **Δ*rhzG*::Kan$^{r}$** | Morphotype 1 | 0 hpi vs 24 hpi | 33,67 | −5,509 to 72,84 | No | ns |
| | | 0 hpi vs 48 hpi | 45 | 5,824 to 84,18 | Yes | ** |
| | Morphotype 2 | 0 hpi vs 24 hpi | −37 | −76,18 to 2,176 | No | ns |
| | | 0 hpi vs 48 hpi | −33 | −72,18 to 6,176 | No | ns |
| | Morphotype 3 | 0 hpi vs 24 hpi | 3,333 | −35,84 to 42,51 | No | ns |
| | | 0 hpi vs 48 hpi | −10,33 | −49,51 to 28,84 | No | ns |
| | Morphotype 4 | 0 hpi vs 24 hpi | 0 | −39,18 to 39,18 | No | ns |
| | | 0 hpi vs 48 hpi | −1,333 | −40,51 to 37,84 | No | ns |
| [illegible] | Morphotype 1 | 0 hpi vs 24 hpi | −10 | −49,18 to 29,18 | No | ns |

| | | | | | | |
|---|---|---|---|---|---|---|
| | | 0 hpi vs 48 hpi | 13,67 | −25,51 to 52,84 | No | ns |
| | Morphotype 2 | 0 hpi vs 24 hpi | 27 | −12,18 to 66,18 | No | ns |
| | | 0 hpi vs 48 hpi | 8 | −31,18 to 47,18 | No | ns |
| | Morphotype 3 | 0 hpi vs 24 hpi | −16 | −55,18 to 23,18 | No | ns |
| | | 0 hpi vs 48 hpi | −20,33 | −59,51 to 18,84 | No | ns |
| | Morphotype 4 | 0 hpi vs 24 hpi | −1,333 | −40,51 to 37,84 | No | ns |
| | | 0 hpi vs 48 hpi | −1 | −40,18 to 38,18 | No | ns |
| **Δ*sctT*::Kan$^r$** | Morphotype 1 | 0 hpi vs 24 hpi | −7 | −46,18 to 32,18 | No | ns |
| | | 0 hpi vs 48 hpi | 5 | −34,18 to 44,18 | No | ns |
| | Morphotype 2 | 0 hpi vs 24 hpi | 18 | −21,18 to 57,18 | No | ns |
| | | 0 hpi vs 48 hpi | 12,67 | −26,51 to 51,84 | No | ns |
| | Morphotype 3 | 0 hpi vs 24 hpi | −10,67 | −49,84 to 28,51 | No | ns |
| | | 0 hpi vs 48 hpi | −16,33 | −55,51 to 22,84 | No | ns |
| | Morphotype 4 | 0 hpi vs 24 hpi | −0,6667 | −39,84 to 38,51 | No | ns |
| | | 0 hpi vs 48 hpi | −1,333 | −40,51 to 37,84 | No | ns |

Abbreviations: SS: sum of squares, df: degrees of freedom, MS: mean square, mean Diff.: mean difference, 95% CI: 95% confidence interval, ns: not significant, * *p°<°0.0332*, ** *p°<°0.0021*, *** *p°<°0.0002*.

**Supplementary Table 9.** Approximate probabilities (p) of Brown-Forsythe test, one-way analysis of variance (ANOVA), and Tukey HSD Post Hoc Test for the reinfection efficiency of *Rhizopus microsporus* with the following *Burkholderia rhizoxinica* transcription-activator like effector (BAT) mutant strains: **a,** Δ*bat1*::Apra$^r$, **b,** Δ*bat2*::Kan$^r$, or **c,** Δ*bat3*::Kan$^r$. The number of endohyphal bacteria in each morphotype (indicated by green fluorescence) 24 hours post infection (hpi) was compared the number of endohyphal bacteria 48 hours after co-cultivation (24 hpi vs 48 hpi). Homogeneous data (non-significant Brown-Forsythe) is shown in black numbers and non-homogeneous data (significant Brown-Forsythe) is highlighted in red numbers. P-values with *p<0.05* were considered statistically significant (highlighted in grey).

**a, Δ*bat1*::Apra$^r$**

| **Brown-Forsythe Test** | |
|---|---|
| F (DFn, DFd) | 0.7611 (7, 16) |
| P value | 0.6269 |
| P value summary | ns |
| Are SDs significantly different (P < 0.05)? | No |

| **ANOVA Table** | **SS** | **DF** | **MS** | **F (DFn, DFd)** | **P value** |
|---|---|---|---|---|---|
| Treatment (between columns) | 77257 | 7 | 11037 | F (7, 16) = 39.96 | P<0.0001 |
| Residual (within columns) | 4419 | 16 | 276.2 | | |
| Total | 81675 | 23 | | | |

**Tukey HSD Post Hoc**

| **Morphotype** | **Time Comparison** | **Mean Diff.** | **95% CI** | **P<0.05?** | **Summary** |
|---|---|---|---|---|---|
| Morphotype 1 | 24 hpi vs 48 hpi | 4 | −45.33 to 53.33 | No | ns |
| Morphotype 2 | 24 hpi vs 48 hpi | 11.6 | −37.73 to 60.93 | No | ns |
| Morphotype 3 | 24 hpi vs 48 hpi | −60.9 | −113.2 to −8.582 | Yes | * |
| Morphotype 4 | 24 hpi vs 48 hpi | −127.6 | −176.9 to −78.27 | Yes | *** |

**b, Δ*bat2*::Kan$^r$**

| **Brown-Forsythe Test** | |
|---|---|
| F (DFn, DFd) | 0.5561 (7, 24) |
| P value | 0.7834 |
| P value summary | ns |
| Are SDs significantly different (P < 0.05)? | No |

| **ANOVA Table** | **SS** | **DF** | **MS** | **F (DFn, DFd)** | **P value** |
|---|---|---|---|---|---|
| Treatment (between columns) | 16372 | 7 | 2339 | F (7, 24) = 33.65 | P<0.0001 |
| Residual (within columns) | 1668 | 24 | 69.5 | | |
| Total | 18040 | 31 | | | |

**Tukey HSD Post Hoc**

| **Morphotype** | **Time Comparison** | **Mean Diff.** | **95% CI** | **P<0.05?** | **Summary** |
|---|---|---|---|---|---|
| Morphotype 1 | 24 hpi vs 48 hpi | 4.2 | −13.26 to 21.66 | No | ns |
| Morphotype 2 | 24 hpi vs 48 hpi | 0.6 | −16.86 to 18.06 | No | ns |
| Morphotype 3 | 24 hpi vs 48 hpi | −0.6667 | −23.21 to 21.88 | No | ns |
| Morphotype 4 | 24 hpi vs 48 hpi | −67.67 | −90.21 to −45.12 | Yes | *** |

**c, Δ*bat3*::Kan$^r$**

| **Brown-Forsythe Test** | |
|---|---|
| F (DFn, DFd) | 1.366 (7, 16) |
| P value | 0.2847 |
| P value summary | ns |
| Are SDs significantly different (P < 0.05)? | No |

| **ANOVA Table** | **SS** | **DF** | **MS** | **F (DFn, DFd)** | **P value** |
|---|---|---|---|---|---|
| Treatment (between columns) | 17834 | 7 | 2548 | F (7, 16) = 1.598 | P=0.2067 |
| Residual (within columns) | 25516 | 16 | 1595 | | |
| Total | 43350 | 23 | | | |

**Tukey HSD Post Hoc**

| **Morphotype** | **Time Comparison** | **Mean Diff.** | **95% CI** | **P<0.05?** | **Summary** |
|---|---|---|---|---|---|
| Morphotype 1 | 24 hpi vs 48 hpi | 15.25 | −77.06 to 107.6 | No | ns |
| Morphotype 2 | 24 hpi vs 48 hpi | −5 | −97.31 to 87.31 | No | ns |
| Morphotype 3 | 24 hpi vs 48 hpi | −7.333 | −113.9 to 99.26 | No | ns |
| Morphotype 4 | 24 hpi vs 48 hpi | −123.4 | −223.1 to −23.71 | Yes | ** |

Abbreviations: SS: sum of squares, df: degrees of freedom, MS: mean square, mean Diff.: mean difference, 95% CI: 95% confidence interval, ns: not significant, * *p°<°0.0332*, ** *p°<°0.0021*, *** *p°<°0.0002*.

**Supplementary Table 10.** Approximate probabilities (p) of Brown-Forsythe test, one-way analysis of variance (ANOVA), and Tukey HSD Post Hoc Test for the formation of septa following reinfection of *Rhizopus microsporus* with *Burkholderia rhizoxinica* transcription-activator like effector (BAT) mutant strains (Δ*bat1*::Apra$^r$, Δ*bat2*::Kan$^r$, or Δ*bat3*::Kan$^r$). **a,** Comparison of the number of septa formed per mycelium area in percent of the positive control (*R. microsporus* ATCC62417/S co-incubated with wild-type *B. rhizoxinica*; % of B1WT) after 24 hours and 48 hours of co-cultivation (24 hpi vs 48 hpi). **b,** Comparison of the number of septa formed per cell number in percent of the positive control (% of B1WT) after 24 hpi and 48 hpi. Homogeneous data (non-significant Brown-Forsythe) is shown in black numbers and non-homogeneous data (significant Brown-Forsythe) is highlighted in red numbers. P-values with *p<0.05* were considered statistically significant (highlighted in grey).

**a**

| **Brown-Forsythe Test** | |
|---|---|
| F (DFn, DFd) | 1.346 (11, 24) |
| P value | 0.2601 |
| P value summary | ns |
| Are SDs significantly different (P < 0.05)? | No |

| **ANOVA Table** | **SS** | **DF** | **MS** | **F (DFn, DFd)** | **P value** |
|---|---|---|---|---|---|
| Treatment (between columns) | 583462 | 11 | 53042 | F (11, 24) = 1.662 | P=0.1440 |
| Residual (within columns) | 765831 | 24 | 31910 | | |
| Total | 1349293 | 35 | | | |

**Tukey HSD Post Hoc**

| **Strain** | **Time Comparison** | **Mean Diff.** | **95% CI** | **P<0.05?** | **Summary** |
|---|---|---|---|---|---|
| Δ*bat1*::Apra$^r$ | 24 hpi vs 48 hpi | −126 | −651.9 to 399.9 | No | ns |
| Δ*bat2*::Kan$^r$ | 24 hpi vs 48 hpi | −324.7 | −850.6 to 201.2 | No | ns |
| Δ*bat3*::Kan$^r$ | 24 hpi vs 48 hpi | −128 | −653.9 to 397.9 | No | ns |

**b**

| **Brown-Forsythe Test** | |
|---|---|
| F (DFn, DFd) | 1.785 (11, 24) |
| P value | 0.1138 |
| P value summary | ns |
| Are SDs significantly different (P < 0.05)? | No |

| **ANOVA Table** | **SS** | **DF** | **MS** | **F (DFn, DFd)** | **P value** |
|---|---|---|---|---|---|
| Treatment (between columns) | 5740573 | 11 | 521870 | F (11, 24) = 57.90 | P<0.0001 |
| Residual (within columns) | 216309 | 24 | 9013 | | |
| Total | 5956882 | 35 | | | |

**Tukey HSD Post Hoc**

| **Strain** | **Time Comparison** | **Mean Diff.** | **95% CI** | **P<0.05?** | **Summary** |
|---|---|---|---|---|---|
| Δ*bat1*::Apra$^r$ | 24 hpi vs 48 hpi | −1427 | −1706 to −1148 | Yes | *** |
| Δ*bat2*::Kan$^r$ | 24 hpi vs 48 hpi | −197.3 | −476.8 to 82.16 | No | ns |
| Δ*bat3*::Kan$^r$ | 24 hpi vs 48 hpi | −127.7 | −407.2 to 151.8 | No | ns |

Abbreviations: SS: sum of squares, df: degrees of freedom, MS: mean square, mean Diff.: mean difference, 95% CI: 95% confidence interval, ns: not significant, * *p°<°0.0332*, ** *p°<°0.0021*, *** *p°<°0.0002*.

**Supplementary Table 11. a,** Approximate probabilities (p) of Brown-Forsythe test and **b,** one-way analysis of variance (ANOVA) for the ratio of live vs dead endohyphal bacteria following reinfection of *Rhizopus microsporus* with *Burkholderia rhizoxinica* wild-type (B1 WT) or *B. rhizoxinica* transcription-activator like effector (BAT) mutant strains (Δ*bat1*::Apra$^r$, Δ*bat2*::Kan$^r$, or Δ*bat3*::Kan$^r$). Homogeneous data (non-significant Brown-Forsythe) is shown in black numbers and non-homogeneous data (significant Brown-Forsythe) is highlighted in red numbers. **c,** The ratio of live/dead bacteria inside the fungal hyphae was compared between *B. rhizoxinica* wild-type (B1 WT) and each of the BAT-deficient mutants for each individual morphotype 48 hours post infection using the Tukey HSD Post Hoc Test. P-values with $p<0.05$ were considered statistically significant (highlighted in grey).

**a**

| **Brown-Forsythe Test** | |
|---|---|
| F (DFn, DFd) | 0.7152 (15, 64) |
| P value | 0.7602 |
| P value summary | ns |
| Are SDs significantly different (P < 0.05)? | No |

**b**

| **ANOVA Table** | **SS** | **DF** | **MS** | **F (DFn, DFd)** | **P value** |
|---|---|---|---|---|---|
| Treatment (between columns) | 189.5 | 15 | 12.63 | F (15, 64) = 0.7350 | P=0.7404 |
| Residual (within columns) | 1100 | 64 | 17.19 | | |
| Total | 1289 | 79 | | | |

**c**

**Tukey HSD Post Hoc**

| **Morphotype** | **Strain Comparison** | | | **Mean Diff.** | **95% CI** | **P<0.05?** | **Summary** |
|---|---|---|---|---|---|---|---|
| Morphotype 1 | B1 WT | vs | Δ*bat1*::Apra$^r$ | −0.075 | −1.032 to 0.8818 | No | ns |
| | | | Δ*bat2*::Kan$^r$ | 0.3 | −0.6568 to 1.257 | No | ns |
| | | | Δ*bat3*::Kan$^r$ | 0.417 | −0.5398 to 1.374 | No | ns |
| Morphotype 2 | B1 WT | vs | Δ*bat1*::Apra$^r$ | 0.1269 | −0.8299 to 1.084 | No | ns |
| | | | Δ*bat2*::Kan$^r$ | 0.251 | −0.7058 to 1.208 | No | ns |
| | | | Δ*bat3*::Kan$^r$ | 0.3043 | −0.6525 to 1.261 | No | ns |
| Morphotype 3 | B1 WT | vs | Δ*bat1*::Apra$^r$ | −0.7021 | −1.659 to 0.2547 | No | ns |
| | | | Δ*bat2*::Kan$^r$ | −0.2131 | −1.170 to 0.7437 | No | ns |
| | | | Δ*bat3*::Kan$^r$ | 0.03581 | −0.9210 to 0.9926 | No | ns |
| Morphotype 4 | B1 WT | vs | Δ*bat1*::Apra$^r$ | 1.087 | 0.1306 to 2.044 | Yes | * |
| | | | Δ*bat2*::Kan$^r$ | 0.5655 | −0.3914 to 1.522 | No | ns |
| | | | Δ*bat3*::Kan$^r$ | 1.302 | 0.3454 to 2.259 | Yes | ** |

Abbreviations: SS: sum of squares, df: degrees of freedom, MS: mean square, mean Diff.: mean difference, 95% CI: 95% confidence interval, ns: not significant, * *p°<°0.0332*, ** *p°<°0.0021*, *** *p°<°0.0002*.

**Supplementary Table 12.** Approximate probabilities (p) of Brown-Forsythe test, one-way analysis of variance (ANOVA), and Tukey HSD Post Hoc Test for the reinfection efficiency of *Rhizopus microsporus* with the following *Burkholderia rhizoxinica* type 3 secretion system mutant strains: **a,** Δ*sctC*::Kan$^r$ or **b,** Δ*sctT*::Kan$^r$. The number of endohyphal bacteria in each morphotype (indicated by green fluorescence) 24 hours post infection (hpi) was compared the number of endohyphal bacteria 48 hours after co-cultivation (24 hpi vs 48 hpi). Homogeneous data (non-significant Brown-Forsythe) is shown in black numbers and non-homogeneous data (significant Brown-Forsythe) is highlighted in red numbers. P-values with *p<0.05* were considered statistically significant (highlighted in grey).

**a, Δ*sctC*::Kan$^r$**

| **Brown-Forsythe Test** | |
|---|---|
| F (DFn, DFd) | 2.641 (7, 16) |
| P value | 0.0511 |
| P value summary | ns |
| Are SDs significantly different (P < 0.05)? | No |

| **ANOVA Table** | **SS** | **DF** | **MS** | **F (DFn, DFd)** | **P value** |
|---|---|---|---|---|---|
| Treatment (between columns) | 1.7 | 7 | 0.2429 | F (7, 16) = 3.675 | P=0.0147 |
| Residual (within columns) | 1.057 | 16 | 0.06607 | | |
| Total | 2.757 | 23 | | | |

**Tukey HSD Post Hoc**

| **Morphotype** | **Time Comparison** | **Mean Diff.** | **95% CI** | **P<0.05?** | **Summary** |
|---|---|---|---|---|---|
| Morphotype 1 | 24 hpi vs 48 hpi | 0 | −0.7266 to 0.7266 | No | ns |
| Morphotype 2 | 24 hpi vs 48 hpi | 0 | −0.7266 to 0.7266 | No | ns |
| Morphotype 3 | 24 hpi vs 48 hpi | 0 | −0.7266 to 0.7266 | No | ns |
| Morphotype 4 | 24 hpi vs 48 hpi | 0.8047 | 0.07810 to 1.531 | Yes | * |

**b, Δ*sctT*::Kan$^r$**

| **Brown-Forsythe Test** | |
|---|---|
| F (DFn, DFd) | 0.9973 (7, 16) |
| P value | 0.4679 |
| P value summary | ns |
| Are SDs significantly different (P < 0.05)? | No |

| **ANOVA Table** | **SS** | **DF** | **MS** | **F (DFn, DFd)** | **P value** |
|---|---|---|---|---|---|
| Treatment (between columns) | 346 | 7 | 49.42 | F (7, 16) = 0.9976 | P=0.4677 |
| Residual (within columns) | 792.7 | 16 | 49.54 | | |
| Total | 1139 | 23 | | | |

**Tukey HSD Post Hoc**

| **Morphotype** | **Time Comparison** | **Mean Diff.** | **95% CI** | **P<0.05?** | **Summary** |
|---|---|---|---|---|---|
| Morphotype 1 | 24 hpi vs 48 hpi | −2.667 | −22.57 to 17.23 | No | ns |
| Morphotype 2 | 24 hpi vs 48 hpi | 0 | −19.90 to 19.90 | No | ns |
| Morphotype 3 | 24 hpi vs 48 hpi | −0.3333 | −20.23 to 19.57 | No | ns |
| Morphotype 4 | 24 hpi vs 48 hpi | 11.33 | −8.567 to 31.23 | No | ns |

Abbreviations: SS: sum of squares, df: degrees of freedom, MS: mean square, mean Diff.: mean difference, 95% CI: 95% confidence interval, ns: not significant, * *p°<°0.0332*, ** *p°<°0.0021*, *** *p°<°0.0002*.

**Supplementary Table 13.** Approximate probabilities (p) of Brown-Forsythe test, one-way analysis of variance (ANOVA), and Tukey HSD Post Hoc Test for the reinfection of *Rhizopus microsporus* with a rhizoxin-deficient *Burkholderia rhizoxinica* mutant strain (Δ*rhzG*::Kan$^r$). **a,** The number of endohyphal bacteria (indicated by green fluorescence) 24 hours post infection (hpi) was compared the number of endohyphal bacteria 48 hours after co-cultivation (24 hpi vs 48 hpi) for each individual morphotype and for all morphotypes combined (Total). **b,** Comparison of the number of septa formed per mycelium area in percent of the positive control (*R. microsporus* ATCC62417/S co-incubated with wild-type *B. rhizoxinica*; % of B1WT) after 24 hours and 48 hours of co-cultivation (24 hpi vs 48 hpi). **c,** Comparison of the number of septa formed per cell number in percent of the positive control (% of B1WT) after 24 hpi and 48 hpi. Homogeneous data (non-significant Brown-Forsythe) is shown in black numbers and non-homogeneous data (significant Brown-Forsythe) is highlighted in red numbers. P-values with *p<0.05* were considered statistically significant (highlighted in grey).

**a, Reinfection with Δ*rhzG*::Kan$^r$**

| **Brown-Forsythe Test** | |
|---|---|
| F (DFn, DFd) | 1.261 (9, 38) |
| P value | 0.2893 |
| P value summary | ns |
| Are SDs significantly different (P < 0.05)? | No |

| **ANOVA Table** | **SS** | **DF** | **MS** | **F (DFn, DFd)** | **P value** |
|---|---|---|---|---|---|
| Treatment (between columns) | 61587 | 9 | 6843 | F (9, 38) = 1.611 | P=0.1472 |
| Residual (within columns) | 161458 | 38 | 4249 | | |
| Total | 223045 | 47 | | | |

**Tukey HSD Post Hoc**

| **Morphotype** | **Time Comparison** | **Mean Diff.** | **95% CI** | **P<0.05?** | **Summary** |
|---|---|---|---|---|---|
| Morphotype 1 | 24 hpi vs 48 hpi | 8 | −170.9 to 186.9 | No | ns |
| Morphotype 2 | 24 hpi vs 48 hpi | 0.6667 | −178.2 to 179.5 | No | ns |
| Morphotype 3 | 24 hpi vs 48 hpi | −123 | −301.9 to 55.87 | No | ns |
| Morphotype 4 | 24 hpi vs 48 hpi | −100 | −278.9 to 78.87 | No | ns |
| Total | 24 hpi vs 48 hpi | −34.17 | −123.6 to 55.27 | No | ns |

**b, Septa per area**

| **Brown-Forsythe Test** | |
|---|---|
| F (DFn, DFd) | 1.346 (11, 24) |
| P value | 0.2601 |
| P value summary | ns |
| Are SDs significantly different (P < 0.05)? | No |

| **ANOVA Table** | **SS** | **DF** | **MS** | **F (DFn, DFd)** | **P value** |
|---|---|---|---|---|---|
| Treatment (between columns) | 583462 | 11 | 53042 | F (11, 24) = 1.662 | P=0.1440 |
| Residual (within columns) | 765831 | 24 | 31910 | | |
| Total | 1349293 | 35 | | | |

**Tukey HSD Post Hoc**

| **Strain** | **Time Comparison** | **Mean Diff.** | **95% CI** | **P<0.05?** | **Summary** |
|---|---|---|---|---|---|
| Δ*rhzG*::Kan$^r$ | 24 hpi vs 48 hpi | −14.33 | −540.2 to 511.6 | No | ns |

**c, Septa per cell number**

| Brown-Forsythe Test | |
|---|---|
| F (DFn, DFd) | 1.785 (11, 24) |
| P value | 0.1138 |
| P value summary | ns |
| Are SDs significantly different (P < 0.05)? | No |

| ANOVA Table | SS | DF | MS | F (DFn, DFd) | P value |
|---|---|---|---|---|---|
| Treatment (between columns) | 5740573 | 11 | 521870 | F (11, 24) = 57.90 | P<0.0001 |
| Residual (within columns) | 216309 | 24 | 9013 | | |
| Total | 5956882 | 35 | | | |

**Tukey HSD Post Hoc**

| Strain | Time Comparison | Mean Diff. | 95% CI | P<0.05? | Summary |
|---|---|---|---|---|---|
| Δ*rhzG*::$Kan^r$ | 24 hpi vs 48 hpi | −42 | −321.5 to 237.5 | No | ns |

Abbreviations: SS: sum of squares, df: degrees of freedom, MS: mean square, mean Diff.: mean difference, 95% CI: 95% confidence interval, ns: not significant, * *p°<°0.0332*, ** *p°<°0.0021*, *** *p°<°0.0002*.

**Supplementary Table 14.** Primers used in this study.

| Gene | Type | Name | Oligo sequence (5' -> 3') |
|---|---|---|---|
| RBRH_03012 (*awr*) | Vector construction | Awr_ArmA_Fw | ctatagggcgaattgggtacgttgatccatgatgtagacgg |
| | | Awr_ArmA_Rv | acattcatccctggctgagaatgtcaacgg |
| | | Awr_ArmB_Fw | gttcttctgaaaatttagtcgacaagcgcg |
| | | Awr_ArmB_Rv | ctcgagggggggcccggtacaacgttgccaagcggataaaatg |
| | | Kan_Awr_Fw | tctcagccagggatgaatgtcagctactgg |
| | | Kan_Awr_Rv | gactaaattttcagaagaactcgtcaagaag |
| | Control Primer | Awr_CA_fw | aatactgctggaggccaatg |
| | | Awr_C_rv | gcttaccacgaagatggacg |
| | | Awr_D_fw | gttcaccaccgcttcgaaac |
| | | Awr_DB_rv | aattgtcgaccgaaatcagc |
| | | Kan_B_Fw | agtgacaacgtcgagcacag |
| | | Kan_A_Rv | cgttggctacccgtgatatt |
| RBRH_01844 (*bat1*) | Vector construction | Tal1844_ArmA_Fw (SpeI) | gatcactagtctgcctagttccgactcgac |
| | | Tal1844_ArmA_Rv | tccccggaatgaattcactactgggtcaccacggcgctgagc |
| | | Tal1844_ArmB_Fw | gattggctgagaattcagtagtagtgggcggttaccagc |
| | | Tal1844_ArmB_Rv (KpnI) | ctagggtaccatgccagttcaccgtctacc |
| | | Apra_Fw_lg | agtagtgaattcattccggggatccgtcg |
| | | Apra_Rv_EcoRI | actactgaattctcagccaatcgactgg |
| | Control Primer | 84_CA_Fw | cgttctcttcgtcatcgtca |
| | | 84_C_Rv | ggccacatgtacgatctcct |
| | | 84_D_Fw | caacacggcttcaatctgg |
| | | 84_DB_Rv | cgctttatgcgatgtgctta |
| | | Apra_A_Rv | tgcttcggggtcattatagc |
| | | Apra_B_Fw | tcggtcagcttctcaacctt |
| RBRH_01776 (*bat2*) | Vector construction | Tal1776_ArmA_Fw (SpeI) | gatcactagtgccggaccgatattgtaaag |
| | | Tal1776_ArmA_Rv | cggaatgaattcactactgggtcaaaccgatgccgaac |
| | | Tal1776_ArmB_Fw | gctgagaattcagtagtaggcagcggttaccagcattg |
| | | Tal1776_ArmB_Rv (KpnI) | ctagggtaccagtgctgcgtcatagtcgtg |
| | | Kan_Tal_Fw | agtagtgaattcattccggggatcagaagaactcgtcaagaaggc |
| | | Kan_Tal_Rv | actactgaattctcagccaatcggatgaatgtcagctactgggc |

| | | | |
|---|---|---|---|
| | **Control Primer** | 77_D_Fw | taacgagcgcgactacagtg |
| | | 76_C_Rv | gttcgagtgggctgaggtta |
| | | 76_D_Fw | caatcgagcgagtatcgtca |
| | | 76_DB_Rv | cacgccatatgtcatcgttc |
| | | Kan_B_Fw | cgttggctacccgtgatatt |
| | | Kan_A_Rv | agtgacaacgtcgagcacag |
| **RBRH_01777 (*bat3*)** | **Vector construction** | Tal1777_ArmA_Fw(SpeI) | gatcactagtcctgcagcgccttactattc |
| | | Tal1777_ArmA_Rv | cggaatgaattcactactaatcccactgacgttgggtcattc |
| | | Tal1777_ArmB_Fw | gctgagaattcagtagtcgaaagatggctgcacaattgtc |
| | | Tal1777_ArmB_Rv (KpnI) | ctagggtacccaacgtcaagcaccgaatac |
| | | Kan_Tal_Fw | agtagtgaattcattccggggatcagaagaactcgtcaagaaggc |
| | | Kan_Tal_Rv | actactgaattctcagccaatcggatgaatgtcagctactgggc |
| | **Control Primer** | 77_CA_Fw | cgttgatgtcggttcaacac |
| | | 77_C_Rv | gccattgttatcggcaatct |
| | | 77_D_Fw | taacgagcgcgactacagtg |
| | | 77_DB_Rv | ctggtaagctgagcccaaac |
| | | Kan_B_Fw | cgttggctacccgtgatatt |
| | | Kan_A_Rv | agtgacaacgtcgagcacag |
| **RBRH_01776_RBRH_01777 (*bat2_bat3*)** | **Vector construction** | TAL1777_ArmA_Fw (SpeI) | gatcactagtcctgcagcgccttactattc |
| | | TAL1777_ArmA_Rv | cggaatgaattcactactaatcccactgacgttgggtcattc |
| | | TAL1776_ArmB_Fw | gctgagaattcagtagtaggcagcggttaccagcattg |
| | | TAL1776_ArmB_Rv (KpnI) | ctagggtaccatgccagttcaccgtctacc |
| | | Kan_B_Fw | cgttggctacccgtgatatt |
| | | Kan_A_Rv | agtgacaacgtcgagcacag |
| | **Control Primer** | 77_CA_Fw | cgttgatgtcggttcaacac |
| | | 76_DB_Rv | cacgccatatgtcatcgttc |
| | | 77_C_Rv | gccattgttatcggcaatct |
| | | 76_D_Fw | caatcgagcgagtatcgtca |
| | | Kan_B_Fw | cgttggctacccgtgatatt |
| | | Kan_A_Rv | agtgacaacgtcgagcacag |

**Supplementary Figure 1.** Alignment of predicted AWR proteins from endofungal *Burkholderia* species, plant pathogenic *Ralstonia solanaceraum*, and plant-associated *Burkholderia* spp. modified from Sole *et al*. . Sequences obtained in this study are highlighted in bold. GenBank accession numbers are indicated in brackets.

```
                                           10        20        30        40        50        60        70        80        90       100
                                  ....|....|....|....|....|....|....|....|....|....|....|....|....|....|....|....|....|....|....|....|
B._rhizoxinica_B1_(CBW77121)      MCTASPAVEHAGPKSNSEKALGSAARKPAATSRHTAPLDELRSKLGVHLRGKQPPASIGSTMPGRAVGSTMPARAIGVTPADRDTPGYSIDTVGVLGVDE
Burkholderia_sp._B2_(MN840555)    MCTASPAVEHAGPKSNSKKALGSAARKPAATSRHTAPLDELRSKLGVHLRGKQPPASVGSTMPGRAVGSTMPARAIGVTPADRDTPGYSIDTVGVLGVDE
Burkholderia_sp._B3_(MN840550)    VYTASPAVEHASPKSNRKKSLGSTAPKPAATSRRTAALHELRSKIGAPLRGKQPRPSAGAAIP---------ARAIGVTPADYDTPGYSIDTVGVLGVDE
Burkholderia_sp._B4_RBRH_00457    VYTASPAVEHAGPKSNRKKSLGSTAPKPAATSRRTAALHELRSKIGAPLRGKQPRPSVGAAIP---------ARAIGVTPADYDTPGYSIDTIGVLGVDE
B._endofungorum_B5_(MN840553)     VYTASPAGEHAGPKSNHKKSLGFTAPKPAATSRRTAALHELRSKIGAPLRGKQPRPSVGAAIP---------ARAIGVTPADYDTPGYSIDTVGVLGVDE
Burkholderia_sp. B6_(MN840556)    MCTASPAVEHAGPKSNSEKALGSAARKPAATSRHTAPLDELRSKLGVHLRGKQPPASIGSTMPGRAVGSTMPARAIGVTPADRDTPGYSIDTVGVLGVDE
Burkholderia_sp._B7_(MN840551)    VYTASPAVEHAGPKSNRKKSLGSTAPKPAATSRRTAALHELRSKIGAPLRGKQPRSSVGAAIP---------ARAIGVTPADYDTPGYSIDTIGVLGVDE
Burkholderia_sp._B8_(MN840554)    MCTASPAVEHAGPKSNSEKALGTAARKPASTSRHTAPLDELRSKLGIRLRGKQPPASVRSAMP---------ARAIGVTPTDHDTPGYNIDTVGVLGVDE
B._pseudomallei_(ZP04890375)      ETRHLAVSLFTGEEGREALSGWIADFDPGGAARLQRMHQLMYAAAVVLKQNGRAAARLLRNYCG-----------MSTERPDLQLKRQSYSMQRVLVSAR
B._pseudomallei_(ZP02502372)      ETRHLAMSLFTGEEGREALSGWIADFDPGGAARLQRMHQLMYAAAVVLKQNGRAAARLLRNYCG-----------MSTERPDLQLKRQSYSMQRVLVSAR
B._pseudomallei_(ZP02494128)      EARHLAMSLFTGEEGREALSGWIADFDPGGAARLQRMHQLMYAAAVVLKQNGRAAARLLRNYCG-----------MSTERPDLQLKRQSYSMQRVLVSAR
X._euvesicatoria_(WP011347306)    LAEANAEIDRLRSAYAQARDALQSDQAGNSANRPASRTLGSHLAAERNREVEALRTRLGRNENMLTLQRTHAERCRRHARRLQQRLDQVEAKLQQARHSS
X._campestris_pv._musacearum_(    LARTNAQIEQLQSAYAQARDALQSEQASSSASAPAPRTLASHLAAEQRQEMQVLRTQLVRDEQMLSLQRVHAERCQRNARRLQERLAQVEAKLQQAQHSS
X._vasicola_pv._vasculorum_(AV    LARTNAQIEQLQSAYAQARDALQSEQASSSASAPAPRTLASHLAAEQRQEMQVLRTQLVRDEQMLSLQRVHAERCQRNARRLQERLAQVEAKLQQAQHSS
X._citri_(ARR12757)               LADANAEIDQLRSAYAQARDALQSDQASSSANRPAPRTLGSHLAAERNREIEALRTRLGRDENMLTLQRTHAERCQRNARRLQERLAQVDAKLQQARHSS
X._campestris_(AKS22319)          LADTNAEIDQLRSAYAQARDALRSDPASSSANRPASRTLASELVAEQDREIEALRTRLGYDEKMLTLQGTHAERCRRNTRRLQARLAQVDAKLEQARHSS
X._oryzae_(AKO19890)              LARTNAQIDQLQSAYAQARDALQSDQASSSASAPASRTLASHLAAEQSQEIQALRTQLARDEQMLSLQSVHAERCQRNARRLQERLVQVEATLKQAQHSS
X._oryzae_(ACD59124)              LARTNAQIDQLQSAYAQARDALQSDQASSSTSAPASRTLASHLAAEQSQEIQALRTQLARDEQMLSLQSVHAERCQRNARRLQERLVQVEATLKQAQHSS
Burkholderia_sp._(WP013592484)    -------GAKLAHASIERTAAFIAEGSAARGSLPMPKFMNPRAIRDTLRRLRNREANAAAREWGQTLDDHFMQPVIAVNRLFPIPADASREAADARQAAI
Burkholderia_sp._(ADN59567)       EIAIGAHGAKLADASIARTTASLAAGDATRGALPVPKFMNPRAIRDTLRRLRNREANATAREWADTLDEHFMQPVIAVNRLFPVPANASREVADARQAAI
R._solanacearum_(YP003748288)     MSATKP-RPEVLQSRPVPAGTRPEDLPGVPDLKAPQTAARQRKIDATAKAAFNPPTSIAESMRGHLGLGRTKVAPQSVSIAEAMSSEAGLQVHLARLIGQ
R._solanacearum_(BAH04968)        ASKHSGRPVNTVADYSPPARTVSAPPDLPTTADRDRASRSLARTAGLVQRAQRGLQKLQGTRHARAPKPDASSTRLPVTVPDSAPRYRQLLAQGFAATQD
R._solanacearum_(BAD42384)        ASKHSGRPVNTVADYSPPARTVSAPPDLPTTADRDRASRSLARTAGLVQRAQRGLQKLQGTRHARAPKPEASSTRLPVTVPDSAPRYRQLLAQGFAATQD
R._solanacearum_(YP003747289)     AAINVTSVPGFTKTTIDPLP--IPKRPPPPSLGKTPRPAGQHKLDAIDRQVDNLTAKLYPPLPTGTRADPEAKRAIGSERAFASAVEALFVPSGASVEAK
R._solanacearum_(WP013209172)     AAISVTAVPGFTKTTIDPLP--VPKRPPPPKLGKVPKPAGQHKLDGIDRQVHNLTEKLYPPLPAGLHSEPDAKRSIAHEWAFAGAVEALFVPSGASAEAA
R._solanacearum_(WP011002069)     IIFPFYSKTGTTPTRAQPKPDVVPALKPAPRRMRPPAAPGRKHQRNVDAVVPPAMATSPRAEIRFPLTTADAQRYLRAQWRQAETGGRFDQAAVDRLTTT
R._solanacearum_(WP003275417)     MATTGPERAKARPSYPIPAR---ADMPPAPPTGPAHTPARQRKIDAAVRATFNPGPSIAELLRHPLGR---KPAAAPVPITEAMGTAEGMQARLGQLFGS
R._solanacearum_(WP003274509)     AAINVTSVPGFTKTTIDPLP--IPKRPPPPSLGKTPRPAGRHKLDAIDRQVDNLTSKLYPPLPTGTRGDPEAKRAIGSQRAFASAVEALFVPWGASVEAK
R._solanacearum_(EAP71656)        MSATRP-RPEVLQSRPVPAGTRPEDLPGVPDLKAPQTAARQRKIDATAKAAFNPPTSIAESMRGHLGLGRTKVTPQSVSIAEAMGSEAGLQVHLARLIGQ
R._solanacearum_(EAP71039)        MATTGPERAKARPSYPIPAR---ADMPPAPPTGPAHTPARQRKIDAAVRATFNPGPSIAELLRHPLGR---KPAAAPVPITEAMGTAEGMQARLGQLFGS
R._solanacearum_(CBJ39728)        AAINVTAVPGFTKTTIDPLP--IPKRPPPPNLSKTPKPAGQHKLDRIDRQVDNLTRKLYPPLPSGQHGDPEAKRAIASGRAFASAVEALFVPSGVGANAV
R._solanacearum_(CBJ35355)        MSATKP-RPEVLQSRPVPAGTRPEDLPSVPATGPAQPAARQRKIDATVKATFNPPSSIAESMRSHLGLGQAKVT-QAVSISEAMSSQAGLQVHLARLLGQ
R._solanacearum_(CAD17997)        ---------NALKAYPPPALSEASALQPATAREQARAARSQAATAALVQRTQQGLQRMRGTDNSHAAASAEPGARQPVTVPESARQYRQLLAQQFAATRD
R._solanacearum_(BAH47286)        IIFPFYSKTGTTPTRAQPKPDVVPALKPAPRRMRPPAAPGRKHQRNVDAVVPPAMATSPRAEIRFPLTTADAQRYLRAQWRQAETGGRFDQAAVDRLTTT
R._solanacearum_(BAH47283)        AAINVTAVPGFTKTTIDPLP--IPKRQPPPNLPKAPKPAGQHKLDRIDRQVHNLTQKLYPPLPSGLHADPEAKRAIASSRAFVSAVEALFVPSGAGVYAV
R._solanacearum_(BAH04967)        ---------NALKAYTPPALSEASALQPATAREQARAARSQAATAALVQRTQQGLQRMRGTDNSHAAASAEPGARQPVTVPESARQYRQLLAQQFASTRD
R._solanacearum_(BAD42389)        MSATRP-RPEVLQSRPVPAGTRPEDLPSVPATGPAQPAARQRKIDATVKAAFNPSSSIAESMRSHLGLGQDKVARQSVSIAEAMSSEAGMRVHLARLLSQ
```

```
                                          110       120       130       140       150       160       170       180       190       200
                                 ....|....|....|....|....|....|....|....|....|....|....|....|....|....|....|....|....|....|....|....|
B._rhizoxinica_B1_(CBW77121)     PRHWTRAQLLNHLVEANYPGPTLATAELAYQLHTSLDDGTGEVNAARVQLVLFAMHEAGLRDNTRAMMD-------------------------------
Burkholderia_sp._B2_(MN840555)   PRHWTRAQLLNHLVEANYPGPTLATAELAYQLHTSLDDGTGEVNAARVQLVLFAMHEAGLRDNTRAMMD-------------------------------
Burkholderia_sp._B3_(MN840550)   PRLWTRAQLLNHLVEANYPGPNLATAELAHQLHTGLDDGTGKVSAARAQLVLFAMHEAGLRD--------------------------------------
Burkholderia_sp._B4_RBRH_00457   PRLWTRAQLLNHLVEANYPGPNLATAELAHQLHTGLDDGTGKVSAARAQLVLFAMHEAGLRD--------------------------------------
B._endofungorum_B5_(MN840553)    PRLWTRAQLLNHLVEANYPGPNLATAELAHQLHTGLDDGTGKVSAARAQLVLFAMHEAGLRD--------------------------------------
Burkholderia_sp. B6_(MN840556)   PRHWTRAQLLNHLVEANYPGPTLATAELAYQLHTSLDDGTGEVNAARVQLVLFAMHEAGLRDNTRAMMD-------------------------------
Burkholderia_sp._B7_(MN840551)   PRLWTRAQLLNHLVEANYPGPNLATAELAHQLHTGLNDGTGKVSAARAQLVLFAMHEAGLRD--------------------------------------
Burkholderia_sp._B8_(MN840554)   PRYWTRAQLLNHLVEANYPGPNLVTAELAHQLHTGLDDGTGKVSAARAQLVLFAMHDAGVRD--------------------------------------
B._pseudomallei_(ZP04890375)     GTSLDILGIPPSITGREHHAEQKSVYRNALAIAQYLHEKLDAAYKPDSLEDMLAAAQIVSAETLGGSGDRTEREYHFSR---------------------
B._pseudomallei_(ZP02502372)     GTSLDILGIPPSITGREHHAEQKSVYRNALAIAQYLHEKLDAAYKPDSLEDMLAAAQIVSAEALGGSGDRTEREYHFSR---------------------
B._pseudomallei_(ZP02494128)     GTSLDILGIPPSITGREHHAEQKSVYRNALAIAQYLHEKLDAAYKPDSLEDMLAAAQIVSAEALGGSGDRTEREYHFSR---------------------
X._euvesicatoria_(WP011347306)   RINTEEVWQHEQTVNALEAANGDMQRVAEHAAAQPPADEARPVAAEIVADTMDALLGALPGFR-------EETIDPVSR---------------------
X._campestris_pv._musacearum_(   RVNTKELWQHEQTVSALEAANGDMQRAVETAAAQPPANEARRAPDDSVADAMAALLDALPGFR-------EDTIGPMSR---------------------
X._vasicola_pv._vasculorum_(AV   RVNTKELWQHEQTVSALEAANGDMQRAVETAAAQPPANEARRAPDDSVADAMAALLDALPGFR-------EDTIGPMSR---------------------
X._citri_(ARR12757)              RINTEELWQYEQTVSALEAANGDMQRVVEHAAAHPPADEALPVAADIVADTMEALLEALPGFR-------EQTIDPVNR---------------------
X._campestris_(AKS22319)         RINTEELWQQEQTVSVLEAADGDMQRGVEHAAAQLPADEARPVASDIVADTMQTLLDVLPGFR-------EETLDPVSR---------------------
X._oryzae_(AKO19890)             RVNTTELWQQDHTVSALEAANGDMQRAVETAAAQSPADEARRAPDDSVAEAMAALLDALPGFR-------EDTIGPMSR---------------------
X._oryzae_(ACD59124)             RVNTKELWQQDHTVSALEAADGDMQRAVETAAAQSPADEARRAPDDSVAEAMAALLDALPGFR-------EDTIGPMSR---------------------
Burkholderia_sp._(WP013592484)   NRLLDGLTLRRPRPARADASLPPAVERLPQACVVPLAQALAEASDGDPILAARILDRLQQPF-------DIAG---------------------------
Burkholderia_sp._(ADN59567)      DSLLDGLTLRRPRPAQADPTLPLTVDRLPQECVVPVADALAEASGGDPALAARILHRLQQPL-------DIGGLSGVSAIGDTSSTRDTAGTAENSGISG
R._solanacearum_(YP003748288)    LVHAPAVHTLHQALLGGLPQGAESASTHPLAQHVLVAEALANACGGDAVRATAALEALKAGT-------FLPTSHAEGE---------------------
R._solanacearum_(BAH04968)       AGAVGRLADRLRALAPPLAEEAG--ERTRYARPMLLAKVLEHTCGQDAAAALRVLDRLTGGM-------RLGAPTFAAPDG-------------------
R._solanacearum_(BAD42384)       AGAVGRLADRLRALAPPLAEEAG--ERTRYARPMLLAKVLEHTCGQDAAAALRVLDRLTGGM-------RLGAPTFAAPDG-------------------
R._solanacearum_(YP003747289)    AETAPHLHNLLRDLTRIAEAQPDAVDSTRYAGMLQCARALAAATGGNAADACAALAHLRTHF-------SLTDGSDNAP---------------------
R._solanacearum_(WP013209172)    SR----LPDLLGHLKQVADAQPGVADATRYAGMLQCARALATATGGNAADACVALEHLRTHF-------SLTDGSGGAP---------------------
R._solanacearum_(WP011002069)    LT-----ARLLATSAATGNGVHEDSGEKACARAIMVCDALAQATGGDPARAHDALLALLDTT-------SHADAR-------------------------
R._solanacearum_(WP003275417)    RAGAPGMQALHTALLGTFAARLVSASARPFARHVLLAEALANASEGDAVRAMAALRVLQSGR-------FLPMLFPENQ---------------------
R._solanacearum_(WP003274509)    AETARHLHNLLRDLTQIAETQPDAADSTRYAGMLQCARALVAATGGNAADACAALGHLRTHF-------SLTDGSDSES---------------------
R._solanacearum_(EAP71656)       PAHAPAVRTLHRALLDRLPRVAESASAQPFAQHVLVTEALANACGGDAVRATAALRALKAGA-------FLPTLHAEGE---------------------
R._solanacearum_(EAP71039)       RAGAPGMQALHTALLGTFAARLVSASARPFARHVLLAEALANASEGDAVRAMAALRVLQSGR-------FLPMSFPENQ---------------------
R._solanacearum_(CBJ39728)       PG----LRNLIHGLARMADAQPDAVDTTRYAGMLQCARALATATGGRADEACRAFEHLREHF-------SLTDGSDGAA---------------------
R._solanacearum_(CBJ35355)       PVHAPTVRAFHDALLDSLPQTTESASERPFAQHVLLAEALANACGGDTVRATAALHALKAGE-------FLPTSHAADE---------------------
R._solanacearum_(CAD17997)       TGAVARLSARLEALAPP-ADAVG--EHLRYARPTLVAEALAHTCGHDAAAALRVLASLENEA-------RLDTAAFGTPDAPARA---------------
R._solanacearum_(BAH47286)       LT-----ARLLATSAATGNGVHEDSGEKACARAIMVSDALAQATGGDPARAHDALLALLDTT-------SHADAR-------------------------
R._solanacearum_(BAH47283)       PG----LQTLIDGLAGMAEAQPDAVDTTRYAGMLQCARALATATGGRADEACRALEHLRDRF-------SLTDGSNGAP---------------------
R._solanacearum_(BAH04967)       TGAVTRLSARLEALAPP-ADAVG--EHLRYARPTLVAEALAHTCGHDAAAALRVLASLENEA-------RLDTAAFGTPDAPARS---------------
R._solanacearum_(BAD42389)       PLHAPAVRALHGALLDSLPQATESASGRGFAQHVLLAEALVNACGGDTIRATAALRALKAGE-------FLPTAHGTGD---------------------
```

```
                                           210       220       230       240       250       260       270       280       290       300
                                  ....|....|....|....|....|....|....|....|....|....|....|....|....|....|....|....|....|....|....|....|
B._rhizoxinica_B1_(CBW77121)      -----------------------------STMATMVLQQLRLLNFDQPFHDAPTPTGDARLPDQQLQAWRASWETARVLARTEEGFETLLKLR-------
Burkholderia_sp._B2_(MN840555)    -----------------------------STMATMVLQQLRVLNFDQPFHDAPTPAGDAHLPDQQLQAWRASWETARVLARTEEGFETLLKLR-------
Burkholderia_sp._B3_(MN840550)    -----------------------------STTATMMLRQLRTLNFDQPFHDASAAAGGAHLPDQR-QAWRASWETARVLARTEYGFETLLKLR-------
Burkholderia_sp._B4_RBRH_00457    -----------------------------STTATMVLRQLRALNFDQPFHDASAAVGGAHLPDQR-QAWRASWETARVLARTEYGFETLLKLR-------
B._endofungorum_B5_(MN840553)     -----------------------------STTATMVLRQLRTLNFDQPFHDASAAAGGAHLPDQQ-QAWRASWETARVLARTEYGFETLLKLR-------
Burkholderia_sp. B6_(MN840556)    -----------------------------STMATMVLQQLRLLNFDQPFHDAPTPTGDARLPDQQLQAWRASWETARVLARTEEGFETLLKLR-------
Burkholderia_sp._B7_(MN840551)    -----------------------------STTATMMLRQLRTLNFDQPFHDASAVAGGAHLPDQR-QAWRASWETARVLARTEYGFETLLKLR-------
Burkholderia_sp._B8_(MN840554)    -----------------------------STTATKVLRQLRALNFDQPFHDVSAAAAGAHLPDQQ-QAWRASWETARVLARTEYGFETLLKLR-------
B._pseudomallei_(ZP04890375)      --------------------------VVDNLGSIYQEERLSIDALACAANLISIDRDNGAKALGAVAMLSSSEQADVMDALEQAGELVRLLPLGIRRDIA
B._pseudomallei_(ZP02502372)      --------------------------VVDNLGSIYQEERLSIDALACAANLISIDRDNGAKALGAVAMLSSSEQADVMDALEQAGELVRLLPLGIRRDIA
B._pseudomallei_(ZP02494128)      --------------------------VVDNLGSIYQEERLSIDALACAANLISIDRDNGAKALGAVAMLSSSEQADVMDALEQAGELVRLLPLGIRRDIA
X._euvesicatoria_(WP011347306)    -------------------------QVLRAWADGLHGRSAAFGGDALQPVDALRIALQALAIASEGAAAAAADVLRRLGMVRLRELIPPPQVSIAAEAAQ
X._campestris_pv._musacearum_(    -------------------------QILRTWAGGLQGRSAAFGTDALQPVDALNIALQALSIASEGDATEAANVLQRLGEVQLRELIPPPQVSNAAEAVP
X._vasicola_pv._vasculorum_(AV    -------------------------QILRTWAGGLQGRSAAFGTDALQPVDALNIALQALSIASEGDATAAANVLQRLGEVQLRELIPPPQVSNAAEAVP
X._citri_(ARR12757)               -------------------------QVLRAWADGLHGRSAAFGSDALQPVDALRIALQALAIASDGDATAAASTLQRLGVVQLRELIPPPQVSIAAEAVP
X._campestris_(AKS22319)          -------------------------QVLRVWADGLHARSAAFGTEALQPVDALRIALQALSIASGGDATAAANALQRLSVVRMRELIPPPQVSIAAEAVP
X._oryzae_(AKO19890)              -------------------------QVLRTWAEGLHRRSAAFGTDALQPVDALNIALQALSIASQSDATTAANVLQRLGEVQLRELIPPPQVSNAAEAVP
X._oryzae_(ACD59124)              -------------------------QVLRTWADGLHRRSAAFGTDALQPVDALNIALQALSIASQSDATTAANVLQRLGEVQLRELIPPPQVSNAAEAVP
Burkholderia_sp._(WP013592484)    ----------------------------DAAHAAPDELERRAWRCAQDLAHARCAALEGLLRLQFGDGARHAERGNGAAARLVVYLQAGRKLR------S
Burkholderia_sp._(ADN59567)       TSDASGTSGTSNTSNTSNTTTIRAAPAQHPSQAAPADLEQRAWRCAQNLAHARCAALEGLLRLQYGDSARHAERGNGMAARLVVYLQAAHKLR------S
R._solanacearum_(YP003748288)     ----------------TASPRPGNGGANGSRIGRMAPDGPGVRRPGNVGANTLHSAPQPAAEDAMLDAMHAAQELAGIGDVGMQALLALMPRLKPAKHEP
R._solanacearum_(BAH04968)        ----------------------------SPDAHEP-ALIRQASAASDAFFDALSDFHDAAPAAETGAAWRTAQLLARYHAGFDALCSLMQIPDVPEQRQA
R._solanacearum_(BAD42384)        ----------------------------SPDAHES-ALIRQASAASDAFFDALSDFHDAAPAAETGAAWRTAQLLARYHAGFDALCSLMHIPDVPEQRQA
R._solanacearum_(YP003747289)     -----------------------PSPEQKQAWSAAKLLAHTASGFDALLALRPALAEVDAARPDDG-MNHRMKREGLRTFLQAADHLAARMPA-----DT
R._solanacearum_(WP013209172)     -----------------------ATPAQKHAWSTAKLLAHTASGFDALLALRPALAEVDARHPDDG-MNGRMKREGLRTFLQASDLLAARMPAGVTPPQT
R._solanacearum_(WP011002069)     --------------------------VRADVLALQTALGMTAIGLDTLLAIAPHVLPQR------GSAAPEIQREALRHALRAADHLLRRKPADAPGPTS
R._solanacearum_(WP003275417)     ----------------PASP-LGAPGFSDTGAGALHPAR-------SSFPDDGMGLSPAQTARAKLDALHVGQALASIGDVGMQTLAASIPGLEPARHEP
R._solanacearum_(WP003274509)     -----------------------PSPEQKHAWSAAKLLAHTASGFDSLLALRPALAEVDAVRPDDG-MNDRMKREGLRTFLQAADHLAARMPA-----DT
R._solanacearum_(EAP71656)        ----------------TASPRSSNAGADASRIARMALDGPGVRRTGSAGADALHSAPQPAAEDAMLDAMHAAQELAGIGDVGMQALLALMPRLKPARHEP
R._solanacearum_(EAP71039)        ----------------PASP-LRAPRFSDTGAGALHPAR-------SSFPDDGMGLSPAQTARAKLDALHVGQALASIGDVGIQTLAASIPGLEPALHEP
R._solanacearum_(CBJ39728)        -----------------------PTPAQTQAWSTAKLLAHTASGFDALLALRPALAKVDATLPDDDHMNGRMKREGLRTFLQAADLMAARLPAGVAPPQT
R._solanacearum_(CBJ35355)        ----------------TASPRTSHASVRIASAAQVAPNGPDTRGIDNADADTSPSTRRVLAKDAMLDAMHAAQELAGIGDVGMQALLALMPPLRPAKREP
R._solanacearum_(CAD17997)        ----------SVSSATFHDAISEQEGPDEPDGPPPQSPSARSSASSEAFFDAMSVLEEPGEPEDAGAAWRTAQLLSRYHAGFEALCDLMRLPQTPAHRHA
R._solanacearum_(BAH47286)        --------------------------VQADVLALQTALGMTAIGLDTLLAIAPHVLPQRGL--AEGSAAPEIQREALRHALRAADHLLRRKPADAPGPTS
R._solanacearum_(BAH47283)        -----------------------PIPAQMHAWSTAKLLAHTASGFDALLALRPALAEVDATLPDDDNMNGRMKREGLRTFLQAADLMAARLPAGVAPPQT
R._solanacearum_(BAH04967)        ----------SVSSATFHDAISEQEAPNEPDGPPPQSPSARSSASSEAFFDAMSVLEEPGEPEDAGAAWRTAQLLSRYHAGFEALCDLMRLPQTPAHRHA
R._solanacearum_(BAD42389)        ----------------AASPRPSDAGAEDAATAQAEPAGQHARRIGNASANAIHPTRQTPVEHAMLDAMRAAQGLAGIGDVGMQALLAVMPPLQPAKREP
```

```
                                             310       320       330       340       350       360       370       380       390       400
                                    ....|....|....|....|....|....|....|....|....|....|....|....|....|....|....|....|....|....|....|....|
B._rhizoxinica_B1_(CBW77121)        AEKCAPLSRSVDRDMLHVLLDSADRIDNHVRLTQPHDS------------HATPVHAAPDFDDPSPSGTDKRQQWLQGHLAMKGQDAKSTLLASQVFAIA
Burkholderia_sp._B2_(MN840555)      AEKCAPLSRSVDRDMLHVLLDSADRIDNHVRLTQPHDS------------HATPVHAAPDFDDPSPSGTDKRQQWLQGHLAMNGQDAKSTLLASQVFAIA
Burkholderia_sp._B3_(MN840550)      AEKCAPLSRSADRDMLHVLLDTAEWIENHVRLAQQRDG------------HATSVHAALDFDDPSPSGTAKRQQWLQSHLAMNGQDAKATLLASQVFAIA
Burkholderia_sp._B4_RBRH_00457      AEKCAQLSRSANRDMLHVLLDTAEWIENHVRLAQQRDG------------HATSVHAALDFDDPSPSGTAKRQQWLQSHLAMNGQDAKATLLASQVFAIA
B._endofungorum_B5_(MN840553)       AEKSAPLSRSADRDMLHVLLDTADRIETHVRLVQQRDG------------HATSVHAALDFDDPSPSGTAKHQQWLQSHLAMNGQDAKATLLASQVFAIA
Burkholderia_sp. B6_(MN840556)      AEKCAPLSRSVDRDMLHVLLDSADRIDNHVRLTQPHDS------------HATPVHAAPDFDDPSPSGTDKRQQWLQGHLAMKGQDAKSTLLASQVFAIA
Burkholderia_sp._B7_(MN840551)      AEKCAPLSRSADRDMLHVLLDTAERIENHVRLAQQRDG------------HATSVHAALDFDDPSPSGTAKRQQWLQSHLAMNGQDAKATLLASQVFAIA
Burkholderia_sp._B8_(MN840554)      AEKFAPLSRSADRDMLHVLLDTAERIENYVRFVQQHDG------------HATPMHAALDFDDSSPSGTAKRQQWLQGHLAMNGQDAKATLLASQVFAIA
B._pseudomallei_(ZP04890375)        TVADLDKVLNANGEGPGARASVSSVDPMGQASAQKAIGKLRAALAVAP-DIGNIPIGAAARAAIPMQQKIAYFKWRNG-LADERKARLVFERLYKLNKYA
B._pseudomallei_(ZP02502372)        TVADLDKVLNANGEGPGARASVSSVDPMGQASAQKAIGKLRAALAVAP-DIGNIPIGAAARAAIPMQQKIAYFKWRNG-LADERKARLVFERLYKLNKYA
B._pseudomallei_(ZP02494128)        TVADLDKVLNANGEGPGARASVSSVDPMGQASAQKAIGKLRAALAVAP-DIGNIPIGAAARAAIPMQQKIAYFKWRNG-LADERKARLVFERLYKLNKYA
X._euvesicatoria_(WP011347306)      PMEIVTACVGLLASVPRGMQVLSHMLREEATPPSREWLEAAQVHLRASHRLALTPDAEQAERSWLGDAQMGARHAVHGDSPLQGLQSATDAQRSAFHAFR
X._campestris_pv._musacearum_(      PMEAVTTCVGLLASVPRGMQVLLHTLCEQPTSPAREWLEAAQVYLRASHRLALTPDAEQAERAWLADAQMGARHAVHGATPLQGLQSATDAQRSAFHAFR
X._vasicola_pv._vasculorum_(AV      PMEAVTTCVGLLASVPRGMQVLLHTLCEQPTSPAREWLEAAQVYLRASHRLALTPDAEQAERAWLADAQMGARHAVHGATPLQGLQSATDAQRSAFHAFR
X._citri_(ARR12757)                 TMETVTACVGLLASVPRGMQVLSHMLREEPTPPSREWLEAAQIHLRASHRLALTPDAEQAERTWLGDAQMGARHAVHGDSPLQGLQSATDAQRGAFHAFR
X._campestris_(AKS22319)            PMETVTACVGLLASVPRGMQVLSHMLREEPTPASREWLEAAQVHLRASHRLALTQDAEQAERTWLGDAQMGARHAVHGASPLQGLQSATDAQRGAFHAFR
X._oryzae_(AKO19890)                PMEAVTTCVGLLVSVPRGMQVLLHMLREEPTAPAREWLEAAQVHLRASQRLALTPDAQQAERAWLANAQMGARHAVHGATPLQGLQSATDAQRSAFHAIR
X._oryzae_(ACD59124)                PMEAVTTCVGLLVSVPRGMQVLLHMLREEPTSPAREWLEAAQVHLRASHRLALTPDAEQAERAWLADAQMGARHAVHGATPLQGLQSATDAQRSAFHAIR
Burkholderia_sp._(WP013592484)      IDADLGATPADLNHESTYRLFADDPYGRHAMLALDAARRLPP--------DAADAHDTG---ACEIEAITAYRLWQLG-FDESGPGSDLARASQLLFDGG
Burkholderia_sp._(ADN59567)         MDADLGATPADLNDDAAYRMFIEDPYARHALLALDAARRLPP--------DAASVHDTG---DCEIEAITAYRLWQMG-FDESGPGSDLERAAQLLFDGG
R._solanacearum_(YP003748288)       LECVLQAAEQVTTETKGAQVRLPDIIGHAEKVADTRNDTLACKVLR---AQVKALRAPDTYDALSRADKSAVFAWRQG-FRTDDRHSLLSQTQQRLAK-F
R._solanacearum_(BAH04968)          AHAFLQAADALQHGSVKPAGSPQALLDKHPIGSGTPEESLAIK-------TLHGAAAVLRGEAPTPEQAGALFAWRQG-FREEGPGTALDKTKARIGR--
R._solanacearum_(BAD42384)          AHAFLQAADALQHGSVKPAGSPQALLDKHPIGSGTPEESLAIK-------TLHGAAAVLRGEAPTPEQAGALFAWRQG-FREEGPGTALDKTKARIGR--
R._solanacearum_(YP003747289)       PGALLQRARSHLASASDRHSLAGDGLAVHALLCAADLHADPG--------KAAHAIGDR-------TQVAAYVAWRSG-YREGGKGSALERSLGRMNK-F
R._solanacearum_(WP013209172)       PHAQWQDAHNRLSHAADRTSLGGDGLVLNALLCAAKVHAAPH--------DAANAIDDR-------TQVAAYVAWRSG-YREGGKGTALERSLGRMNK-F
R._solanacearum_(WP011002069)       LEALAALADPVLPAVERGRPGSSAGASVAPPGKAPAWLADPGNALAIKALHAANALRADPAAQCPPHLAQAYLAWRNG-FDREGPGTDLAKAQQRLFK-L
R._solanacearum_(WP003275417)       LEFVLQSAERITQETRGRKVHLGEILAHADAVRGTSANTLACKVLR---AEVRGLGAEHPVSGLDRADKAAVFAWRQG-FRSDEKRSPLRRTQERFAK-F
R._solanacearum_(WP003274509)       PGALLERARSHLASASDRHSLAGDGLAVHALLCAADLHADPS--------KAAHAIGDR-------TQVAAYVAWRSG-YREGGKGSALERSLGRMNK-F
R._solanacearum_(EAP71656)          LECVLQAAEQITTETKGAQVRFPDIIAHAEKVAGTRDDTLACKVLR---AEVKALRTPGIYDALSRADKSAVFAWRQG-FRTDDRHSLLSQTQQRLAK-F
R._solanacearum_(EAP71039)          LEFVLQSAERITQETRGRKVHLGEILAHADAVRGTSADTLACKVLR---AEVRGLGAEHPVSGLDRADKAAVFAWRQG-FRSDEKRSPLRRTQERFAK-F
R._solanacearum_(CBJ39728)          PHAQLQDARARLVAATDRASLAGDGLALNALLCAAQVHAQPH--------EAAQAVKDP-------AQVAAYVAWRSG-YREGGKGSALERSVGRMHK-F
R._solanacearum_(CBJ35355)          LEFVLQAAEQVAAETKGAQVRLPDIIEHAEKMAGTRGDTLACKVLR---AEVKALRAADTHDALSRADKSAVFAWRQG-FRSDDKHSLLSQTQQRLAK-F
R._solanacearum_(CAD17997)          AHAFLQAADALQRGAAGQAASPQALLQRCPPGSGTPDDSLAIK-------TLHAAAAQLRGEALSPEQTGALFAWRQG-FRAEGPGSDLAKVKARTAK--
R._solanacearum_(BAH47286)          LEALAALADPVLPAVERGRPGSSAGASVAPPGKAPAWLADPGNALAIKALHAANALRADPAAQCPPHLAQAYLAWRNG-FDREGPGTDLAKAQQRLFK-L
R._solanacearum_(BAH47283)          PHAQLQDARARLDAAADRASLAGDGLALNALVCAAQVHAEPH--------EAAHAVDDR-------AQVAAYVAWRSG-YREGGKGSALERSLGRMHK-F
R._solanacearum_(BAH04967)          AHAFLQAADALQRGAA----SPRALLQRCPPGSGAPDDSLAIK-------TLHAAAAQLRGEALSPEQTGALFAWRQG-FRAEGPGSDLAKVKARTAK--
R._solanacearum_(BAD42389)          LEFVLQAAEQVSVETKGAQVLLPDIIGHAEKMAGTRNDTLACKVLR---AEVKALRAADTYDALSRADKSAVFAWRQG-FLTDDKHSLLSQTQQRLAK-F
```

```
                                          410       420       430       440       450       460       470       480       490       500
                                 ....|....|....|....|....|....|....|....|....|....|....|....|....|....|....|....|....|....|....|....|
B._rhizoxinica_B1_(CBW77121)     KKFFDQT-----------------------EPPHSASGPTPEGRFASFIETLTPNEKEALFDWQQR---FHESGRGTDKAKASGRLNNVRKVVTSIIGKL
Burkholderia_sp._B2_(MN840555)   KKFFDQT-----------------------EPPHSASGPTPEGRFASFIETLTPNEKEALFDWQQR---FHESGRGTDKAKASGRLNNVRKVVTSIIGKL
Burkholderia_sp._B3_(MN840550)   KKYFDQT-----------------------EHPHSTSGSTPEDRFASFIRTLAPSEKEALFDWQQR---FHESGRGTDKAKASGRLNNVRKVVTSIIGKV
Burkholderia_sp._B4_RBRH_00457   KKYFDQT-----------------------EHPHSTSGSTPEDRFASFIRTLAPSEKEALFDWQQR---FHESGRGTDKAKASGRLNNVRKVVTSIIGKV
B._endofungorum_B5_(MN840553)    KKYFDQT-----------------------EPPHSTSGSTPEDRFASFIRTLAPSEKEALFDWQQR---FHESGRGTDKAKASGRLNNVRKVVTSIIGKV
Burkholderia_sp._B6_(MN840556)   KKFFDQT-----------------------EPPHSASGPTPEGRFASFIETLTPNEKEALFDWQQR---FHESGRGTDKAKASGRLNNVRKVVTSIIGKL
Burkholderia_sp._B7_(MN840551)   KKYFDQT-----------------------EHPHSTSGSTPEDRFASFIRTLAPSEKEALFDWQQR---FHESGRGTDKAKASGRLNNVRKVVTSIIGKV
Burkholderia_sp._B8_(MN840554)   KKYFDQT-----------------------EHPHSASGSTPEDRFASFIKTLTPSEKEALFDWRQR---FHESGRGTDKAKASGRLNHFRKVITSIIGKV
B._pseudomallei_(ZP04890375)     DRAIPRG-----------------------KNLTGRLRRIGDDIAGAFGHRKNPVGMLEEGLLLGG---------FLREDVFNEKIAPAFHSLIEKYRES
B._pseudomallei_(ZP02502372)     DRAIPRG-----------------------KNLTGRLRRIGDDIAGAFGHRKNPVGMLEEGLLLGG---------FLREDVFNEKIAPAFHSLIEKYRES
B._pseudomallei_(ZP02494128)     DRAIPRG-----------------------KNLTGRLRRIGDDIAGAFGHRKNPVGMLEEGLLLGG---------FLREDVFNEKIAPAFHSLIEKYRES
X._euvesicatoria_(WP011347306)   NGYETTRAG--------------------SPYARVNACLQMFAQWAVAGSGRRGRRALNPFNALELG--MKVGASEALPTAVRRANDAFREAAGHLTSWL
X._campestris_pv._musacearum_(   NGYETTRAG--------------------SPYARVNASLQMFAQWAVAGGETRGKRALNPFNALKLG--MKVGASEALPTAVRRANDAFREAAGHLTSWL
X._vasicola_pv._vasculorum_(AV   NGYETTRAG--------------------SPYARVNASLQMFAQWAVAGGETRGKRALNPFNALKLG--MKVGASEALPTAVRRANDAFREAAGHLTSWL
X._citri_(ARR12757)              NGYETTGAG--------------------SPYARVNACLQMFAQWAVAGSAPRGKRALNPFNALALG--MKVGASEALPTAERRANDAFREAAGHLTSWL
X._campestris_(AKS22319)         NGYETTRTG--------------------SPYARVNACLQMFAQWAVAGSRRRGNSDLNPFNALALG--MKVGASEALPTAVRRANDAFREAAGHLTSWL
X._oryzae_(AKO19890)             NGYETTRAG--------------------SPYARVNASLQMFAQWAVAGGQTRGKRALNPFNALKLG--MKVGASEALPTAGRRANDAFREAAGHLSNWL
X._oryzae_(ACD59124)             NGYETTRAG--------------------SPYARVNASLQMFAQWAVAGGQTRGKRALNPFNALKLG--MKVGASEALPTAGRRANDAFREAAGHLSNWL
Burkholderia_sp._(WP013592484)   TTWVARGAERQARKDRALGPIDGAPPAHRFAHARARIGLAVHDMPRAFGKNRSPIGVSSLLGADTPFFDTTRGRYDTALIAARDALLDYSHAQLPAAATP
Burkholderia_sp._(ADN59567)      TTWVARGAERQARKDRALGPIDGAPPAQRFAQARARVGLAVHDVPRAFGRNRTPIDTSSLLGADTPFFDTTRGQYDSALIAAREALLDYSNEQLPCAATP
R._solanacearum_(YP003748288)    RKYVSRAETRDKLDRMRHDP---------SQSTAATVRLVANNVQRAFSHKKSPLLGMGRYGALG----TGTRHVPTDLLELDKHLRTAIDELKGHVAAR
R._solanacearum_(BAH04968)       --FVRR------------------------VIPRVEKHGWRTMLWQMIGKKASPLSSAR-LGLQA----AHRKNLAGEYKDYVDALRSAATTLAARYAQA
R._solanacearum_(BAD42384)       --FVRR------------------------VIPRVEKHGWRTMLWQMIGKKASPLSSAR-LGLQA----AHRKNLAGEYKDYVDALRSAVTTLAARYAQA
R._solanacearum_(YP003747289)    TAWARR------------------------AEHRANHRLAAFDPRRLLGMRKTPVIAAA-YGTGG----ANLGLLNGESKALYDTVQEGIEAMLAHSVVL
R._solanacearum_(WP013209172)    TTWARR------------------------AEHRANHRLAAFDPRRLLGMRKSPLTAAA-YGTGG----ANLGLLNREAQALHDTVKDGLEAMLAHSTVL
R._solanacearum_(WP011002069)    FAYAER------------------------AARTG---MAARAAAGFFGTQKSPLSALQNFGTAG----VMLGHPDDEFARFTAALAP---VKTQLADRL
R._solanacearum_(WP003275417)    RKYVARAEKRDALNKMHFDPRNPI--RSAGKLATKTAGLVTNNVQRVFSHQKSPLLGLGKYGALA----AGAQLAPTDQVRLDEHLITAMGELKAYLESS
R._solanacearum_(WP003274509)    TAWARR------------------------AEHRANHRLAAFDPRRLLGMRKTPVIAAA-YGTGG----ANLGLLNGESKALYDTVQEGIEAMLAHSVVL
R._solanacearum_(EAP71656)       RKYVSRAETRDKLDRMRHDP---------SQSTAATARLVANNVQRAFSHKKSPLLAMGRYGTLG----TGTRHVPTDLLELDKHLRTAIDALKAHVEAR
R._solanacearum_(EAP71039)       RKYVARAEKRDALNKMHFDPRNPI--RSAGKLATKTAGLVTNNVQRVFSHQKSPLLGLGKYGALA----AGAQLAPTDQVRLDEHLITAMGELNAYLKSS
R._solanacearum_(CBJ39728)       TTWARR------------------------AEHRATHPLAAFDPRRLLGMRKSPLTAAA-YGTGG----ANLGLLNQEAQALHDTVKRGIEAMQAHSTEL
R._solanacearum_(CBJ35355)       RKYVSRAETRDKLDKARHDPGN------PMQSSVATARLVANNAQRVFSHKKSPLLAMGKYGSLA----AGTKHVPMDQVELDKHLRTAIDALRVRLEAQ
R._solanacearum_(CAD17997)       --FVSR------------------------TIRRVENHGWRSTLWQVVGKKKSPLSAAR-LGMQA----AHRVSLDKEYTAYAASLRTALGMLGARYREA
R._solanacearum_(BAH47286)       FTYAER------------------------AARTG---MAARAAAGFFGTQKSPLSALQNFGTAG----VMLGHPDDEFARFTAALAP---VKTQLADRL
R._solanacearum_(BAH47283)       TTWARR------------------------AEHRATHPLAAFDPRRLLGMRKSPLTAAA-YGTGG----ANLGLLNQEAQALHDTVKHGIEAMQAHSTAL
R._solanacearum_(BAH04967)       --FVNR------------------------TIRRVENHGWRSTLWQVVGKKKSPLSAAR-LGMQA----AHRVSLAKEYTAYAASLRTALGMLGARYREA
R._solanacearum_(BAD42389)       RKYVSRAETRNALNEARQDP---------ARSTAATARLVANNVQRAFSRKKSPLLAMGKYGSLA----AGTRHVPTDQVELDKHMRTAIDALKDRLQTR
```

```
                                     510       520       530       540       550       560       570       580       590       600
                                ....|....|....|....|....|....|....|....|....|....|....|....|....|....|....|....|....|....|....|....|
B._rhizoxinica_B1_(CBW77121)    VPGEVAAFFNRRPGLHETDKAKASHRLNKFVSEVATRDREHYGSHLIQRIFGTMKAP-----MVAATRIKSGAGKATE----------------------
Burkholderia_sp._B2_(MN840555)  VPGEVAAFFNRRPGLHETDKAKASHRLNKFVSEVATRDREHYGSHLIQRVFGTMKAP-----MVAATRIKSGAGKATE----------------------
Burkholderia_sp._B3_(MN840550)  VPAKVAALFNQRPGLHETDKAKASHRLNKFVSEVALRDREHCGSHLIQRIRGAMKAP-----MIAT-RIPSGASRAAE----------------------
Burkholderia_sp._B4_RBRH_00457  VPAKVAALFNQRPGLHETDKAKASHRLNKFVSEVALRDREHCGSHLIQRIRGAMKAP-----MIAT-RIPSGASRAAE----------------------
B._endofungorum_B5_(MN840553)   VPGKVAALFNQRPGLHETDKAKASHRLNKFVSEVAPRDREHCGSHLVQRIRGAMKAP-----MIAT-RIPSGASRAAE----------------------
Burkholderia_sp. B6_(MN840556)  VPGEVAAFFNRRPGLHETDKAKASHRLNKFVSEVATRDREHYGSHLIQRIFGTMKAP-----MVAATRIKSGAGKATE----------------------
Burkholderia_sp._B7_(MN840551)  VPAKVAALFNQRPGLHETDKAKASHRLNKFVSEVALRDREHCGSHLIQRIRGAMKAP-----MIAT-RIPSGASRAAE----------------------
Burkholderia_sp._B8_(MN840554)  VPGEVAALFNQRPDLRETDKAKASHRLNKFVSEVALRDRKHCGSHLIQRIWGAMKAP-----MIAT-RIPSGASRAAE----------------------
B._pseudomallei_(ZP04890375)    GAGAGAGAGAGAGAGVAKARSEVVSRVFEKLKIDGFSSKIEIKSDDLKKWVALAVAG------GRNDAVKRSLEFESIR----------------RDLKN
B._pseudomallei_(ZP02502372)    ----GAGAGAGAGAGVAKARSEVVSRVFEKLKIDGFSSKIEIKSDDLKKWVALAVAG------GRNDAVKRSLEFESIR----------------RDLKN
B._pseudomallei_(ZP02494128)    ----GAGAGAGAGAGVAKARSEVVSRVFEKLKIDGFSSKIEIKSDDLKKWVALAVAG------GRNDAVKRSLEFESIR----------------RDLKN
X._euvesicatoria_(WP011347306)  VARRQIQLAGGRMPSQDELAMQALLEYVQWLPHEQNATDLTFTAKVLGTIERRAQELQRSVAASVNETDDLQPAACHPAIAS-AWGALRAGRMRLPETLR
X._campestris_pv._musacearum_(  VARRHTQSACGQLPSQDELAMQALSEYVQWLPHEQNATDLIFTAKVLAEIEQRALELQDSVTASLSTTDDRQPAACHPAIAS-AWVALRTGRVRLPETLR
X._vasicola_pv._vasculorum_(AV  VARRHTQSACGQLPSQDELAMQALSEYVQWLPHEQNATDLIFTAKVLAEIEQRALELQDSVTASLSTTDDRQPAACHPAIAS-AWVALRTGRVRLPETLR
X._citri_(ARR12757)             VARRQIQRAGGRMPSQDELAMQALLEYAQWLPHEQNATDLTFTAKVLGTIERRAQELQRSVAASVNEADDLQPAACHPAIAS-AWAALRAGRMRLPETLR
X._campestris_(AKS22319)        VARRQIQLAGGRMPSQDELAMQALSEYAHWLPHEQNATDLTFTAKVLGTIERRAQELQRSVAASASESDDLQPAACHPAIAS-AWVALRAGRMRLPETLR
X._oryzae_(AKO19890)            VARRHIQMTCGQLPSQDELAMQALAECVQWLPHEQSATELIFTAKVLAEIEQRARELQDSVTASLSTTDARQPAACHPAIAS-AWVALRTSRMRLPETMR
X._oryzae_(ACD59124)            VARRHIQMTCGQLPSQDELAMQALAECVQWLPHEQSATELIFTAKVLAEIEQRARELQDSVTASLSTTDARQPAACHPAIAS-AWVALRTSRMRLPETLR
Burkholderia_sp._(WP013592484)  GRMLTHALNAERLQAWSAARPAVPSDAPLRELQKRRPESFEIGRRDARRMWDAARARVAALRGQGDERMERTVGRTLRMFDD----RERRRRFIDDMTEG
Burkholderia_sp._(ADN59567)     GRMLTHALNAQRLQAWTDARPDVPAHAKPRELQKRRPESFALGRREARRMWDAARARVAGLRVEGDQRTQRNVERALRMFDD----PQRRRRFVDDVTSS
R._solanacearum_(YP003748288)   SGKAQFSIGRHGEVPIVPLRAAILEHWSAASADQ-RPQGYTLDGNAVVDIAEGIRR------ATGKSVVGADGRLPAQ---------------LEQLIGT
R._solanacearum_(BAH04968)      EAAPLDPPGALPWRSPQEMWSSAVLKHWASALAQASPQDCVLTNDILAAIGRQLRDTVTRAFDALAAGIAQATEHDDGLREH---LRASLATLTRLSQGA
R._solanacearum_(BAD42384)      EAAPLDPPGALPWRSPQEMWSSAVLKHWASALAQASPQDCVLTNDILAAIGRQLRDTVTRAFDALAAGIAQATEHDDGLREH---LRASLATLTRLSQGA
R._solanacearum_(YP003747289)   RRLDGGHALPTERRGLLLLREAVLEHWRAQIGTTWRSSKLKLTDADKQGIARAVR------SAAHGTDIDAEAVLGYR--------EFRKLDKLDLKTLA
R._solanacearum_(WP013209172)   RQPGGGHSLSTEQRGLLLLREAVLKHWGAHIDPTWRASKLKLTDADKRDIAQAVR------SAAHGTGIDAKAVLSYR--------EFRKLDKLDLMTLA
R._solanacearum_(WP011002069)   KAASTSRALKTR----CAVRLAAIEQWERRMASKGLRSTFRFSSRDLETVAARARPLLSNRVALHADGSPADLALDPHRNRDAIRAEVKPLRGMTPAQLR
R._solanacearum_(WP003275417)   DAPARAVQR--GAIPLPVLRAAVLAHWADASETR-RPQGHVLDAAAVADIAGRLQLY---PAATATQSADVAKRLPTQ---------------LTSLIGT
R._solanacearum_(WP003274509)   RRLDGGHALSTERRGLLLLREAVLEHWRAQIGTTWRSSKLKLTDADKRDIARAVR------SAAHGTDVDAEAVLGYR--------EFRKLGKLDLKTLA
R._solanacearum_(EAP71656)      SGAAQFNIGRHGEVPTVTLRAAILEHWSAASADR-RPQGYTLDGNAVVDIAEGIRR------ATGKSVVGADGRLPAQ---------------LEQLIGT
R._solanacearum_(EAP71039)      DAPARAVQR--GAIPLPVLRAAVLAHWADASETR-RPQGHVLDAAAVADIAGRLQLY---PAATATQSADVAKRLPAQ---------------LTSLIGT
R._solanacearum_(CBJ39728)      RGLNDRHALSTEQRGLLLLREAVLQHWGASIGTTWRSSKLKLSDHDKRAIADRVR------KAAPGARVDAKDVLTYR--------EFRKLDKLDLKTLA
R._solanacearum_(CBJ35355)      SGEARFSVGRHGEVPTVILRTAILEHWSAASADK-RPQGYTLDGNAAVHIAECIHR------ATGQSVVRADGKLPKQ---------------LERLIGT
R._solanacearum_(CAD17997)      LALSP-AHADLPWKQAEQMWQGAILQHWASLPADAPLSAYVLTHDTLRTIGEQLRHTVLRVREELTISLAHSAEAGDTPDPG---MLDKLRTLVELSRQA
R._solanacearum_(BAH47286)      KAASTSRALKTR----CAVRLAAIEQWERRMASKGLRSTFRFSSRDLEAVAARARPLLSNRVALHADGSPADLALDPHRNRDAIRAEVKPLRGMTPAQLR
R._solanacearum_(BAH47283)      RALNGRHALSTEQRGLLVLREAVLQHWAQSIGTTWRSSKLKLSDHDKRAIADWVR------KAAPGARVDAEAVLAYR--------EFRKLDKLDLKTLV
R._solanacearum_(BAH04967)      LALSP-AHADLPWKQAEQMWQGAILQHWASLPADAPLSAYVLTHDTLRTIGEQLRHTVLRVREELTISLAHAAEAGDTPDPG---MLDKLKTLVELSRQA
R._solanacearum_(BAD42389)      SGEARFSLGRYGEVPTVALRGAILEHWSATSESK-RPQGYTLDGNAMVDIAERLHR------AIGESVVRADGRLPQQ---------------LERLIGT
```

```
                                         610       620       630       640       650       660       670       680       690       700
                                 ....|....|....|....|....|....|....|....|....|....|....|....|....|....|....|....|....|....|....|....|
B._rhizoxinica_B1_(CBW77121)     --ESSAADAQASAPD-------------------YASAASTAGAAEPTRAASTAIQSI-TTAVADASDRGWDALKRIVPSLRPGMAVEFSDGGALTVTTE
Burkholderia_sp._B2_(MN840555)   --ESSAADAQASAPD-------------------YASAASTAGAAEPTRAASTATQSI-TTAVADASDSGWDALKRIVPSLRPGMAVEFSDGGALTVTTE
Burkholderia_sp._B3_(MN840550)   --KSSAADMQASASD-------------------HASAASTAEASELTRAASTATQSV-TTAVADATDSGWDALKHIVPSLRPGMAVEFSDGGELAVTTE
Burkholderia_sp._B4_RBRH_00457   --KSSAADMQASAPD-------------------DASAASIAEASELTRAASTATQSV-TTAVADATDSGWDALKHIVPSLRPGMAVEFSDGGELAVTTE
B._endofungorum_B5_(MN840553)    --KSSAADLQASAPD-------------------HASAASTTEASELTRAASTATQSV-TTAVADATDSGWDALKHIVPSLRPGMAVEFSDGGELAVTTE
Burkholderia_sp. B6_(MN840556)   --ESSAADAQASAPD-------------------YASAASTAGAAEPTRAASTAIQSI-TTAVADASDRGWDALKRIVPSLRPGMAVEFSDGGALTVTTE
Burkholderia_sp._B7_(MN840551)   --KSSAADMQASASD-------------------HASAASTAEASELTRAASTATQSV-TTAVADATDSGWDALKHIVPSLRPGMAVEFSDGGELAVTTE
Burkholderia_sp._B8_(MN840554)   --KSSAADRQASAPD-------------------HASAASTAEASELTRAASTATQSV-TTAVTDATDSGWDALKRIVSSLRPGMAVEFSDAGKLTFTTE
B._pseudomallei_(ZP04890375)     GFSVDVLERWAAEAG----------------------LDDGNFRRQLNDLRSVSNNGYGEIPELGDRASVFSAFSEIVARYRAGKSLKFSSGGVGGLALG
B._pseudomallei_(ZP02502372)     GFSVDVLERWAAEAG----------------------LDDGNFRRQLNDLRSVSNNGYGVIPELGDRASVFSAFSEIVARYRAGKSLKFSSGGVGGLALG
B._pseudomallei_(ZP02494128)     GFSVDVLERWAAEAG----------------------LDDGNFRRQLNDLRSVSNNGYGEIPELGDRASVFSAFSEIVARYRAGKSLKFSSGGVGGLALG
X._euvesicatoria_(WP011347306)   LIQRGLLDHAGQPSDLPVG--------AATLDEELASDARTHFHQAIGHASRLLHEGD--TSRVRSPRALFECLRNLLERLEWRDKLRLTEQRVIGLNTT
X._campestris_pv._musacearum_(   LIQRGLLDHPVQPIDTPDG--------AAHLGKELASVARNNFHQAVGNASRFLHEGD--TSRVRSPRALFACLRNVMERLEWRDKLRITEQRVIGLNTT
X._vasicola_pv._vasculorum_(AV   LIQRGLLDHPVQPIDTPDD--------AAPLGKELASVARNNFHQAVGNASRFLHEGD--TSRVRSPRALFACLRNVMERLEWRDKLRITEQRVIGLNTT
X._citri_(ARR12757)              LIQRGLLDHAGQPSDLPVG--------AAPLDEELASVARDHFHRAVGHASRFLHEGD--TSRVKSPRALFECLRNLMERLEWRDKLRLTEQRVIGLNTT
X._campestris_(AKS22319)         LIQRGLLDHAGQPTDMPIG--------AAPLDEELASVARNHFHQAVGHASRFLHEGD--TSRVRSPRALFECLRNLMERLEWRDKLRITEQRVIGLNTT
X._oryzae_(AKO19890)             LIQRGLLDHAIQPNDTPAG--------AAPLDKELASLARNHFHQAVGNASRFLHEGD--TSRVRSPRALFACLRNMMERLEWRDKLRITEQRVIGLNTT
X._oryzae_(ACD59124)             LIQRGLLDHAIQPNDTPAG--------AAPLDKELASVARNHFHQAVGNASRFLHEGD--TSRVRSPRALFACLRNVMERLEWRDKLRITEQRVIGLNTT
Burkholderia_sp._(WP013592484)   GFTLAALESWFNASG---------------MAAADAARAEPTARHLQAAHDLLANPGI-APLDRITTRNLRKQHLALMAQNVPGTGFTSYDGNALGIDAS
Burkholderia_sp._(ADN59567)      GFTLAALERCFNASG---------------MAADHPVRAQPAQRNLQAAHDLLDNPGI-APLQRITTHNLREQHLALMAQNIPGTGFTSYDGNALGIDAS
R._solanacearum_(YP003748288)    RLNHATLKEWARDAA---------------MPQRTETGEETAFSSAMRRARNILRPDK-DKPADMSADSMRDYLKNFMADHNIGNSMTFADGGALGVNTS
R._solanacearum_(BAH04968)       GLPP-ARDSPIHPPPGLIGLAPSGALLRSWSP----AAPAGPLEKALKSAERIEQQDD-LQLREKTIEAARELIESLLTTIDAGGKLRLASGSALGVNTG
R._solanacearum_(BAD42384)       GLPP-ARDSPIHPPPGLIGLAPSGALLRSWSP----AAPAGPLEKALKSAERIEQQDD-LQLREKTIEAARELIESLLTTIDAGGKLRLASGSALGVNTG
R._solanacearum_(YP003747289)    KWIDEAASADSNPAA---------------SPMLRRAKAKIVEAGDIARGRPIKLKGT-TLADFRDAMTDAIGTMPLGNYVRYFDGGTYGLNANMTVNQH
R._solanacearum_(WP013209172)    RWLDEAASFDPDPSA---------------SPTLHLARTKIAEAGGIERGRPIKPQGA-TMADFRNAMTGAIDAMPLGNYVRYFDGGTYGASINMTVNQH
R._solanacearum_(WP011002069)    TWADEAWRLSGQAVP---------------EPIR--ANIDLVES---RLAGDIRPKPG-DTNAQLDAIGALVKQMPDIYDIRVSSGGTVGLGGVPSQSLA
R._solanacearum_(WP003275417)    RLTHATLEHWGHDAA---------------MPQRSVDGKETVFAGAMRRARNILDPGK-DVPADMTADSMRAFLKNFVIEHNGGNSMTFTDGGALGINTG
R._solanacearum_(WP003274509)    KWIDEAASADSNPSA---------------SPMLRRAKAKIVEAGDIARGRPIKLKGT-TLADFRDAMTDAIGTMPLGNYVRYFDGGTYGLNANMTVNQH
R._solanacearum_(EAP71656)       RLDHATLGTWAHDAA---------------MPQRTETGEETAFSGAMRRARNILRPDQ-DKPADMTADSMRDHLKNFMADHNIGNSMTFADGGALGVNTS
R._solanacearum_(EAP71039)       RLTHATLEHWGHDAA---------------MPQRSVDGKETVFAGAMRRARNILDPGK-DVPADMTADSMRAFLKNFVIEHNGGNSMTFTDGGALGINTG
R._solanacearum_(CBJ39728)       RWLDEARAFDSRASE---------------SPMLREAAEDLVKAGDIERGRPIKPQGT-TLADFREALTGAIDQMPLGNYVRYFDGATYGANTNMTVNQH
R._solanacearum_(CBJ35355)       QLSHATLKAWARDAA---------------MPQRTETGEETDFSSAMRRARNILKPGR-DKPLDMTAGNMREFLKNFMAEHNVGNTMTFTDGGALGVNTG
R._solanacearum_(CAD17997)       NLNLGASNLPVTPPPQFIDAYPSGAVLRKWSEQVDAGAASGPLRKALDTAERIEQQDD-LKLQEAAIDAARGVIESLITTIEGGGKLRLASGTSLGVNTG
R._solanacearum_(BAH47286)       TWADEAWRLSGQAVP---------------EPVR--ANIDLVES---RLAGDIRPKPG-DTNAQLDAIGALVKQMPDIYDIRVSSGGTVGLGGVPSQSLA
R._solanacearum_(BAH47283)       RWLDEARAFDPRASE---------------SPTLRAAADNIVKAGDIERGRPIKPRGT-TLADFREALTGAIGQMPLGNYVRYFDGATYGVNTNMTVNQH
R._solanacearum_(BAH04967)       NLNLGASNLHVAPPPQFIDAYPSGAVLRKWSEQVDAGAASGPLRKALDTAERIEQQDD-LKLQEAAIDAARGVIESLITTIEGGGKLRLASGTSLGVNTG
R._solanacearum_(BAD42389)       RLSHATLATWARDAA---------------MPQRTETGEETAFSSAMRRARNILEPGK-DKPVDMTADSMRAFLKNFTGEHNVGNAMTFTDGGALGVNTG
```

```
                                              710       720       730       740       750       760       770       780       790       800
                                     ....|....|....|....|....|....|....|....|....|....|....|....|....|....|....|....|....|....|....|....|
B._rhizoxinica_B1_(CBW77121)         RVSASVTAAVSQATQASPVPLPVVVPSINAKRSRKRKATVQMG-CTEQHAWLFIGTAHATGRGAGAGVLVGASFPGSLVGGTVGVQF--------YDSEH
Burkholderia_sp._B2_(MN840555)       RVSASVTAAVSQAAQASPVPLPVVVPSINAKRSRKRKATVQMG-CTEQHAWLFIGTAHATGRGAGAGVLVGASFPGSLVGGTVGVQF--------YDSEH
Burkholderia_sp._B3_(MN840550)       RVSASVTAAVSQAAQASPVPLPVVVPSINAKRSRKRKATVQMG-CTEQHAWLFIGTAHATGDGAGVGVLVGASLPGSLVGGTVGVQF--------YDSEH
Burkholderia_sp._B4_RBRH_00457       RVSASVTAAVSQAAQASPVPLPVVVPSINAKRSRKRKATVQMG-CTEQHAWLFIGTAHATGDGAGVGVLVGASLPGSLAGGTVGVQF--------YDSEH
B._endofungorum_B5_(MN840553)        RVSASVTAAVSEAAQASPVPLLVVVPSINAKRSRKRKATVQMG-CTEQHAWLFIGTAHATGDGAGVGVLVGASLPGSLVGGAVGVQF--------YDSEH
Burkholderia_sp. B6_(MN840556)       RVSASVTAAVSQATQASPVPLPVVVPSINAKRSRKRKATVQMG-CTEQHAWLFIGTAHATGRGAGAGVLVGASFPGSLVGGTVGVQF--------YDSEH
Burkholderia_sp._B7_(MN840551)       RVSASVTAAVSQAAQASPVPLPVVVPSINAKRSRKRKATVQMG-CTEQHAWLFIGTAHATGDGAGVGVLVGASLPGSLVGGTVGVQF--------YDSEH
Burkholderia_sp._B8_(MN840554)       RVSASITAAVSQAAQASPVPLPVVVPSINTKRSRERKATVQMG-CTEQHAWLFIGTAHATGDGAGVGVLAGASLPGSLVGGTTSVQF--------YDSEH
B._pseudomallei_(ZP04890375)         DSQSIGPLVSVSPT---------------LGAGGGRVATVEIGGASGRGGLIAVTRANSQKLKVGASVFAGPQFIHVVRAGGSANLELVS-------AEQ
B._pseudomallei_(ZP02502372)         DSQSIGPLVSVSPT---------------LGAGGGRVATVEIGGASGRGGLIAVTRANSQKLKVGASVFAGPQFIHVVRAGGSANLELVS-------AEQ
B._pseudomallei_(ZP02494128)         DSQSIGPLVSVSPT---------------LGAGGGRVATVEIGGASGRGGLIAVTRANSQKLKVGASVFAGPQFIHVVRAGGSANLELVS-------AEQ
X._euvesicatoria_(WP011347306)       PLSAALAIIPSGLG---------LKLSAGGQTSSDRSVEIYMG---RTGLSLQIGRQKARQFNASTGISAGVLLPGTEHAPVGITGAAEWR----MKKES
X._campestris_pv._musacearum_(       PLSAALAIIPSGLG---------LRLSAGGQISSDRSVEIYMG---RTGLSLQIGQQKARQFQASTGINAGVLLPGTERAPVGITGAADWR----IKKES
X._vasicola_pv._vasculorum_(AV       PLSAALAIIPSGLG---------LRLSAGGQISSDRSVEIYMG---RTGLSLQIGQQKARQFQASTGINAGVLLPGTERAPVGITGAADWR----IKKES
X._citri_(ARR12757)                  PLSAALAIIPSGLG---------LKLSAGGQTSSDRSVEIYMG---RTGLSLQIGRQKARQFNASTGISAGVLLPGTEHAPVGITGAAEWR----MKKES
X._campestris_(AKS22319)             PLSAALAIIPSGLG---------LKLSAGGQTSSDRSVEIYMG---RTGLSLQIGRQKARQFNASTGISAGVLLPGTERAPVGITGAAEWR----MKKES
X._oryzae_(AKO19890)                 PLSAALAIIPSGLG---------LKLSAGGQVSSDRSVEIYMG---RTGLSLQIGQQKARQFQASTGISAGVLLPGTEHAPVGITGAAEWR----IKKES
X._oryzae_(ACD59124)                 PLSAALAIIPSGLG---------LKLSAGGQVSSDRSVEIYMG---RTGLSLQIGQQKARQFQASTGISAGVLLPGTEHAPVGITGAAEWR----IKKES
Burkholderia_sp._(WP013592484)       AWANLPAKLLVSAG-------------PIVTVLGGRDSLYSIG-TAYEGGQMIFGTRSRVNGALGAQGFVGAALDVGGVAGMAGGWASSSV-----GGYL
Burkholderia_sp._(ADN59567)          AWGNLPARLLVSAG-------------PIVTVLGGRDSMYSIG-TAYEGGQMMFGTRSRVNGTLGAQGFVGATLDVAGVAGMAGAWAASSV-----GGYL
R._solanacearum_(YP003748288)        ALTLNLARIVKRFKGGF-----ALTPVFDAQASVTRSAAFSIG-TTAHGGDIFIGRQRQIGGQLGGGLTAGYTTPTVANEDNFSASVGVSANVTLFGLEH
R._solanacearum_(BAH04968)           TLSTGLQNAGHALAIP-------LSVQADFHASRKRQAVVEFG-RGAHGRDLFIGTDKTVHAAAGCGGAIGYDIQVLANRIRASLGMSVVPIDVERGKRV
R._solanacearum_(BAD42384)           TLSTGLQNAGHALAIP-------LSVQADFHASRKRQAVVEFG-RGAHGRDLFIGTDKTVHAAAGCGGAIGYDIQVLANRIRASLGMSVVPIDVERGKRV
R._solanacearum_(YP003747289)        EFTHNSPVPAMGLG-------------PGAKALHGRHAFVEIG-SSSYGGEVFIGTDKRSSTGAGLGAYAGLKFGTEDWHLSAGFSAGVAH-----SQDR
R._solanacearum_(WP013209172)        AFSHNSLVPALGFG-------------PGAKTLRGRHAFVEIG-SSSYGGEVFIGTDKRSSDGTGMGAYAGLKFGTEDWHLSAGFSAGVTH-----SHDR
R._solanacearum_(WP011002069)        ALSSHLGVPHVSVL-------------PDGGYLQGRHAVIDIG-SNQHFGHLFIGTESRKSLYGGLGGYAGWSFG-QDGMANVGVSGGLRR-----SRDW
R._solanacearum_(WP003275417)        ALSVNVAMAAAAAT-QV-----LVQPIVDVQASVARSAVFTIG-TSSHGGEIFIGRQRQLAGQVGGGVMIGYVPSTEPGEAAFAARLGVAVNTTLFGLEH
R._solanacearum_(WP003274509)        EFTHNSPVPAMGLG-------------PGAKALHGRHAFVEIG-SSSYGGEVFIGTDKRSSTGAGLGAYAGLKFGTEDWHLSAGFSAGVAH-----SQDR
R._solanacearum_(EAP71656)           ALTLNLARIVKRLKGGF-----ALTPVFDAQASVARSAAFSIG-TTAHGGDIFIGRQRQIGGQLGGGLTAGYTTPTVADENSVSASVGVSANVTLFGLEH
R._solanacearum_(EAP71039)           ALSVNVAMAAAAAT-QV-----LVQPIVDVQASVARSAVFTIG-TSSHGGEIFIGRQRQLAGQVGGGVMIGYVPSTEPGETAFAARLGVAVNTTLFGLEH
R._solanacearum_(CBJ39728)           AFSHNSLVPSMGLG-------------PGARELTGRHAFVEIG-SSSYGGEVFIGTDKRSSTGAGVGLYAGFKIGIKNWHLSAGFSAGVAH-----AHDR
R._solanacearum_(CBJ35355)           ALTLNLARIVKRLKGGV-----TLTPVLDVQGSVARSAAFNIG-TTAHGGEIFIGRQRQIGGQIGGGITAGYSPPTDPGEDSFSASFGVSANVTLFGLEH
R._solanacearum_(CAD17997)           TISTGLQNARQALAVP-------VAIQADFHASHKRQAVVEIA-RGSHGRDLFVGTDKTVLAAAGFGASVGYDFKVLQNRIRAMVGASVVPIDVEKGERV
R._solanacearum_(BAH47286)           ALSSHLGVPHVSVL-------------PDGGYLQGRHAVIDIG-SNQHFGHLFIGTESRKSLYGGLGGYAGWSFG-QDGMANVGVSDGLRR-----SRDW
R._solanacearum_(BAH47283)           EFSHNSLVPSMGLG-------------PGARELTGRHAFVEIG-SSSYGGEVFIGTDKRSSTGVGAGLYAGFKVGIKNWHLSAGFSAGVAH-----AHDR
R._solanacearum_(BAH04967)           TISTGLQNARQALAVP-------VAIQADFHASHKRQAVVEIA-RGSHGRDLFVGTDKTVLAAAGFGASVGYDFKVLQNRIRAMVGASVVPIDVEKGERV
R._solanacearum_(BAD42389)           ALTLNLASLVKRFKGGF-----ALTPVFDAQGSVARSAAFNIG-TTAHGGEIFIGRQRQVVGQLGGGLTAGYTTPTDADENSFSAGVGISGNVTLFGLEH
```

```
                                              810       820       830       840       850       860       870       880       890       900
                                         ....|....|....|....|....|....|....|....|....|....|....|....|....|....|....|....|....|....|....|....|
B._rhizoxinica_B1_(CBW77121)             ARPRGILVQVARQPKADGSGY-------DDEAMEARMGHIIDTMMSLSPAGATAESRGTREQCWNALAQSLVGAR--DVSVGWIDGTVDEKRHGSSTGPV
Burkholderia_sp._B2_(MN840555)           ARPRGILVQVARQPKADGSGY-------DDEAMEARMGHIIDTMMSLSPAGATAESRGTREQCWNALAKSLVGAR--DVSVGWIDGTVDEKRHGSSTGPV
Burkholderia_sp._B3_(MN840550)           ARPRGILVQVARQLKEDGSGY-------DDEAMQASVNHILDTMKSLSPAGAAAESRGTSEQCWNTLAQSLVGAR--DVSVGWIEGTVDEKRHGASTGPL
Burkholderia_sp._B4_RBRH_00457           ARPRGILVQVARQLKEDGSGY-------DDEAMQASMNHILDTMKSLSPAGAAAESRGTSEQCWNTLAQSLVGAR--DVSVGWIEGTVDEKRHGASMGPL
B._endofungorum_B5_(MN840553)            ARPRGILVQVARQLKEDGSGY-------DDEVMQARVNHILDTMKSLLPAGAAAKSRGTSEQCWNTLAQSLVGAR--DVSVGWIEGTVDEKRHGASMGPL
Burkholderia_sp. B6_(MN840556)           ARPRGILVQVARQPKADGSGY-------DDEAMEARMGHIIDTMMSLSPAGATAESRGTREQCWNALAQSLVGAR--DVSVGWIDGTVDEKRHGSSTGPV
Burkholderia_sp._B7_(MN840551)           ARPRGILVQVARQLKEDGSGY-------DDEAMQASVNHILDTMKSLSPAGAAAESRGTSEQCWNTLAQSLVGAR--DVSVGWIEGTVDEKRHGASTGPL
Burkholderia_sp._B8_(MN840554)           ARPRGILVQVARQLKDDGSGY-------DDEAMQASVNHILDTMKSLSSAGATAESRGTSEQCWNTLAQSLVGAR--DVSVGWIEGTVDEKRHAASTGPL
B._pseudomallei_(ZP04890375)             TREKGAIVRFVPRAGSLGGGE----------AWRGWAQDCLSIIGNSADS----------ADLLERLAEEYVNAG--EIGIGMTEKEENVISSSAAVGAG
B._pseudomallei_(ZP02502372)             SREKGAIVRFVPRAGSLGGGE----------AWRGWAQDCLSIIGNSADS----------ADLLERLAEEYVNAG--EIGIGMTEKEENVISSSAAVGAG
B._pseudomallei_(ZP02494128)             TREKGAIVRFVPRAGSLGGGE----------AWRGWAQDCLSIIGNSADS----------ADLLERLAEEYVNAG--EIGIGMTEKEENVISSSAAVGAG
X._euvesicatoria_(WP011347306)           GIEHGVQIRVPRRGKGQELEQ------------RAQFLAMFEHLLQLAEQSGDDAGQLSERDFLGELLAHHPSIT--VGLIGQAERNTSATESSLSVAAG
X._campestris_pv._musacearum_(           GVEHGVQIRVPRRGKGQELEQ------------RAQFLAMFEHLLQLAEQSGGDTAQLTERDFLAELLAHHPSIT--VGLIGHAERNSSTTESSVSVAAG
X._vasicola_pv._vasculorum_(AV           GVEHGVQIRVPRRGKGQELEQ------------RAQFLAMFEHLLQLAEQSGGDTAQLTERDFLAELLAHHPSIT--VGLIGHAERNSSTTESSVSVAAG
X._citri_(ARR12757)                      GIEHGVQIRVPRRGKGQELEQ------------RAQFLAMFEHLLQLAEQSGGDAGQLSERDFLGELLAHHPSIT--VGLIGHAERNATATESSLSVAAG
X._campestris_(AKS22319)                 SIEHGVQIRVPRRGKGQELEQ------------RAQFLAMFEHLLQLAEQSGGDAGQLTERDFLGELLAHHPSIT--VGLIGHAERNASATESSLSVAAG
X._oryzae_(AKO19890)                     GIEHGVQIRVPRRGKGQELEQ------------RAQFLAMFEHLLQLAEQSGGDTSQLTERDFLAELLAHHPSIT--VGLIGHAERNSSTTESSVSVAAG
X._oryzae_(ACD59124)                     GIEHGVQIRVPRRGKGQELEQ------------RAQFLAMFEHLLQLAEQSGGDTAQLTERDFLAELLAHHPSIT--VGLIGHAERNSSTTESSVSVAAG
Burkholderia_sp._(WP013592484)           AREQSVVLRARRLAPTPATAA------LEAELWHKDSRQMVNAYWDAAEQ-AQTPEEFMQILAAGIADNERISL--SSQNNNATSITATPLTVGGVARAN
Burkholderia_sp._(ADN59567)              AREESVVLRARRLAPTPATAE------LEAELWHKDSQQMVNAYWDAAEQ-AQTPDEFMQILAAGIADNERISL--SSQNNNATTVTATPLTVGGVARAN
R._solanacearum_(YP003748288)            TNPVGVRLRFSTQRKEDGSSM------NSD-AMRKQMSDMIDYMFDACG---PAKRHLSPAQLWEDFAVHHFDQ--KNLSVGWADYSSLNSKMGGSVTLT
R._solanacearum_(BAH04968)               GVMFRAVRQLDRAQQTVGGYHDWEAVKYDDTATRQALISLSNWLFEQAT--EHRQRPLEREAIWNALALHCGTG--NTISVGWVDQRKRRLRHKLRLSGG
R._solanacearum_(BAD42384)               GVMFRAVRQLDRAQQTVGGYHDWEAVKYDDTATRQALISLSNWLFEQAT--EHRQRPLEREAIWNALALHCGTG--NTISVGWVDQRKRRLRHKLRLSGG
R._solanacearum_(YP003747289)            SAPAGVIIRTG---LSYGADG------KATSAWRDNVAEVTRFLFQTAAL-GQSTRPVPPERMWEQFAERFFRT--PDISVNWRDQRRSSETVTKHGSAA
R._solanacearum_(WP013209172)            SAPAGVIIRTG---LSYDAEG------KPTSAWRDNVADVTRFLFQTAAV-GQSARPVPPDRMWEQFADRFFRT--PDISVNWRDQQRSNHTVTKHGSAA
R._solanacearum_(WP011002069)            GGPRGVTIRTR---RSDDEQP------GRPDAWRTTMLEVLHATRSAGPN-SDAPR--NAREMWGSLARRFWND--PACSINWTDSRSTTSATSGSASAT
R._solanacearum_(WP003275417)            NHPVGVRLRFARQRNASGNDM------VDKGVMRRQMETMVDYLFDACG---PAKRHLSPAALWEDFANHHFDQ--ENLSVAWQDYSALSSRMGVSVTGS
R._solanacearum_(WP003274509)            SAPAGVIIRTG---LSYGADG------KATSAWRDNVAEVTRFLFQTAAL-GQATRPVPPERMWEQFAERFFRT--PDISVNWRDQRRSSETVTKHGSGA
R._solanacearum_(EAP71656)               TNPVGVRLRFSTQRKEDGSGM------NSD-AMRKQMSDMIDYMFDACG---PAKRHLSPAQLWEDFAVHHFDQ--KNLSVGWADYSTLNSKMGGSVTLT
R._solanacearum_(EAP71039)               NHPVGVRLRFARQRNASGNDM------VDKGVMRRQMETMVDYLFDACG---PAKRHLSPAALWEDFANHHFDQ--ENLSVAWQDYSALSSRMGVSVTGS
R._solanacearum_(CBJ39728)               SAPVGVIVRTG---LTYGTDG------KATNAWRDNVADVTRFLFQSAAE-GQAARPVPPDRMWEQFSARFFRT--PDISVNWRDQRRSSHTVTKHGSGA
R._solanacearum_(CBJ35355)               TNPTGVRLRFTTQRKEDGTAM------NSD-AMRKQMSDMVDYMFDACG---PAKRHLSPSELWEDFAVHHFDQ--KHLSVGWEDYSSLNSKMGGAITAT
R._solanacearum_(CAD17997)               GVMYRAVRQVT---QAAGPHADWDAVKYDDAATREAMIALSNWLFQQAT--EHRERPPGADAVWNALAQHCSGAGADAISVGWVDQRKHRLRHKLRVGGG
R._solanacearum_(BAH47286)               GGPRGVTIRTR---RSDDEQP------GRPDAWRTTMLEVLHAARSAGPN-SDAPR--NAREMWGSLARRFWND--PACSINWTDSRSTTSATSGSASAT
R._solanacearum_(BAH47283)               SAPVGVIVRTG---LTYGADG------KATSAWRDNVADVTRFLFQTAAE-GQAARPVPPDRMWEQFSARFFRT--PDISVNWRDQRRSSHTVTKHGSGA
R._solanacearum_(BAH04967)               GVMYRAVRQVT---QAAGPHADWDAVKYDDAATREAMIALSNWLFQQAT--EHRERPPGADAVWNALAQHCSGAGADAISVGWVDQRKHRLRHKLRVGGG
R._solanacearum_(BAD42389)               TNPTGVRLRFTTQRKDDGSAM------NSD-AMRKQMSDMVDYMFEACG---PDKRHLSPSALWEDFAVHHFDQ—KNLSVAWEDYASVNSKMGASVTLT
```

```
                                          910       920       930       940       950       960       970       980       990      1000
                                 ....|....|....|....|....|....|....|....|....|....|....|....|....|....|....|....|....|....|....|....|
B._rhizoxinica_B1_(CBW77121)     ASAGVPIGLPHPFGMSTSVSVPVSGKHVHN-ISTSRSDSGRMSTEKRHAERSIEVKVEVRVGASVSATLSSNPKSGISTG-------------------P
Burkholderia_sp._B2_(MN840555)   VSAGVPIGLPHPFGMSTSVSVPVSGKHVNN-ISTSRSDSGRMPTEKRHAERSIEVKVEVKVGASVSATLSSNPKSGVNTG-------------------P
Burkholderia_sp._B3_(MN840550)   VSTGVPIGLPAPLSTSVSASVPVSGKHVNS-ISTSRNDSGRMSTEKRHTERSIEVKIEAKVGASISATTSSSPKSGVSTG-------------------P
Burkholderia_sp._B4_RBRH_00457   VSTGVPIGLPAPLSTSVSASVPVSGKHVNS-ISTSRSDSGRMSTEKRHTERSIEVKIEAKVGASISATTSSSPKSGVSTG-------------------P
B._endofungorum_B5_(MN840553)    VSAGVPLGLPAPFSTSVSASVPVSGKHVNS-ISTSRSDSGRMSTEKRHTERSIEVKIETKVGASISATTSNSPKSGVSTG-------------------P
Burkholderia_sp. B6_(MN840556)   ASAGVPIGLPHPFGMSTSVSVPVSGKHVHN-ISTSRSDSGRMSTEKRHAERSIEVKVEVRVGASVSATLSSNPKSGISTG-------------------P
Burkholderia_sp._B7_(MN840551)   VSTGVPIGLPAPLSTSVSASVPVSGKHVNS-ISTSRNDSGRMSTEKRHTERSIEVKIEAKVGASISATTSSSPKSGVSTG-------------------P
Burkholderia_sp._B8_(MN840554)   VSVGIPNVLPAPLGASVSASVPVSGKHVNS-ISTSRSDSGHMSTEKRHAERSIEVKVEAKVGASVSAATSSSPKSSVSTG-------------------P
B._pseudomallei_(ZP04890375)     VKASSSGG---DFRGGAGVSASIGVKREWRRHKAVRELDNKMGNHKFDQSSSTSVTWNAGLNASESYQLNPTPDFSTSAS------------------AS
B._pseudomallei_(ZP02502372)     VKASSSGG---DFRGGAGVSASIGVKREWRRHKAVRELDNKMGNHKFDQSSSTSVTWNAGLNASESYQLNPTPDFGTSAS------------------AS
B._pseudomallei_(ZP02494128)     VKASSSGG---DFRGGAGVSASIGVKREWRRHKAVRELDNKMGNHKFDQSSSTSVTWNAGLNASESYQLNPTPDFSTSAS------------------AS
X._euvesicatoria_(WP011347306)   VRVGNMDGQPRRATLGASLGVKARRETSRS----QTPIEGYMTMVLKDSSAQSRVEIAGRASASVVARQWTDPGTGNAPP-----------TPVARLSMA
X._campestris_pv._musacearum_(   VRVGNMDGRPRRATLGASLGIKARRETSRT----QTPIEGYMTMVLKDSSAQSRVEIAGRASASLVAQQWTQPSTGNAPP-----------NPIARLSMA
X._vasicola_pv._vasculorum_(AV   VRVGNMDGRPRRATLGASLGLKARRETSRT----QTPIEGYMTMVLKDSSAQSRVEIAGRASASLVAQQWTQPSTGNAPP-----------NPIARLSMA
X._citri_(ARR12757)              VRVGNMDGRPRRATLGASLGLKARRETSRS----QTPIEGYMTMVLKDSSAQSRVEIAGRASASLVAQQWTHPSTGNAPP-----------TPVARLSMA
X._campestris_(AKS22319)         VRVGNMDGKPRRATLGASLGLKARRETSRS----QTPIEGYMTMVLKDSSAQSRVELAGRASASVVAQQWTHPSIGNAPP-----------TPVARLSMA
X._oryzae_(AKO19890)             VRVGNMDGRPRRATLGASLGLKARRETSRS----QTPIEGYMTMVLKDSTAQSRVDIAGRASASLVAQQWTQPSAGNAPP-----------SPVARLSMA
X._oryzae_(ACD59124)             VRVGNMDGRPRRATLGASLGLKARRETSRS----QTPIEGYMTMVLKDSTAQSRVDIAGRASASLVAQQWTQPSAGNAPP-----------SPVARLSMA
Burkholderia_sp._(WP013592484)   VPGASSS-----ERVGGYAGVSSG-VTLYS-AADLKESGALASTIGATAVQGSLTASVG--AQFAPRAFALGDGTGEIGS---------------VSPGA
Burkholderia_sp._(ADN59567)      VPGANSP-----ERVGAYAGVSSG-VTLYS-AADLKESGALASTIGATAVQGTLSASAG--TQFAPRALMLGDGAGEVGS---------------VSPGA
R._solanacearum_(YP003748288)    ARAGVTTEDGQAIRAGGSIGYGVT-WNPFT-MGSRKEKSGTAPILREDRGSAHIQTLTATASGQLPAVPLPVEPDGAANG---------------LGVPG
R._solanacearum_(BAH04968)       VQARISSK-GAPIRVGPSARLTAE-ATSRD-ATATRETAGIMRVEQSVQGHGGHLTLRAGLTYSAGKMFYTSAAPDAATG----SDATGGRHRITQGFNG
R._solanacearum_(BAD42384)       VQARISSK-GAPICVGPSARLTAE-ATSRD-ATATRETAGIMRVEQSVQGHGGHLTLRAGLTYSAGKMFYTSAAPDAATG----SDATGGRHRITQGFNG
R._solanacearum_(YP003747289)    MRVAAGP-----VRLGPAFSVGHD-HVLAS-KNDRVDTNGWLRGVERARARASNVHVGASLAALAPAVGHFSNRSGFPDT---------------ITLPS
R._solanacearum_(WP013209172)    VRVAAGP-----VRVGPAFAVGHE-RLLAS-KNDRIDDNGWLRGVERARAQSSSVHVSASLAAVAPAIGHFSNHSGFPDS---------------ITLPS
R._solanacearum_(WP011002069)    ARVGTAD-----TKWGPALGATLR-HVARA-AHRQWDKTGNHAIDVSTHNSGRATAVAATLVEALPGIP-VPNGSGHLAA---------------LSFPT
R._solanacearum_(WP003275417)    ARLGVKTDGGETVRTGGLFGYAFN-WSPFT-RGKRQEASGRFPVMRNERGVMYSHTVTVATGTQLPSVPLPDSHEGVSDS---------------LGLPN
R._solanacearum_(WP003274509)    VRVAAGP-----VRLGPAFSVGHD-QVLAS-KNDRVDTNGWLRGVERARARASNVHVGASLAALAPAVGHFSNRSGFPDS---------------ITLPS
R._solanacearum_(EAP71656)       ARAGVTTEDGQAIRAGGSIGYGVT-WNPFT-MGSRKEKSGTAPILREDRGSAHIHTLTATASGQLPAVPLPAQPDGATNS---------------LGVPA
R._solanacearum_(EAP71039)       ARLGVKTDGGETVRTGGLFGYAFN-WSPFT-RGKRQEASGRFPVMRNERGVMYSHTVTVATGTQLPSVPLPDSHEGVSDS---------------LGLPN
R._solanacearum_(CBJ39728)       VRVAAGQ-----VRVGPAFSVGHD-RVLAS-KNDRVDTNGWLRGVERARARASNVHVSGSLAAVAQGVGHFSNRSGFPES---------------VTLPS
R._solanacearum_(CBJ35355)       ARAGVKTADGQTVRGGGSIGYGFT-WNPFT-MGERKEKSGTAPILREDRGSAHIHTVTVAASGQLPGVPLPDSLDGAANS---------------LGVPS
R._solanacearum_(CAD17997)       VNVRVTAP-AAPVRVGVGVRLTGE-ITSRD-TTAMRETTGIMRAEQYSQGHGGHMTVRAGVNYAVGKRFYTSPAQDAAAAGPTASVPSGGRRSITQGFNG
R._solanacearum_(BAH47286)       ARVGTAD-----TKWGPALGATLR-HVARA-AHRQWDKTGNHAIDVSTHNSGRATAVAATLVEALPGIP-VPNGSGHLAA---------------LSFPT
R._solanacearum_(BAH47283)       VRVAAGA-----VRVGPAFSVGHD-QVLAS-KNDRVDTNGWLRGVERARARASNVHVSGSLAAVAQGVGHFSNRSGFPES---------------VTLPS
R._solanacearum_(BAH04967)       VNVRVTAP-AAPVRVGVGVRLTGE-ITSRD-TTAMRETTGIMRAEQYSQGHGGHMTVRAGVNYAVGKRFYTSPAQDAAAAGPTASVPSGERRSITQGFNG
R._solanacearum_(BAD42389)       ARAGVTTQDGQTVRAGGSVGYGFT-WNPFT-IGQRQEKSGIAPILREDRGSAHLHAVTVAASVQLPAGPLPDTPDGAANS---------------LGVPS
```

```
                                           1010      1020      1030      1040      1050      1060      1070      1080      1090      1100
                                   ....|....|....|....|....|....|....|....|....|....|....|....|....|....|....|....|....|....|....|....|
B._rhizoxinica_B1_(CBW77121)       SSKRLWDKQLYQAKVSLSRTLQSYQGKIEPRACFLDVDFLDATAYEYATRQLLDHNGDRDDAGIKAVRTYLEQKRAQSAKPLADQAASAVKNVLELGNGT
Burkholderia_sp._B2_(MN840555)     SSKRLWDKQLYQAKVSLSRTLQSYQGKIEPRACFLDVDFPDATAYEYATRQLLDHNGDRDDAGIKAVRTYLEQKRAQSAKPLADQAASAVKNVLELGNGT
Burkholderia_sp._B3_(MN840550)     TSKRLWDKQIYQVKVSLSRTLQTHHGKIDPRACFLDVDFPDATAYEYATRQLLNHDGDHDDTGIKAVGTYLEQKRAQSTEPLANQVASAVKNVLELGNGT
Burkholderia_sp._B4_RBRH_00457     TSKRLWDKQIYQVKVSLSRKLQTHHGKIDPRACFLDVDFPDATAYEYATRQLLDHDGDHDDTGIKAVGTYLEQKRAQSTEPLANQVASAVKNVLELGNGT
B._endofungorum_B5_(MN840553)      TSKRLWDKQIYQVKVSLSRTLQTHHGKIDPRACFLDVDFPDATAYEYATRQLLDHDGAHDDAGIKAVGTYLEQKRAQSAEPLANQVASAVKNVLELGNGT
Burkholderia_sp. B6_(MN840556)     SSKRLWDKQLYQAKVSLSRTLQSYQGKIEPRACFLDVDFLDATAYEYATRQLLDHNGDRDDAGIKAVRTYLEQKRAQSAKPLADQAASAVKNVLELGNGT
Burkholderia_sp._B7_(MN840551)     TSKRLWDKQIYQVKVSLSRTLQTHHGKIDPRACFLDVDFPDATAYEYATRQLLNHDGDHDDTGIKAVGTYLEQKRAQSTEPLANQVASAVKNVLELGNGT
Burkholderia_sp._B8_(MN840554)     TSKRLWDKQIYQVKVSLSRTVQSYQGKIDPRACFLDVNFPDATTYEYATRQLLDHDGDHDDTGIKAVGTYLKQKRTQSAKLLTDQVASAVKNMLELGNGT
B._pseudomallei_(ZP04890375)       AQAFSVGGTLFSAKEGASVAIRLRDGMVDSELAILERTRNRGRRGVELIARDYPLWVKAMGKTDAEGEAALNR-----------FFASELGRGVNRPLRP
B._pseudomallei_(ZP02502372)       AQAFSIGGTLFSVKEGTSVAIRLRDGMVDAELAILDRTRNGGRRAVDLIARDYPIWVKAMGKTDAEGEAALNR-----------FFASELGRGVNRPLRP
B._pseudomallei_(ZP02494128)       AQAFSIGGTLFSVKEGTSVAIRLRDGMVDAELAILDRTRNGGRRAVDLIARDYPIWVKAMGKTDAEGEAALNR-----------FFASELGRGVNRPLRP
X._euvesicatoria_(WP011347306)     TLDLGYSRELRSAGSTTFSTLWMFNNEIDPVRTDRGFEFFNFASFEREVKRNWQLWTHYGISKLQGKVDDQR------LYMVAERQLQDFIDRVRLHMQD
X._campestris_pv._musacearum_(     ALDLGYSREVHSAGSNTFSTLWMFNNEIDPVRTDRGFEFFDVASFEREVKRNWQLWTHYGISKLQGKVDDQR------LYMVAERQLQDFVDRVRIHMQD
X._vasicola_pv._vasculorum_(AV     ALDLGYSREVHSAGSNTFSTLWMFNNEIDPVRTDRGFEFFDVASFEREVKRNWQLWTHYGISKLQGKVDDQR------LYMVAERQLQDFVDRVRIHMQD
X._citri_(ARR12757)                TLDLGYSRELRSAGSNTFSTLWMFNNEIDPVRTDRGFEFFNFASFEREVKRNWQLWTHYGISKLQGKVDEQR------LYMVAERQLQDFIDRVRLHMQD
X._campestris_(AKS22319)           TLDLGYSRELHSAGSNTFSTLWMFNNEIDPVRTDRGFEFFDFASFEREVKRNWQLWTHYGISKLQGKVDDQR------VYMVAERQLQDFVDRVRLHMQD
X._oryzae_(AKO19890)               ALDLGYSRELHLAGSNTFSTLWMFNNEIDPVRTDRGFEFFNFASFEREVKRNWQLWTNYGISKLQGKVDDHR------LYMVAERQLQDFIDRVRIHMQD
X._oryzae_(ACD59124)               SLDLGYSRELHLAGSNTFSTLWMFNNEIDPVRTDRGFEFFNFASFEREVKRNWQLWTHYGISKLQGKVDDQR------LYMVAERQLQDFIDRVRIHMQD
Burkholderia_sp._(WP013592484)     KTLGGAAVELTPFGRSSFVFAMRDNRGFVAEYTMQDRMFRNADDWARSVRAN-------PAWAHGIGA----------------ERLQQVIEQTQQ--ES
Burkholderia_sp._(ADN59567)        KTLASAAVEATPFGRSSFVFAMRDNRGFVAEYTMQDRMFRNADDWARSVRAN-------PAWLQGIGA----------------QRLQQVIEQTQL--ES
R._solanacearum_(YP003748288)      VPVASATYMLGNGGFNANIRTLVERGRLSEAFTYRDLSERNIDDFLKFANDPARRKEWEAMCGAEQGAYAEHG--AGGPDLGKKRLDDFLGKIQEM--AR
R._solanacearum_(BAH04968)         GTPLSWSKQFAELGHTVEVRTIHLNGRLADRVSYSDKRYKSGTEFLALIRTRQRVWIDMLSRKPGMTREAAE------------QEFKVFCDMVER--HS
R._solanacearum_(BAD42384)         GTPLSWSKQFAELGHTVEVRTIHLNGRLADRVSYSDKRYKSGTEFLALIRTRQRVWIDMLSRKPGMTREAAE------------QEFKVFCDMVER--HS
R._solanacearum_(YP003747289)      TPIVGLSANILPSGTSVTLRRVDEHGRLNPRFIRRLVEFIDPSAFLSHMQAR----MPRIARTDASRE----------------KVGAFMSAMKDM--GT
R._solanacearum_(WP013209172)      APIVGVAANIMPSGTSVTLRRVDEHGRLNPRFIRRLVEFIDPSAFLSHMQAR----LPRIAHTDASRA----------------KVGAFMSAMDDM--GT
R._solanacearum_(WP011002069)      QPYVGIGTTLFTTQTNAALRIGRDSGRIIPKHTFRDTEFGTFKAFKQFVDTHRSEWLTALGGTDDARR----------------RLDEMVATVKAR--AT
R._solanacearum_(WP003275417)      VPIHSATYMLGEGGFNAIVRTLMEHGRHSETFTYRDLSERSIHDFLKIANEPGRRKAWEALCAAEQGGDAARG---------KERLDDFLDKVREM--AR
R._solanacearum_(WP003274509)      TPIVGLSANILPSGTSVTLRRVDEHGRLNPRFIRRLVEFIDPSAFLSHMQAR----MPRIARTDASRE----------------KVGAFMSAMKDM--GT
R._solanacearum_(EAP71656)         VPVASATYMVGNGGFNANIRTLVERGRLSEAFTYRDLSERSIDDFLKFANDPARRKEWEAVCGAEQGAYAEHG--EGGPDLGKKRLDDFLDKIREM--AR
R._solanacearum_(EAP71039)         VPIHSATYMLGEGGFNAIVRTLMEHGRHSETFTYRDLSERSIHDFLKIANEPGRRKAWEALCAAEQGGDAARG---------KERLDDFLDKVREM--AR
R._solanacearum_(CBJ39728)         TPIVGLSANILPSGTSVTLRRVDEHGRLNPRFIRRLVEFINPSAFLSHMQAR----LPQIAHSDASRA----------------KVGAFMSAMKDM--GT
R._solanacearum_(CBJ35355)         VPVASATYMLGNGGFNATVRTLLERGRLSEAFTYRDLSERNIGDFLKFANDPARRKEWEALCSAEQGAYADTGHGEAGPDLGKKRLDDFLSKIQEM--AR
R._solanacearum_(CAD17997)         GTPLTYTKQFAEHSHTAKLKTVHLDGRLADRVCYFDKEYKSAKEYLALIRSAQDTWIDMLSRKPGGTVEKAR------------KEFEDYCATVEQ--HT
R._solanacearum_(BAH47286)         QPYVGIGTTLFATQTNAALRIGRDSGRIIPKHTFRDTEFGTFKAFKQFVDTHRSEWLTALGGTDDARR----------------RLDEMVATVKAR--AT
R._solanacearum_(BAH47283)         VPIVGASANILPSGTSVTLRRVDEHGRLNPRFIRRLVEFIDPSAFLSHMQAR----LPQIAHSDASRA----------------KVGAFMSAMRDM--GT
R._solanacearum_(BAH04967)         GTPLTYTKQFAEHSHTAKLRTVHLDGRLADRVCYFDKEYKSAKEYLALIRSAQDTWIDMLSRKPGCTVEKAR------------KEFEDYCATIEQ--HT
R._solanacearum_(BAD42389)         VPIASATYMLGNGGFNATIRTLMERGRLSEAFTYRDLSERNINDFIKFANDPARRKEWEALCSAEQGAYAAHGKADAGP--GKKRLDDFLDKIQEM-AR
```

```
                                         1110      1120      1130
                                ....|....|....|....|....|....|...
B._rhizoxinica_B1_(CBW77121)    RQTDSHIHRYALTDPAAEQINLLQAHFEQHYLR
Burkholderia_sp._B2_(MN840555)  RQTDSHIHRYALTDPAAEQINLLQAHFEQHYLR
Burkholderia_sp._B3_(MN840550)  RQMHSHIHRYALTDPAAEQINLLQAHFEQHYLR
Burkholderia_sp._B4_RBRH_00457  RQMHSHIHRYALTDPAAEQINLLQAHFEQHYLR
B._endofungorum_B5_(MN840553)   RQTDSHIHRYALTDPAAEQINLLQAHFEQHYLR
Burkholderia_sp. B6_(MN840556)  RQTDSHIHRYALTDPAAEQINLLQAHFEQHYLR
Burkholderia_sp._B7_(MN840551)  RQMHSHIHRYALTDPAAEQINLLQAHFEQHYLR
Burkholderia_sp._B8_(MN840554)  RQTDSHIHRYALTDPAAEQINLLQAHFEQHYLR
B._pseudomallei_(ZP04890375)    EVPETYVASYRMTKEAASALNHYRAQIDLEQML
B._pseudomallei_(ZP02502372)    EVPETYVASYRMTKEAASALNHYRAQIDLEQML
B._pseudomallei_(ZP02494128)    EVPETYVASYRMTKEAASALNHYRAQIDLEQML
X._euvesicatoria_(WP011347306)  NKFASLIVDWVLQAEAAPRLDALRAQAQLLRVA
X._campestris_pv._musacearum_(  NKFASLIVDWVLQAEAAPRLDALRAQSQLLRVA
X._vasicola_pv._vasculorum_(AV  NKFASLIVDWVLQAEAAPRLDALRAQSQLLRVA
X._citri_(ARR12757)             NKFASLIVDWVLQAEAAPHLDALRAQAQLLRVA
X._campestris_(AKS22319)        NKFASMIVDWVLQAEAAPRLDALRAQAQLLRVA
X._oryzae_(AKO19890)            NKFASLIVDWVLQAEAAPRLDALRAQAQLLRVA
X._oryzae_(ACD59124)            NKFASLIVDWVLQAEAAPRLDALRAQAQLLRVA
Burkholderia_sp._(WP013592484)  SGEFSFGERWAIREERVPRLNQYLTLSNQLRAR
Burkholderia_sp._(ADN59567)     SGEFSFGERWVIADEYVPRLNQYLTLSNQLRAR
R._solanacearum_(YP003748288)   P-NQAHYMRWRLGEQERLAMDDYMAAARMAERG
R._solanacearum_(BAH04968)      GPNVSYLYRERLKPAIAREIEGQLDLADMHRAL
R._solanacearum_(BAD42384)      GPNVSYLYRERLKPAIAREIEGQLDLADMHRAL
R._solanacearum_(YP003747289)   QGNLAYGESWKIRPEVTEVLNAYTDEIQLILKC
R._solanacearum_(WP013209172)   QGNLAYGESWKIRPEVTEVLNAYTDEIQLILKC
R._solanacearum_(WP011002069)   AGNLIMGERMHMTDEAARRL-------------
R._solanacearum_(WP003275417)   P-NQAYYMRWRLGTEERLAMDDYLGVAKIAERG
R._solanacearum_(WP003274509)   QGNLAYGESWKIRPEVTEVLNAYTDEILLILKC
R._solanacearum_(EAP71656)      P-NQAHYMRWRLGEQERLAIDDYMAAARMAERG
R._solanacearum_(EAP71039)      P-NQAYYMRWRLGTEERLAMDDYLGVAKIAERG
R._solanacearum_(CBJ39728)      QGNLAYGESWKIRPEVTEVLNAYTDEIQLMLKC
R._solanacearum_(CBJ35355)      P-NQAHYMRWRLGEQERLAMDDYMAAARMAERG
R._solanacearum_(CAD17997)      GPHIRYLLRERLRPAAAREVEGHLDLANLHRAT
R._solanacearum_(BAH47286)      AGNLIMGERMHMTDEAARRL-------------
R._solanacearum_(BAH47283)      QGNLAYGESWKIRPEVTEVLNAYTDEIQLMLKC
R._solanacearum_(BAH04967)      GPHIRYLLRERLRPAAAREVEGHLDLANLHRAT
R._solanacearum_(BAD42389)      P-NQAHYMRWRLGEQERLAMDDYMAAARMADRG
```

**Supplementary Figure 2**. **Confirmation of correct *awr* mutant by PCR**. **a,** Schematic representation of the construction of knock out vectors. **b,** Schematic representation of the confirmation of the successful gene inactivation. The PCR products of the mutant strain correspond to amplicons A and B; whereas products C and D are amplified from the wild-type. The mutants were checked for the absence of contaminating wild-type by amplifying an internal fragment (amplicons **I**) form the wild-type gene of interest. The removal of the knock-out vector was confirmed by amplification of the counter selection marker gene *pheS* (amplicons pheS). **c,** Confirmation of correct *awr* mutant by PCR. PCR products from *B. rhizoxinica* wild-type (B1WT) and mutant (Δ*awr*::Kan$^r$) genomic DNA were obtained using control primers listed in Supplementary Table 5. Bands corresponding to the expected size are indicated by asterisks (*).

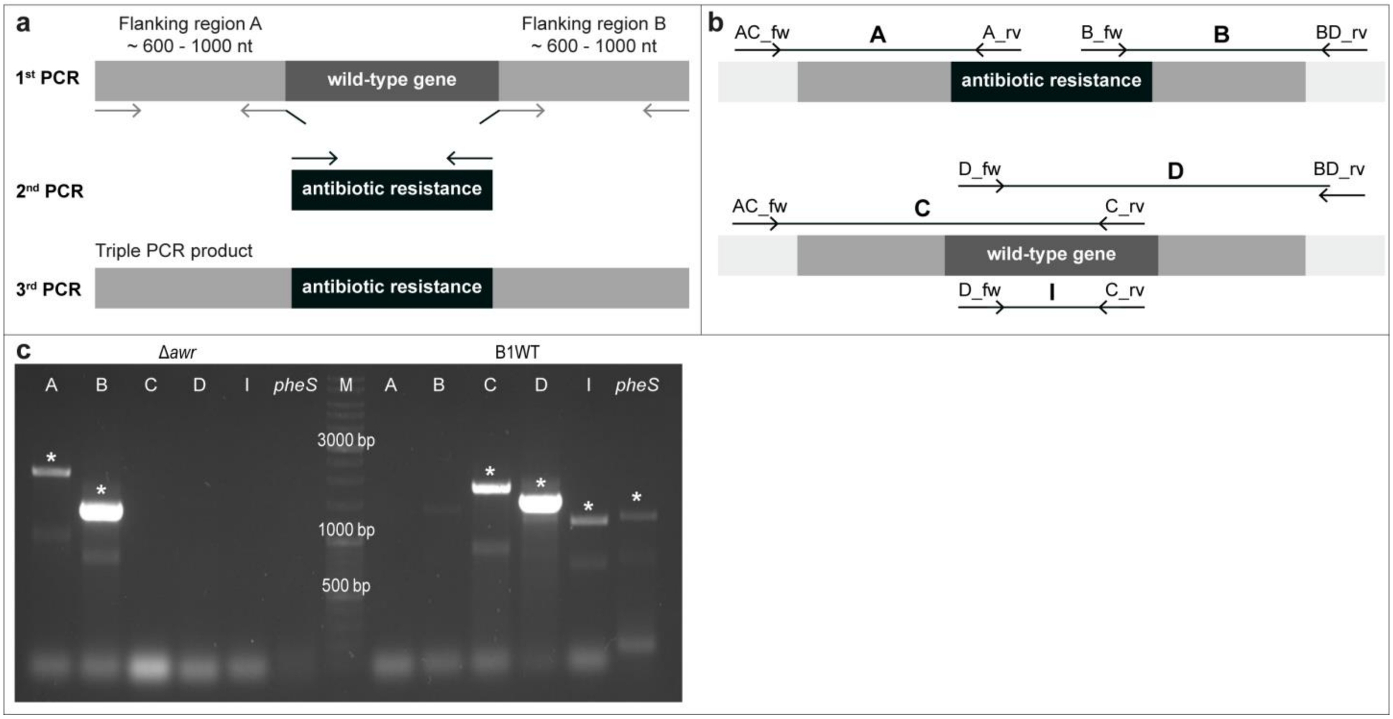

**Supplementary Figure 3.** Alignment of predicted transcription-activator like effector (TALE) proteins from endofungal *Burkholderia* species (BAT: *Burkholderia* transcription-activator like effectors) and plant pathogenic *Ralstonia solanaceraum* and *Xanthomonas* species. Sequences obtained in this study are highlighted in bold. GenBank accession numbers are indicated in brackets.

```
                                             10        20        30        40        50        60        70        80        90       100
                                     ....|....|....|....|....|....|....|....|....|....|....|....|....|....|....|....|....|....|....|....|
B.rhizoxinica_B1_(E5AV36)            MAGNIGGAQALQAVLSLGPALRERGFSQPDIVKIAGNTGGAQALQAVLDLELTLVEHGFSQPDIVRITGNRGGAQALQAVLALELTLRERGFSQPDIVKI
Burkholderia_sp._B2_(MN891944)       MAGNIGGAQALQAVLSLGPALRERGFSQPDIVKIAGNTGGAQALQAVLDLELTLVEHGFSQPDIVRITGNRGGAQALQAVLALELTLRERGFSQPDIVKI
Burkholderia_sp._B4_(MN840538)       MAGNDGGAQALQAVLELEPAFRERGFSQPDIVKMAGNNGGAQALQAGLELEPAFRERGFNPTDMVKIAGNNGGAQALQAVLELEPALRERGFSQPDIVKM
B._endofungorum_B5_(MN891945)        MAGNIGGAQALQAVLSLGPALRERGFSQPDIVKIAGNTGGAQALQAVLDLELTLVERGFSQPDIVRITGNRGGAQALQAVLALELTLRERGFSQPDIVKI
Burkholderia_sp._B6_(MN840537)       MAGNIGGAQALQAVLSLGPALRERGFSQPDIVKIAGNTGGAQALQAVLDLELTLVEHGFSQPDIVRITGNRGGAQALQAVLALELTLRERGFSQPDIVKI
Burkholderia_sp._B7_(MN840539)       MAGNDGGAQALQAVFELEPAFRERSFSQPDIVKMAGNNGGAQALQAVLELDPAFRERGFNPTDIVKIAGNNGGAQALQAVLELEPALRERGFSQPDIVKM
Burkholderia_sp._B7_(MN840540)       IAGNIGGAQALQAVLELEPAFRERGFSQPDIVKMAGNSGGAQALQAGLELEPAFRERGFNPTDIVKIAGNNGGAQALQAVLELEPALRERSFSQPDIVKM
Burkholderia_sp._B8_(MN840541)       MAGNIGGAQALQAVLELEPAFRERGFSQPDIVKMAG----------------------------------------------------------------
Burkholderia_sp._B13_(SIT73265       MAGNDGGAQALQAVLELEPAFRERGFSQPDIVKMAGNNGGAQALQAGLELEPAFRERGFNPTDMVKIAGNNGGAQALQAVLELEPALRERGFSQPDIVKM
Burkholderia_sp._B14_(SIT71710       MAGNDGGAQALQAVFELEPAFRERSFSQPDIVKMAGNNGGAQALQAVLELDPAFRERGFNPTDIVKIAGNNGGAQALQAVLELEPALRERGFSQPDIVKM
Burkholderia_sp._B14_(SIT64981       IAGNIGGAQALQAVLELEPAFRERGFSQPDIVKMAGNSGGAQALQAGLELEPAFRERGFNPTDIVKIAGNNGGAQALQAVLELEPALRERSFSQPDIVKM
B._rhizoxinica_B1_(E5AW45)           IAGNSGGAQALQAVLELEPAFRERGFGQPDIVKMASNIGGAQALQAVLELEPALRERGFSQPDIVEMAGNIGGAQALQAVLELEPAFRERGFSQSDIVKI
Burkholderia_sp._B2_(MN840542)       IAGNSGGAQALQAVLELEPAFRERGFGQPDIVKMASNIGGAQALQAVLELEPALRERGFSQPDIVEMAGNIGGAQALQAVLELEPAFRERGFSQSDIVKI
Burkholderia_sp._B6_(MN840543)       IAGNSGGAQALQAVLELEPAFRERGFGQPDIVKMASNIGGAQALQAVLELEPALRERGFSQPDIVEMAGNIGGAQALQAVLELEPAFRERGFSQSDIVKI
Burkholderia_sp._B3_(MN840545)       IEKHYGGSATLAFISNQHDELAQ-VLSRADILKIASYDCAAQALQAVLDCGPMLGKRGFSRADIVKIAGNGGGAQALQAVLELEPTFRERGFSRGDTVKI
Burkholderia_sp._B7_(MN840544)       IEKHYGGSATLEFISNQHDKLAQ-VLSRADILKIASYDCAAQALQAVLDCGPMLGKRGFSRADIVKIAGNGGGAQALQAVLELEPTFRERGFSRGDTVKI
Burkholderia_sp._B14_(SIT64975       IEKHYGGSATLEFISNQHDKLAQ-VLSRADILKIASYDCAAQALQAVLDCGPMLGKRGFSRADIVKIAGNGGGAQALQAVLELEPTFRERGFSRGDTVKI
B._rhizoxinica_B1_(E5AW43)           IEKHSGGADALEFISNKYDALTQ-VLSRADILKIACHDCAAHALQAVLDYEQVFRQRGFARADIIKITGNGGGAQALKAVVVHGPTLNECGFSQADIVRI
Burkholderia_sp._B2_(MN840547)       IEKHSGGADALEFISNKYDALTQ-VLSRADILKIACHDCAAHALQAVLDYEQVFRQRGFARADIIKITGNGGGAQALKAVVVHGPTLNECGFSQADIVRI
Burkholderia_sp._B6_(MN840546)       IEKHSGGADALEFISNKYDALTQ-VLSRADILKIACHDCAAHALQAVLDYEQVFRQRGFARADIIKITGNGGGAQALKAVVVHGPTLNECGFSQADIVRI
Burkholderia_sp._B4_(MN840549)       ----------------------------------------------VLDYEQVFRQRGFARVDIIKITGNDGGAQALKAVIVHG----------------
Burkholderia_sp._B13_(SIT73247       ----------------------------------------------MLDYEQVFRQRGFARVDIIKITGNDGGAQALKAVIVHG----------------
Burkholderia_sp._B7_(MN840548)       ----------------------------------------------VLDYEQVFRQRGFARVDIIKITGNDGGAQALKAVVVHG----------------
AvrXa10_X._oryzae_(Q56830.1)         QQQEKIKPKVRSTVAQHHEALVGHGFTHAHIVALSQHP-AALGTVAVTYQDIIRALPEATHEDIVGVGKQWSGARALEALLTEAGELRGPPLQLDTGQLL
AvrBs3_X._euvesicatoria_(P1472       QQQEKIKPKVRSTVAQHHEALVGHGFTHAHIVALSQHP-AALGTVAVKYQDMIAALPEATHEAIVGVGKQWSGARALEALLTVAGELRGPPLQLDTGQLL
Brg11_R._solanacearum_(Q8XYE3.       EQIRKLKQESLSEIAKYHTTLTGQGFTHADICRISRRR-QSLRVVARNYPELAAALPELTRAHIVDIARQRSGDLALQALLPVATALTAAPLRLSASQIA
Brg11_R._solanacearum_(Q68A49.       EQIRKLKQESLSEIAKYHTTLTGQGFTHADICRISRRR-QSLRVVARNYPELAAALPELTRAHIVDIARQRSGDLALQALLPVATALTAAPLRLSASQIA
```

```
                                           110       120       130       140       150       160       170       180       190       200
                                  ....|....|....|....|....|....|....|....|....|....|....|....|....|....|....|....|....|....|....|....|
B.rhizoxinica_B1_(E5AV36)         AGNSGGAQALQ---AVLDLELTFRERGFSQADIVKIAG--------------------------------------------------------------
Burkholderia_sp._B2_(MN891944)    AGNSGGAQALQ---AVLDLELTFRERGFSQADIVKIAG--------------------------------------------------------------
Burkholderia_sp._B4_(MN840538)    ASNSGGAQALQ---VVLELEPTLRERDFRQADIVKIAS--------------------------------------------------------------
B._endofungorum_B5_(MN891945)     AGNSGGAQALH---AVLDLELTFRERGFSQADIVKIAG--------------------------------------------------------------
Burkholderia_sp._B6_(MN840537)    AGNSGGAQALQ---AVLDLELTFRERGFSQADIVKIAG--------------------------------------------------------------
Burkholderia_sp._B7_(MN840539)    ASNSGGAQALQ---AGLELEPTLRERDFRQADIVKMAS--------------------------------------------------------------
Burkholderia_sp._B7_(MN840540)    ASNSGGAQALQ---AVLVLEPTLRERDFRQADIVKMAS--------------------------------------------------------------
Burkholderia_sp._B8_(MN840541)    --NIGGAQALQ---AVLELEPMLRECDFRQADIVKIAG--------------------------------------------------------------
Burkholderia_sp._B13_(SIT73265    ASNSGGAQALQ---VVLELEPTLRERDFRQADIVKIAS--------------------------------------------------------------
Burkholderia_sp._B14_(SIT71710    ASNSGGAQALQ---AGLELEPTLRERDFRQADIVKMAS--------------------------------------------------------------
Burkholderia_sp._B14_(SIT64981    ASNSGGAQALQ---AVLVLEPTLRERDFRQADIVKMAS--------------------------------------------------------------
B._rhizoxinica_B1_(E5AW45)        AGNIGGAQALQ---AVLELEPTLRESDFRQADIVNIAG--------------------------------------------------------------
Burkholderia_sp._B2_(MN840542)    AGNIGGAQALQ---AVLELEPTLRESDFRQADIVNIAG--------------------------------------------------------------
Burkholderia_sp._B6_(MN840543)    AGNIGGAQALQ---AVLELEPTLRESDFRQADIVNIAG--------------------------------------------------------------
Burkholderia_sp._B3_(MN840545)    ASNNGGAQALQALQAVLELEPTLREHGFSRADIVRIAG--------------------------------------------------------------
Burkholderia_sp._B7_(MN840544)    AGNGGGAQALP---AVLELEPTLREHGFSRADIVRIAG--------------------------------------------------------------
Burkholderia_sp._B14_(SIT64975    AGNGGGAQALP---AVLELEPTLREHGFSRADIVRIAG--------------------------------------------------------------
B._rhizoxinica_B1_(E5AW43)        ADNIGGAQALK---AVLEHGPTLNERDYSGADIVKIAG--------------------------------------------------------------
Burkholderia_sp._B2_(MN840547)    ADNIGGAQALK---AVLEHGPTLNERDYSGADIVKIAG--------------------------------------------------------------
Burkholderia_sp._B6_(MN840546)    ADNIGGAQALK---AVLEHGPTLNERDYSGADIVKIAG--------------------------------------------------------------
Burkholderia_sp._B4_(MN840549)    --------------------STLNERGYSGANIVKIAG--------------------------------------------------------------
Burkholderia_sp._B13_(SIT73247    --------------------STLNERGYSGANIVKIAG--------------------------------------------------------------
Burkholderia_sp._B7_(MN840548)    --------------------PTLNERGYSGADIVKIAG--------------------------------------------------------------
AvrXa10_X._oryzae_(Q56830.1)      KIAKRGGVTAVEAVHAWRNALTGAPLNLTPDQVVAIASNIGGNQALETVQRLLPVLCQAHG-LTPDQVVAIASHGGGKQALETVQR-LLPVLCQAHGLTP
AvrBs3_X._euvesicatoria_(P1472    KIAKRGGVTAVEAVHAWRNALTGAPLNLTPEQVVAIASHDGGKQALETVQRLLPVLCQAHG-LTPQQVVAIASNGGGKQALETVQR-LLPVLCQAHGLTP
Brg11_R._solanacearum_(Q8XYE3.    TVAQYGERPAIQALYRLRRKLTRAPLHLTPQQVVAIASNTGGKRALEAVCVQLPVLRAAPYRLSTEQVVAIASNKGGKQALEAVKAHLLDLLGAPYVLDT
Brg11_R._solanacearum_(Q68A49.    TVAQYGERPAIQALYRLRRKLTRAPLHLTPQQVVAIAS--------------------------------------------------------------

                                           210       220       230       240       250       260       270       280       290       300
                                  ....|....|....|....|....|....|....|....|....|....|....|....|....|....|....|....|....|....|....|....|
B.rhizoxinica_B1_(E5AV36)         ----------------------------------------------------------------------------------------------------
Burkholderia_sp._B2_(MN891944)    ----------------------------------------------------------------------------------------------------
Burkholderia_sp._B4_(MN840538)    ----------------------------------------------------------------------------------------------------
B._endofungorum_B5_(MN891945)     ----------------------------------------------------------------------------------------------------
Burkholderia_sp._B6_(MN840537)    ----------------------------------------------------------------------------------------------------
Burkholderia_sp._B7_(MN840539)    ----------------------------------------------------------------------------------------------------
Burkholderia_sp._B7_(MN840540)    ----------------------------------------------------------------------------------------------------
Burkholderia_sp._B8_(MN840541)    ----------------------------------------------------------------------------------------------------
Burkholderia_sp._B13_(SIT73265    ----------------------------------------------------------------------------------------------------
Burkholderia_sp._B14_(SIT71710    ----------------------------------------------------------------------------------------------------
Burkholderia_sp._B14_(SIT64981    ----------------------------------------------------------------------------------------------------
B._rhizoxinica_B1_(E5AW45)        ----------------------------------------------------------------------------------------------------
Burkholderia_sp._B2_(MN840542)    ----------------------------------------------------------------------------------------------------
Burkholderia_sp._B6_(MN840543)    ----------------------------------------------------------------------------------------------------
Burkholderia_sp._B3_(MN840545)    ----------------------------------------------------------------------------------------------------
Burkholderia_sp._B7_(MN840544)    ----------------------------------------------------------------------------------------------------
Burkholderia_sp._B14_(SIT64975    ----------------------------------------------------------------------------------------------------
B._rhizoxinica_B1_(E5AW43)        ----------------------------------------------------------------------------------------------------
Burkholderia_sp._B2_(MN840547)    ----------------------------------------------------------------------------------------------------
Burkholderia_sp._B6_(MN840546)    ----------------------------------------------------------------------------------------------------
Burkholderia_sp._B4_(MN840549)    ----------------------------------------------------------------------------------------------------
Burkholderia_sp._B13_(SIT73247    ----------------------------------------------------------------------------------------------------
Burkholderia_sp._B7_(MN840548)    ----------------------------------------------------------------------------------------------------
AvrXa10_X._oryzae_(Q56830.1)      DQVVAIASNIGGKQALATVQR-LLPVLCQDHGLTPDQVVAIASHGGGKQALETVQR-LLPVLCQDHGLTPDQVVAIASNIGGKQALETVQRLLP-VLCQD
AvrBs3_X._euvesicatoria_(P1472    QQVVAIASNSGGKQALETVQR-LLPVLCQAHGLTPEQVVAIASNGGGKQALETVQR-LLPVLCQAHGLTPEQVVAIASNIGGKQALETVQALLP-VLCQA
Brg11_R._solanacearum_(Q8XYE3.    EQVVAIASHNGGKQALEAVKADLLDLRGAPYALSTEQVVAIASHNGGKQALEAVKADLLELRGAPYALSTEQVVAIASHNGGKQALEAVKAHLLDLRGVP
Brg11_R._solanacearum_(Q68A49.    ----------------------------------------------------------------------------------------------------
```

```
                                        310       320       330       340       350       360       370       380       390       400
                                ....|....|....|....|....|....|....|....|....|....|....|....|....|....|....|....|....|....|....|....|
B.rhizoxinica_B1_(E5AV36)       ----------------------------------------------------------------------------------------------------
Burkholderia_sp._B2_(MN891944)  ----------------------------------------------------------------------------------------------------
Burkholderia_sp._B4_(MN840538)  ----------------------------------------------------------------------------------------------------
B._endofungorum_B5_(MN891945)   ----------------------------------------------------------------------------------------------------
Burkholderia_sp._B6_(MN840537)  ----------------------------------------------------------------------------------------------------
Burkholderia_sp._B7_(MN840539)  ----------------------------------------------------------------------------------------------------
Burkholderia_sp._B7_(MN840540)  ----------------------------------------------------------------------------------------------------
Burkholderia_sp._B8_(MN840541)  ----------------------------------------------------------------------------------------------------
Burkholderia_sp._B13_(SIT73265  ----------------------------------------------------------------------------------------------------
Burkholderia_sp._B14_(SIT71710  ----------------------------------------------------------------------------------------------------
Burkholderia_sp._B14_(SIT64981  ----------------------------------------------------------------------------------------------------
B._rhizoxinica_B1_(E5AW45)      ----------------------------------------------------------------------------------------------------
Burkholderia_sp._B2_(MN840542)  ----------------------------------------------------------------------------------------------------
Burkholderia_sp._B6_(MN840543)  ----------------------------------------------------------------------------------------------------
Burkholderia_sp._B3_(MN840545)  ----------------------------------------------------------------------------------------------------
Burkholderia_sp._B7_(MN840544)  ----------------------------------------------------------------------------------------------------
Burkholderia_sp._B14_(SIT64975  ----------------------------------------------------------------------------------------------------
B._rhizoxinica_B1_(E5AW43)      ----------------------------------------------------------------------------------------------------
Burkholderia_sp._B2_(MN840547)  ----------------------------------------------------------------------------------------------------
Burkholderia_sp._B6_(MN840546)  ----------------------------------------------------------------------------------------------------
Burkholderia_sp._B4_(MN840549)  ----------------------------------------------------------------------------------------------------
Burkholderia_sp._B13_(SIT73247  ----------------------------------------------------------------------------------------------------
Burkholderia_sp._B7_(MN840548)  ----------------------------------------------------------------------------------------------------
AvrXa10_X._oryzae_(Q56830.1)    HGLTPDQVVAIASNIGGKQALETVQRLLPVLCQDHG-LTPDQVVAIASNNGGKQALETVQRLLP-VLCQTHGLTPDQVVAIANHDGGKQALETVQR-LLP
AvrBs3_X._euvesicatoria_(P1472  HGLTPEQVVAIASNIGGKQALETVQALLPVLCQAHG-LTPEQVVAIASNIGGKQALETVQALLP-VLCQAHGLTPEQVVAIASHDGGKQALETVQR-LLP
Brg11_R._solanacearum_(Q8XYE3.  YALSTEQVVAIASHNGGKQALEAVKAQLLDLRGAPYALSTAQVVAIASNGGGKQALEGIGEQLLKLRTAPYGLSTEQVVAIASHDGGKQALEAVGAQLVA
Brg11_R._solanacearum_(Q68A49.  ----------------------------------------------------------------------------------------------------

                                        410       420       430       440       450       460       470       480       490       500
                                ....|....|....|....|....|....|....|....|....|....|....|....|....|....|....|....|....|....|....|....|
B.rhizoxinica_B1_(E5AV36)       ----------------------------------------------------------------------------------------------------
Burkholderia_sp._B2_(MN891944)  ----------------------------------------------------------------------------------------------------
Burkholderia_sp._B4_(MN840538)  ----------------------------------------------------------------------------------------------------
B._endofungorum_B5_(MN891945)   ----------------------------------------------------------------------------------------------------
Burkholderia_sp._B6_(MN840537)  ----------------------------------------------------------------------------------------------------
Burkholderia_sp._B7_(MN840539)  ----------------------------------------------------------------------------------------------------
Burkholderia_sp._B7_(MN840540)  ----------------------------------------------------------------------------------------------------
Burkholderia_sp._B8_(MN840541)  ----------------------------------------------------------------------------------------------------
Burkholderia_sp._B13_(SIT73265  ----------------------------------------------------------------------------------------------------
Burkholderia_sp._B14_(SIT71710  ----------------------------------------------------------------------------------------------------
Burkholderia_sp._B14_(SIT64981  ----------------------------------------------------------------------------------------------------
B._rhizoxinica_B1_(E5AW45)      ----------------------------------------------------------------------------------------------------
Burkholderia_sp._B2_(MN840542)  ----------------------------------------------------------------------------------------------------
Burkholderia_sp._B6_(MN840543)  ----------------------------------------------------------------------------------------------------
Burkholderia_sp._B3_(MN840545)  ----------------------------------------------------------------------------------------------------
Burkholderia_sp._B7_(MN840544)  ----------------------------------------------------------------------------------------------------
Burkholderia_sp._B14_(SIT64975  ----------------------------------------------------------------------------------------------------
B._rhizoxinica_B1_(E5AW43)      ----------------------------------------------------------------------------------------------------
Burkholderia_sp._B2_(MN840547)  ----------------------------------------------------------------------------------------------------
Burkholderia_sp._B6_(MN840546)  ----------------------------------------------------------------------------------------------------
Burkholderia_sp._B4_(MN840549)  ----------------------------------------------------------------------------------------------------
Burkholderia_sp._B13_(SIT73247  ----------------------------------------------------------------------------------------------------
Burkholderia_sp._B7_(MN840548)  ----------------------------------------------------------------------------------------------------
AvrXa10_X._oryzae_(Q56830.1)    VLCQDHGLTPDQVVAIASNIGGKQALATVQR-LLPVLCQAHGLTPDQVVAIASHDGGKQALETVQR-LLPVLCQDHGLTPDQVVAIASN-----------
AvrBs3_X._euvesicatoria_(P1472  VLCQAHGLTPEQVVAIASHDGGKQALETVQR-LLPVLCQAHGLTPQQVVAIASNGGGKQALETVQR-LLPVLCQAHGLTPEQVVAIASNSGGKQALETVQ
Brg11_R._solanacearum_(Q8XYE3.  LRAAPYALSTEQVVAIASNKGGKQALEAVKAQLLELRGAPYALSTAQVVAIASHDGGNQALEAVGTQLVALRAAPYALSTEQVVAIASHDGGKQALEAVG
Brg11_R._solanacearum_(Q68A49.  ----------------------------------------------------------------------------------------------------
```

```
                                          510       520       530       540       550       560       570       580       590       600
                                 ....|....|....|....|....|....|....|....|....|....|....|....|....|....|....|....|....|....|....|....|
B.rhizoxinica_B1_(E5AV36)        ----------------------------------------------------------------------------------------------------
Burkholderia_sp._B2_(MN891944)   ----------------------------------------------------------------------------------------------------
Burkholderia_sp._B4_(MN840538)   ----------------------------------------------------------------------------------------------------
B._endofungorum_B5_(MN891945)    ----------------------------------------------------------------------------------------------------
Burkholderia_sp._B6_(MN840537)   ----------------------------------------------------------------------------------------------------
Burkholderia_sp._B7_(MN840539)   ----------------------------------------------------------------------------------------------------
Burkholderia_sp._B7_(MN840540)   ----------------------------------------------------------------------------------------------------
Burkholderia_sp._B8_(MN840541)   ----------------------------------------------------------------------------------------------------
Burkholderia_sp._B13_(SIT73265   ----------------------------------------------------------------------------------------------------
Burkholderia_sp._B14_(SIT71710   ----------------------------------------------------------------------------------------------------
Burkholderia_sp._B14_(SIT64981   ----------------------------------------------------------------------------------------------------
B._rhizoxinica_B1_(E5AW45)       ----------------------------------------------------------------------------------------------------
Burkholderia_sp._B2_(MN840542)   ----------------------------------------------------------------------------------------------------
Burkholderia_sp._B6_(MN840543)   ----------------------------------------------------------------------------------------------------
Burkholderia_sp._B3_(MN840545)   ----------------------------------------------------------------------------------------------------
Burkholderia_sp._B7_(MN840544)   ----------------------------------------------------------------------------------------------------
Burkholderia_sp._B14_(SIT64975   ----------------------------------------------------------------------------------------------------
B._rhizoxinica_B1_(E5AW43)       ----------------------------------------------------------------------------------------------------
Burkholderia_sp._B2_(MN840547)   ----------------------------------------------------------------------------------------------------
Burkholderia_sp._B6_(MN840546)   ----------------------------------------------------------------------------------------------------
Burkholderia_sp._B4_(MN840549)   ----------------------------------------------------------------------------------------------------
Burkholderia_sp._B13_(SIT73247   ----------------------------------------------------------------------------------------------------
Burkholderia_sp._B7_(MN840548)   ----------------------------------------------------------------------------------------------------
AvrXa10_X._oryzae_(Q56830.1)     ---------------------------------------------------------NGGKQALETVQRLLPVLCQDHG-LTPAQVVAIANHGGGKQALE
AvrBs3_X._euvesicatoria_(P1472   ALLPVLCQAHGLTPEQVVAIASNSGGKQALETVQRLLPVLCQAHGLTPEQVVAIASHDGGKQALETVQRLLPVLCQAHG-LTPEQVVAIASHDGGKQALE
Brg11_R._solanacearum_(Q8XYE3.   AQLVALRAA---------------------------------PYALNTEQVVAIASSHGGKQALEAVRALFPDLRAAPYALSTAQLVAIASNPGGKQALE
Brg11_R._solanacearum_(Q68A49.   ----------------------------------------------------------------------------------------------------

                                          610       620       630       640       650       660       670       680       690       700
                                 ....|....|....|....|....|....|....|....|....|....|....|....|....|....|....|....|....|....|....|....|
B.rhizoxinica_B1_(E5AV36)        ------------------------------------------------------------------------------------------------NDGG
Burkholderia_sp._B2_(MN891944)   ------------------------------------------------------------------------------------------------NDGG
Burkholderia_sp._B4_(MN840538)   ------------------------------------------------------------------------------------------------NGGS
B._endofungorum_B5_(MN891945)    ------------------------------------------------------------------------------------------------NDGG
Burkholderia_sp._B6_(MN840537)   ------------------------------------------------------------------------------------------------NDGG
Burkholderia_sp._B7_(MN840539)   ------------------------------------------------------------------------------------------------NGGS
Burkholderia_sp._B7_(MN840540)   ------------------------------------------------------------------------------------------------NGGS
Burkholderia_sp._B8_(MN840541)   ------------------------------------------------------------------------------------------------NGGS
Burkholderia_sp._B13_(SIT73265   ------------------------------------------------------------------------------------------------NGGS
Burkholderia_sp._B14_(SIT71710   ------------------------------------------------------------------------------------------------NGGS
Burkholderia_sp._B14_(SIT64981   ------------------------------------------------------------------------------------------------NGGS
B._rhizoxinica_B1_(E5AW45)       ------------------------------------------------------------------------------------------------NDGS
Burkholderia_sp._B2_(MN840542)   ------------------------------------------------------------------------------------------------NDGS
Burkholderia_sp._B6_(MN840543)   ------------------------------------------------------------------------------------------------NDGS
Burkholderia_sp._B3_(MN840545)   ------------------------------------------------------------------------------------------------NGGG
Burkholderia_sp._B7_(MN840544)   ------------------------------------------------------------------------------------------------NGGG
Burkholderia_sp._B14_(SIT64975   ------------------------------------------------------------------------------------------------NGGG
B._rhizoxinica_B1_(E5AW43)       ------------------------------------------------------------------------------------------------NGGG
Burkholderia_sp._B2_(MN840547)   ------------------------------------------------------------------------------------------------NGGG
Burkholderia_sp._B6_(MN840546)   ------------------------------------------------------------------------------------------------NGGG
Burkholderia_sp._B4_(MN840549)   ------------------------------------------------------------------------------------------------NGGG
Burkholderia_sp._B13_(SIT73247   ------------------------------------------------------------------------------------------------NGGG
Burkholderia_sp._B7_(MN840548)   ------------------------------------------------------------------------------------------------NGGG
AvrXa10_X._oryzae_(Q56830.1)     TVQRLLPVLCQDHG-LTPVQVVAIASNSGGKQALETVQRLLP-VLCQDHGLTPVQVVAIASNGGGKQALATVQRLLPVLCQDHG-LTPVQVVAIASHDGG
AvrBs3_X._euvesicatoria_(P1472   TVQRLLPVLCQAHG-LTPEQVVAIASHDGGKQALETVQRLLP-VLCQAHGLTPQQVVAIASNGGGRPALETVQRLLPVLCQAHG-LTPEQVVAIASHDGG
Brg11_R._solanacearum_(Q8XYE3.   AVRALFRELRAAPYALSTEQVVAIASNHGGKQALEAVRALFRGLRAAPYGLSTAQVVAIASSNGGKQALEAVWALLPVLRATPYDLNTAQIVAIASHDGG
Brg11_R._solanacearum_(Q68A49.   ------------------------------------------------------------------------------------------------HDGG
```

```
                                         710       720       730       740       750       760       770       780       790       800
                                ....|....|....|....|....|....|....|....|....|....|....|....|....|....|....|....|....|....|....|....|
B.rhizoxinica_B1_(E5AV36)       TQALHAVLDLERMLG--ERGFSRADIVNVAGNNGGAQALKAVLEHEATLNERGFSRADIVKIAGNGGGAQALKAVLE-----------------------
Burkholderia_sp._B2_(MN891944)  TQALHAVLDLERMLG--ERGFSRADIVNVAGNNGGAQALKAVLEHEATLNERGFSRADIVKIAGNGGGAQALKAVLE-----------------------
Burkholderia_sp._B4_(MN840538)  AQALNAVIKLGPTLR--QRGFSRADIVKIAANGGGAQALQAVLKHGPTLGERGFTLTDIVEMAGNNGGAQALKAVLE-----------------------
B._endofungorum_B5_(MN891945)   TQALHAVLDLERMLG--ERGFSRADIVNVAGNNGGAQALKAVLEHEATLNERGFSRADIVKIAGNGGGAQALKAVLE-----------------------
Burkholderia_sp._B6_(MN840537)  TQALHAVLDLERMLG--ERGFSRADIVNVAGNNGGAQALKAVLEHEATLNERGFSRADIVKIAGNGGGAQALKAVLE-----------------------
Burkholderia_sp._B7_(MN840539)  AQALNAVIKLGPTLR--QRGFSRADIVKIAANGGGAQALQAVLKHGPTLGER------------------------------------------------
Burkholderia_sp._B7_(MN840540)  AQALNAVIKLGPTLR--QRGFSRADIVKIAANGGGAQALQAVLKHGPTLGERGFTLTDIVKMAGNNGGAQALKAVLE-----------------------
Burkholderia_sp._B8_(MN840541)  AQALKAVLEHGPTLR--QRGFSRADIVKIAANGGGAQALQAVLKHG------------------------------------------------------
Burkholderia_sp._B13_(SIT73265  AQALNAVIKLGPTLR--QRGFSRADIVKIAANGGGAQALQAVLKHGPTLGERGFTLTDIVEMAGNNGGAQALKAVLE-----------------------
Burkholderia_sp._B14_(SIT71710  AQALNAVIKLGPTLR--QRGFSRADIVKIAANGGGAQALQAVLKHGPTLGER------------------------------------------------
Burkholderia_sp._B14_(SIT64981  AQALNAVIKLGPTLR--QRGFSRADIVKIAANGGGAQALQAVLKHGPTLGERGFTLTDIVKMAGNNGGAQALKAVLE-----------------------
B._rhizoxinica_B1_(E5AW45)      TQALKAVIEHGPRLR--QRGFNRASIVKIAGNSGGAQALQAVLKHGPTLDERGFNLTNIVKIAGNGGGAQALKAVIE-----------------------
Burkholderia_sp._B2_(MN840542)  TQALKAVIEHGPRLR--QRGFNRASIVKIAGNSGGAQALQAVLKHGPTLDERGFNLTNIVKIAGNGGGAQALKAVIE-----------------------
Burkholderia_sp._B6_(MN840543)  TQALKAVIEHGPRLR--QRGFNRASIVKIAGNSGGAQALQAVLKHGPTLDERGFNLTNIVKIAGNGGGAQALKAVIE-----------------------
Burkholderia_sp._B3_(MN840545)  AQALYSVLDVGLTLG--KRSFSRADIVKMARNDGGAQALHTVLKLEPTLGERGFSRGDIVKMAGNIG---------------------------------
Burkholderia_sp._B7_(MN840544)  AQALYSVLDVGPTLG--KRGFSRADIVKIAGNGGGAQALHTVFKLEPTLGERGFSRGDIVKMAGNIGGAQALQAVLELEPAFHERSFCQPDIVKMAGNIG
Burkholderia_sp._B14_(SIT64975  AQALYSVLDVGPTLG--KRGFSRADIVKIAGNGGGAQALHTVFKLEPTLGERGFSRGDIVKMAGNIGGAQALQAVLELEPAFHERSFCQPDIVKMAGNIG
B._rhizoxinica_B1_(E5AW43)      ARALKAVVMHGPTLC--ESGYSGADIVKIASNGGGAQALE------------------------------------------------------------
Burkholderia_sp._B2_(MN840547)  ARALKAVVMHGPTLC--ESGYSGADIVKIASNGGGAQALE------------------------------------------------------------
Burkholderia_sp._B6_(MN840546)  ARALKAVVMHGPTLC--ESGYSGADIVKIASNGGGAQALE------------------------------------------------------------
Burkholderia_sp._B4_(MN840549)  ARAIEAVVMHG-----------------------------------------------------------------------------------------
Burkholderia_sp._B13_(SIT73247  ARAIEAVVMHG-----------------------------------------------------------------------------------------
Burkholderia_sp._B7_(MN840548)  AWALKAVVMHG-----------------------------------------------------------------------------------------
AvrXa10_X._oryzae_(Q56830.1)    KQALETVQRLLPVLC-QDHGLTPDQVVAIASNGG-KQALESIVAQLSRPDPALAALTNDHLVALACLGGRPALDAVKKGLP-------------------
AvrBs3_X._euvesicatoria_(P1472  KQALETVQRLLPVLC-QAHGLTPQQVVAIASNGGGRPALESIVAQLSRPDPALAALTNDHLVALACLGGRPALDAVKKGLP-------------------
Brg11_R._solanacearum_(Q8XYE3.  KPALEAVWAKLPVLRGAPYALSTAQVVAIACISG-QQALEAIEAHMPTLRQASHSLSPERVAAIACIGGRSAVEAVRQGLPV------------------
Brg11_R._solanacearum_(Q68A49.  KPALEAVWAKLPVLRGVPYALSTAQVVAIACISG-QQALEAIEAHMPTLRQAPHSLSPERVAAIACIGGRSAVEAVRQGLPV------------------

                                         810       820       830       840       850       860       870       880       890       900
                                ....|....|....|....|....|....|....|....|....|....|....|....|....|....|....|....|....|....|....|....|
B.rhizoxinica_B1_(E5AV36)       ----------------------------------------------------------------------------------------------------
Burkholderia_sp._B2_(MN891944)  ----------------------------------------------------------------------------------------------------
Burkholderia_sp._B4_(MN840538)  ----------------------------------------------------------------------------------------------------
B._endofungorum_B5_(MN891945)   ----------------------------------------------------------------------------------------------------
Burkholderia_sp._B6_(MN840537)  ----------------------------------------------------------------------------------------------------
Burkholderia_sp._B7_(MN840539)  ----------------------------------------------------------------------------------------------------
Burkholderia_sp._B7_(MN840540)  ----------------------------------------------------------------------------------------------------
Burkholderia_sp._B8_(MN840541)  ----------------------------------------------------------------------------------------------------
Burkholderia_sp._B13_(SIT73265  ----------------------------------------------------------------------------------------------------
Burkholderia_sp._B14_(SIT71710  ----------------------------------------------------------------------------------------------------
Burkholderia_sp._B14_(SIT64981  ----------------------------------------------------------------------------------------------------
B._rhizoxinica_B1_(E5AW45)      ----------------------------------------------------------------------------------------------------
Burkholderia_sp._B2_(MN840542)  ----------------------------------------------------------------------------------------------------
Burkholderia_sp._B6_(MN840543)  ----------------------------------------------------------------------------------------------------
Burkholderia_sp._B3_(MN840545)  ----------------------------------------------------------------------------------------------------
Burkholderia_sp._B7_(MN840544)  GAQALQAVLELEPAFRERGFSRGDIVKMAGNIGGAQALQAGLELEPAFRERGFSRGDIVKMAGNIGGAQALQAVLELEPAFREHGFSQPDIVKIAGNIGG
Burkholderia_sp._B14_(SIT64975  GAQALQAVLELEPAFRERGFSRGDIVKMAGNIGGAQALQAGLELEPAFRERGFSRGDIVKMAGNIGGAQALQAVLELEPAFREHGFSQPDIVKIAGNIGG
B._rhizoxinica_B1_(E5AW43)      ----------------------------------------------------------------------------------------------------
Burkholderia_sp._B2_(MN840547)  ----------------------------------------------------------------------------------------------------
Burkholderia_sp._B6_(MN840546)  ----------------------------------------------------------------------------------------------------
Burkholderia_sp._B4_(MN840549)  ----------------------------------------------------------------------------------------------------
Burkholderia_sp._B13_(SIT73247  ----------------------------------------------------------------------------------------------------
Burkholderia_sp._B7_(MN840548)  ----------------------------------------------------------------------------------------------------
AvrXa10_X._oryzae_(Q56830.1)    ----------------------------------------------------------------------------------------------------
AvrBs3_X._euvesicatoria_(P1472  ----------------------------------------------------------------------------------------------------
Brg11_R._solanacearum_(Q8XYE3.  ----------------------------------------------------------------------------------------------------
Brg11_R._solanacearum_(Q68A49.  ----------------------------------------------------------------------------------------------------
```

```
                                        910       920       930       940       950       960       970       980       990      1000
                                ....|....|....|....|....|....|....|....|....|....|....|....|....|....|....|....|....|....|....|....|
B.rhizoxinica_B1_(E5AV36)       ---------------------------------------------------------------------------HEATLDERGFSRADIVRIAGNGGGA
Burkholderia_sp._B2_(MN891944)  ---------------------------------------------------------------------------HEATLDERGFSRADIVRIAGNGGGA
Burkholderia_sp._B4_(MN840538)  ---------------------------------------------------------------------------HGSTLDERGFTLTDIVKMAGNNGGA
B._endofungorum_B5_(MN891945)   ---------------------------------------------------------------------------HEATLDERGFSRADIVRIAGNGGGA
Burkholderia_sp._B6_(MN840537)  ---------------------------------------------------------------------------HEATLDERGFSRADIVRIAGNGGGA
Burkholderia_sp._B7_(MN840539)  -----------------------------------------------------------------------------------GFTLTDIVKMASNNGGA
Burkholderia_sp._B7_(MN840540)  ---------------------------------------------------------------------------HGPTLDERGFTLTDIVKMAGNNGGA
Burkholderia_sp._B8_(MN840541)  ----------------------------------------------------------------------------------------------------
Burkholderia_sp._B13_(SIT73265  ---------------------------------------------------------------------------HGSTLDERGFTLTDIVKMAGNNGGA
Burkholderia_sp._B14_(SIT71710  -----------------------------------------------------------------------------------GFTLTDIVKMASNNGGA
Burkholderia_sp._B14_(SIT64981  ---------------------------------------------------------------------------HGPTLDERGFTLTDIVKMAGNNGGA
B._rhizoxinica_B1_(E5AW45)      ---------------------------------------------------------------------------HGPTLQQRGFNLTDIVEMAGKGGGA
Burkholderia_sp._B2_(MN840542)  ---------------------------------------------------------------------------HGPTLQQRGFNLTDIVEMAGKGGGA
Burkholderia_sp._B6_(MN840543)  ---------------------------------------------------------------------------HGPTLQQRGFNLTDIVEMAGNGGGA
Burkholderia_sp._B3_(MN840545)  --------------------------------------------------------------------------------------------------GA
Burkholderia_sp._B7_(MN840544)  AQALQAVLELEPAFRERGFSQSNIVKIAGNIGGAQALQAVLELEPMLRECDFRQTDIVKMAGSGGSAQALNAVIKHGPTLRQRGFSQADIVKMAGNNGGA
Burkholderia_sp._B14_(SIT64975  AQALQAVLELEPAFRERGFSQSNIVKIAGNIGGAQALQAVLELEPMLRECDFRQTDIVKMAGSGGSAQALNAVIKHGPTLRQRGFSQADIVKMAGNNGGA
B._rhizoxinica_B1_(E5AW43)      ----------------------------------------------------------------------------------------------------
Burkholderia_sp._B2_(MN840547)  ----------------------------------------------------------------------------------------------------
Burkholderia_sp._B6_(MN840546)  ----------------------------------------------------------------------------------------------------
Burkholderia_sp._B4_(MN840549)  ----------------------------------------------------------------------------------------------------
Burkholderia_sp._B13_(SIT73247  ----------------------------------------------------------------------------------------------------
Burkholderia_sp._B7_(MN840548)  ----------------------------------------------------------------------------------------------------
AvrXa10_X._oryzae_(Q56830.1)    ---------------------------------------------------------------------------HAPELIRRINRRIPERTSHRVADL-
AvrBs3_X._euvesicatoria_(P1472  ---------------------------------------------------------------------------HAPALIKRTNRRIPERTSHRVADH-
Brg11_R._solanacearum_(Q8XYE3.  ---------------------------------------------------------------------------KAIRRIRREKAPVAGPPPASLGPTP
Brg11_R._solanacearum_(Q68A49.  ---------------------------------------------------------------------------KAIRRIRREKAPVAGPPPASLGPTP

                                       1010      1020      1030      1040      1050      1060      1070      1080      1090      1100
                                ....|....|....|....|....|....|....|....|....|....|....|....|....|....|....|....|....|....|....|....|
B.rhizoxinica_B1_(E5AV36)       QALKAVLEHGPTLNERGFNLTDIVEMAANSGGAQALKAVLEHGPTLRQRGLSLIDIVEIASN-GGAQALK--------AVLKYGPVLMQAGRSNEEIVHV
Burkholderia_sp._B2_(MN891944)  QALKAVLEHGPTLNERGFNLTDIVEMAANSGGAQALKAVLEHGPTLRQRGLSLIDIVEIASN-GGAQALK--------AVLKYGPVLMQAGRSNEEIVHV
Burkholderia_sp._B4_(MN840538)  QALKAVLEHGPTLQQRGFTLTDIVKMAGNIGGAQALKAVLEHGPTLRQRDLSLIDIVEIASN-GGAQALK--------AVLKYGPVLTQAGRSNEEIVHV
B._endofungorum_B5_(MN891945)   QALKAVLEHGPTLNERGFNLTDIVEMAANSGGAQALKAVLEHGPTLRQRGLSLIDIVEIASN-GGAQALK--------AVLKYGPVLMQAGRSNEEIVHV
Burkholderia_sp._B6_(MN840537)  QALKAVLEHGPTLNERGFNLTDIVEMAANSGGAQALKAVLEHGPTLRQRGLSLIDIVEIASN-GGAQALK--------AVLKYGPVLMQAGRSNEEIVHV
Burkholderia_sp._B7_(MN840539)  QALKAVLEHGPTLDERGFTLTDIVKMAGNIGGAQALKTVLGHGPTLRQRDLSLIDIVEIASN-GGAQALK--------AVLKYGPVLTQAGRSNEEIVHV
Burkholderia_sp._B7_(MN840540)  QALKAVLEHGPTLDERGFTLTDIVKMAGNIGGAQALKVVLEHGPTLRQRDLSLIDIVEIASN-GGAQALK--------AVLKYGPVLIQAGRSNEEIVQV
Burkholderia_sp._B8_(MN840541)  ----------PTLDERGFTLTDIVKMAGNVGGAQALKAVLEHGPTLRQRDLSLIDIVEIASN-GGAQALK--------AVLKYGPVLTQVGRSNEEIVNV
Burkholderia_sp._B13_(SIT73265  QALKAVLEHGPTLQQRGFTLTDIVKMAGNIGGAQALKAVLEHGPTLRQRDLSLIDIVEIASN-GGAQALK--------AVLKYGPVLTQAGRSNEEIVHV
Burkholderia_sp._B14_(SIT71710  QALKAVLEHGPTLDERGFTLTDIVKMAGNIGGAQALKTVLGHGPTLRQRDLSLIDIVEIASN-GGAQALK--------AVLKYGPVLTQAGRSNEEIVHV
Burkholderia_sp._B14_(SIT64981  QALKAVLEHGPTLDERGFTLTDIVKMAGNIGGAQALKVVLEHGPTLRQRDLSLIDIVEIASN-GGAQALK--------AVLKYGPVLIQAGRSNEEIVQV
B._rhizoxinica_B1_(E5AW45)      QALKAVLEHGPTLRQRGFNLIDIVEMASNTGGAQALKTVLEHGPTLRQRDLSLIDIVEIASN-GGAQALK--------AVLKYGPVLMQAGRSNEEIVHV
Burkholderia_sp._B2_(MN840542)  QALKAVLEHGPTLRQRGFNLIDIVEMASNTGGAQALKTVLEHGPTLRQRDLSLIDIVEIASN-GGAQALK--------AVLKYGPVLMQAGRSNEEIVHV
Burkholderia_sp._B6_(MN840543)  QALKAVLEHGPTLRQRGFNLIDIVEMASNTGGAQALKTVLEHGPTLRQRDLSLIDIVEIASN-GGAQALK--------AVLKYGPVLMQAGRSNEEIVHV
Burkholderia_sp._B3_(MN840545)  QALKAVLEHGPTLDERGFTLTDIVKMAGNIGGAQALKAVLEHGPTLRQRDLSLIDIVEIASN-GGAQALK--------AVLKYGPVLTQAGRSNEEIVHV
Burkholderia_sp._B7_(MN840544)  QALKAVIEHGPTLDERGFTLTDIVKMAGNNGGAQALKVVLEHGPTLRQRGFSQTAIVKMAGNIGGAQALK--------AVLKYGPVLMQAGRSNEEIVNV
Burkholderia_sp._B14_(SIT64975  QALKAVIEHGPTLDERGFTLTDIVKMAGNNGGAQALKVVLEHGPTLRQRGFSQTAIVKMAGNIGGAQALK--------AVLKYGPVLMQAGRSNEEIVNV
B._rhizoxinica_B1_(E5AW43)      ----AVAMHGSTLCERGYCRTDIAKIAGNGGGAQALKAIVMHGPTLCERGYSRTDIVKIADNNGGAQALK--------AVFEHGPALTQAGRSNEDIVNM
Burkholderia_sp._B2_(MN840547)  ----AVAMHGSTLCERGYCRTDIAKIAGNGGGAQALKAIVMHGPTLCERGYSRTDIVKIADNNGGAQALK--------AVFEHGPALTQAGRSNEDIVNM
Burkholderia_sp._B6_(MN840546)  ----AVAMHGSTLCERGYCRTDIAKIAGNGGGAQALKAIVMHGPTLCERGYSRTDIVKIADNNGGAQALK--------AVFEHGPALTQAGRSNEDIVNM
Burkholderia_sp._B4_(MN840549)  -------------------------------------------PTLCERGYSRTDIVKITDNNGGAQALK--------AVFEHGPAFTQAGRSNEDIVDM
Burkholderia_sp._B13_(SIT73247  -------------------------------------------PTLCERGYSRTDIVKITDNNGGAQALK--------AVFEHGPAFTQAGRSNEDIVDM
Burkholderia_sp._B7_(MN840548)  -------------------------------------------PTLCERGYSRTDIVKIADNNGGAQALK--------AVFEHGPALTQAGRSNEDIVDM
AvrXa10_X._oryzae_(Q56830.1)    AHVVRVLGFFQSHSHPAQAFDDAMTQFGMSRHGLAQLFRRVGVTELEARYGTLPPASQRWDRILQASGMKRVKPSPTSAQTPDQASLHAFADSLERDLDA
AvrBs3_X._euvesicatoria_(P1472  AQVVRVLGFFQCHSHPAQAFDDAMTQFGMSRHGLLQLFRRVGVTELEARSGTLPPASQRWDRILQASGMKRAKPSPTSTQTPDQASLHAFADSLERDLDA
Brg11_R._solanacearum_(Q8XYE3.  QELVAVLHFFRAHQQPRQAFVDALAAFQATRPALLRLLSSVGVTEIEALGGTIPDATERWQRLLGRLGFR----PATGAAAPSPDSLQGFAQSLERTLGS
Brg11_R._solanacearum_(Q68A49.  QELVAVLHFFRAHQQPRQAFVDALAAFQTTRPALLRLLSSVGVTEIEALGGTIPDATERWQRLLGRLGFR----PATGAAAPSPDSLQGFAQSLERTLGS
```

```
                                          1110      1120
                                  ....|....|....|....|....|..
B.rhizoxinica_B1_(E5AV36)         AAR------RGGAGRIRKMVAPLLERQ
Burkholderia_sp._B2_(MN891944)    AAR------RGGAGRIRKMVAPLLERQ
Burkholderia_sp._B4_(MN840538)    AAR------RGGAGRIRKMVASLLGGN
B._endofungorum_B5_(MN891945)     AAR------RGGAGRIRKMVAPLLERQ
Burkholderia_sp._B6_(MN840537)    AAR------RGGAGRIRKMVAPLLERQ
Burkholderia_sp._B7_(MN840539)    AAR------RGGAGRIRKMVAPLLGGN
Burkholderia_sp._B7_(MN840540)    AAR------RGGAGRIRKMVAPLLGGN
Burkholderia_sp._B8_(MN840541)    AAR------RGGAGRIRKMVAPLLG--
Burkholderia_sp._B13_(SIT73265    AAR------RGGAGRIRKMVASLLGGN
Burkholderia_sp._B14_(SIT71710    AAR------RGGAGRIRKMVAPLLGGN
Burkholderia_sp._B14_(SIT64981    AAR------RGGAGRIRKMVAPLLGGN
B._rhizoxinica_B1_(E5AW45)        AAR------RGGAGRIRKMVALLLERQ
Burkholderia_sp._B2_(MN840542)    AAR------RGGAGRIRKMVALLLERQ
Burkholderia_sp._B6_(MN840543)    AAR------RGGAGRIRKMVALLLERQ
Burkholderia_sp._B3_(MN840545)    AAR------RGGAGRIRKMVASLLGGN
Burkholderia_sp._B7_(MN840544)    AAR------RGGAGRIRKMVAPLLGRQ
Burkholderia_sp._B14_(SIT64975    AAR------RGGAGRIRKMVAPLLGRQ
B._rhizoxinica_B1_(E5AW43)        AAR------TGAAGQIRKMAAQLSGRQ
Burkholderia_sp._B2_(MN840547)    AAR------TGAAGQIRKMAAQLSGRQ
Burkholderia_sp._B6_(MN840546)    AAR------TGAAGQIRKMAAQLSGRQ
Burkholderia_sp._B4_(MN840549)    AAR------IGAAGQIRKMAAQLSKRQ
Burkholderia_sp._B13_(SIT73247    AAR------IGAAGQIRKMAAQLSKRQ
Burkholderia_sp._B7_(MN840548)    AAR------IGAAGQIRKMAAQLSKRQ
AvrXa10_X._oryzae_(Q56830.1)      PSPMHEGDQTRASSRKRSRSDRAVTGP
AvrBs3_X._euvesicatoria_(P1472    PSPMHEGDQTRASSRKRSRSDRAVTGP
Brg11_R._solanacearum_(Q8XYE3.    PG--MAGQSACSPHRKRPAETAIAPRS
Brg11_R._solanacearum_(Q68A49.    PG--MAGQSACSPHRKRPAETAIAPRS
```

**Supplementary Figure 4. Confirmation of *Burkholderia rhizoxinica* wild-type (B1 WT) and transcription-activator like effector (BATs) mutants by PCR.** *Bat* genes were individually deleted to generate single mutants (**a,** Δ*bat1*::Apra$^r$, **b,** Δ*bat2*::Kan$^r$, and **c,** Δ*bat3*::Kan$^r$). Multi-deletions of *bat* genes were performed to generate double (**d,** Δ*bat2_bat3*::Kan$^r$) and triple (**e,** Δ*bat1*::Apra$^r$-Δ*bat2_bat3*::Kan$^r$) mutant strains. The PCR products of the mutant strain correspond to amplicons A and B; whereas products C and D are amplified from the wild-type. **f,** Mutants were checked for the absence of contaminating wild-type by amplifying an internal fragment (amplicons I) form the wild-type gene of interest. The removal of the knock-out vector was confirmed by amplification of the counter selection marker gene *pheS* (amplicons pheS). PCR products were obtained using control primers listed in Supplementary Table 5. Bands corresponding to the expected size are indicated by asterisks (*).

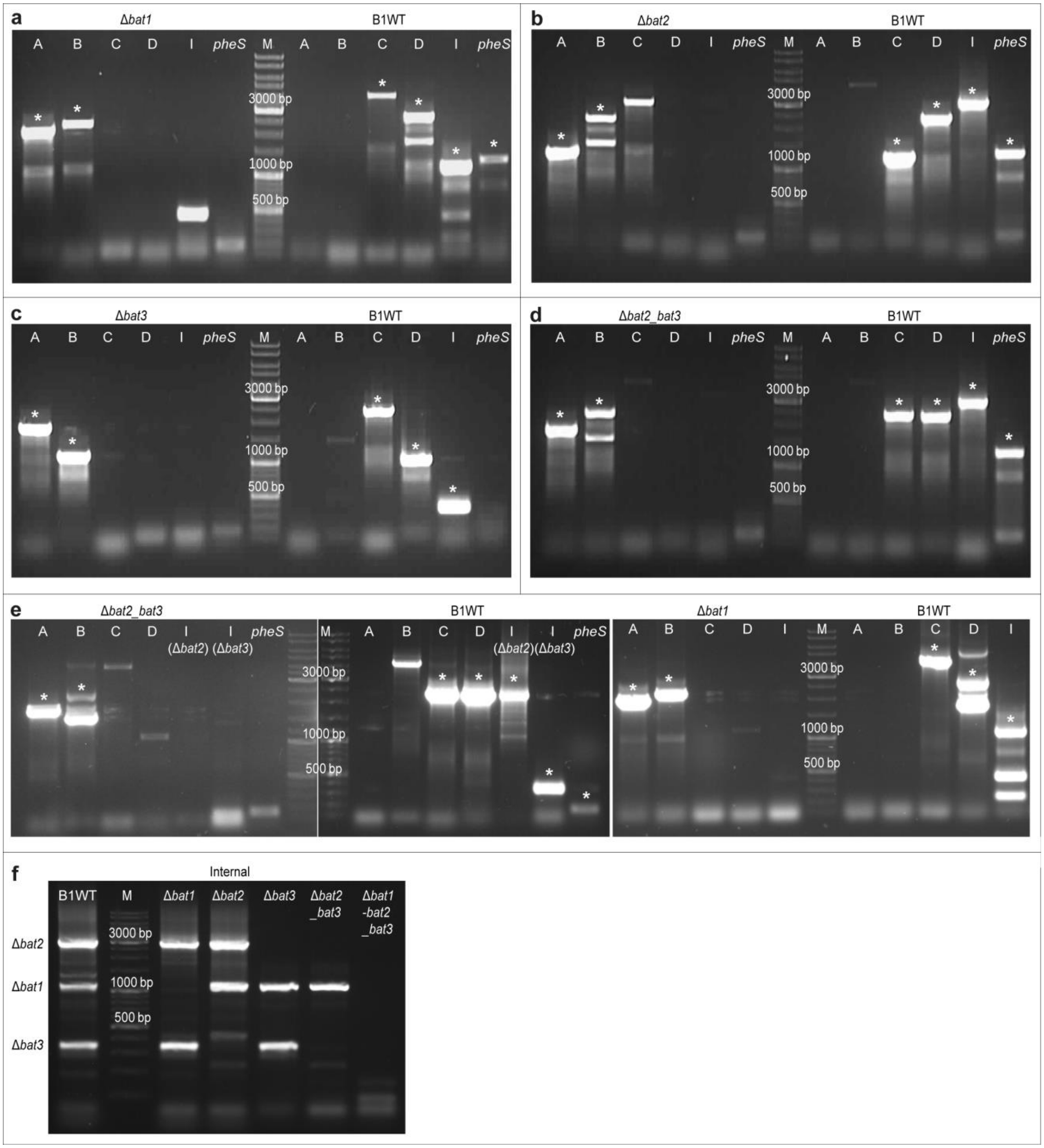

**Supplementary Figure 5. Phenotypic observation of *Rhizopus microsporus* co-cultivated with *Burkholderia rhizoxinica* mutant strains in bacterial-fungal interaction (BFI) devices.** Endosymbiont-free *R. microsporus* ATCC62417/S was co-incubated with *B. rhizoxinica* transcription-activator like effector (BAT) mutant strains (Δ*bat1*::Apra$^r$, Δ*bat2*::Kan$^r$, and Δ*bat3*::Kan$^r$), type 2 secretion system mutant strains (Δ*sctC*::Kan$^r$ and Δ*sctT*::Kan$^r$), or rhizoxin-deficient mutants (Δ*rhiG*::Kan$^r$). Bacterial strains were stained with SYTO9 prior to co-incubation. Microscopic images taken 48 hours post infection (hpi) revealed four different morphotypes. Morphotypes were classified as follows: 1: vegetative side hyphae; 2: vegetative main hyphae (trunk hyphae); 3: vegetative empty side hyphae; 4: abortive sporangiophore. Arrow heads indicate the presence of septa. Scale bar: 10 µm.

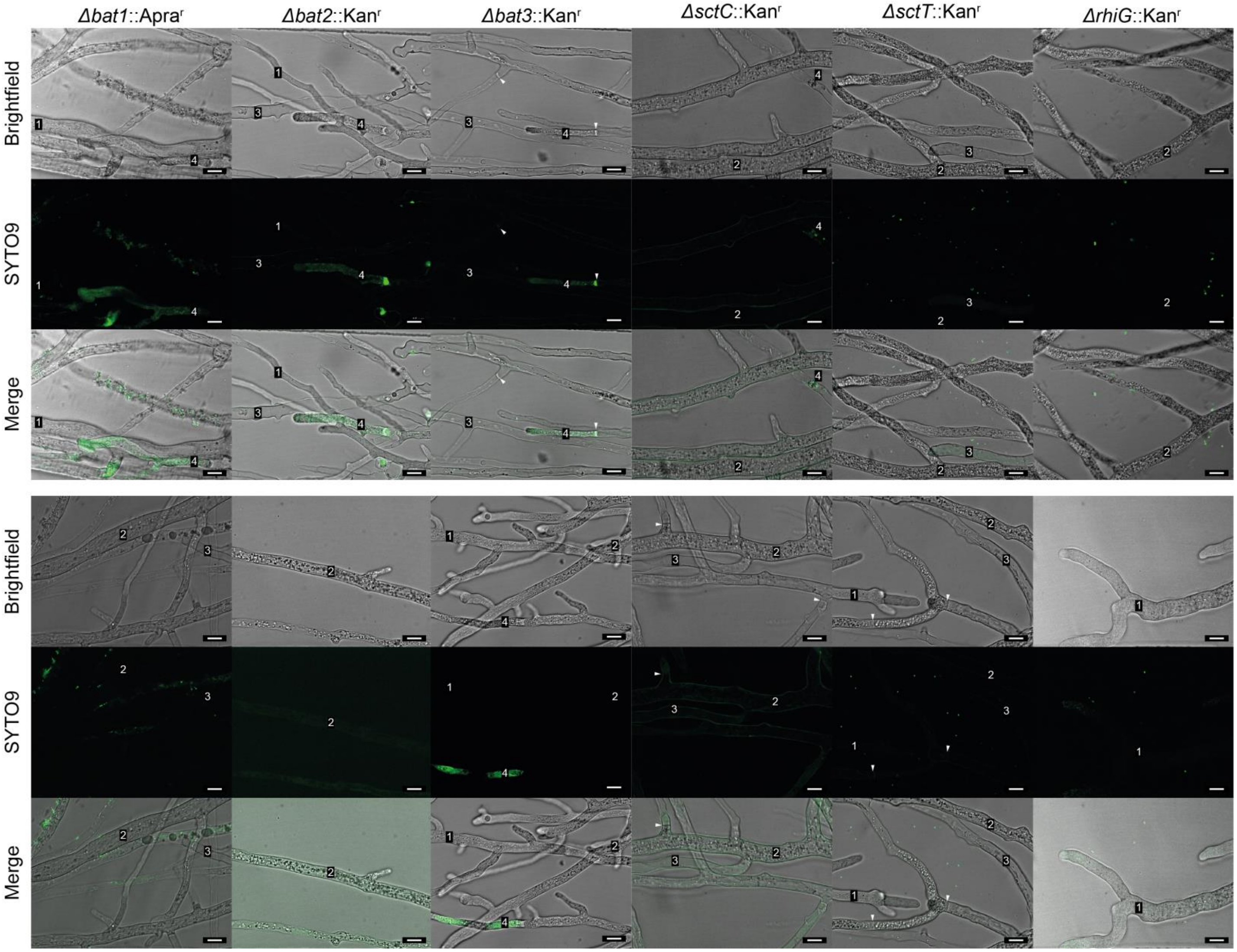

**Supplementary Figure 6. *Burkholderia rhizoxinica* type 3 secretion system (T3SS) mutant strains fail to re-infect *Rhizopus microsporus*.** Endosymbiont-free *R. microsporus* ATCC62417/S was co-incubated with *B. rhizoxinica* T3SS mutant strains (Δ*sctC*::Kan$^r$ or Δ*sctT*::Kan$^r$) for 48 hours. Bacterial cells were stained with SYTO9 prior to co-incubation. Following fluorescence microscopy at 485/498 nm (SYTO9). The integrated density (product of area and mean grey value) per inoculated cell number was calculated using Fiji[4], and then plotted as percent of the positive control (*R. microsporus* ATCC62417/S co-incubated with wild-type *B. rhizoxinica*; % of B1WT). N = 3 biological replicates (16 technical replicates) ± one SEM. One-way ANOVA with Tukey's multiple comparison test (****p<0.05*, significant differences are indicated for morphotype 4, Supplementary Table 12).

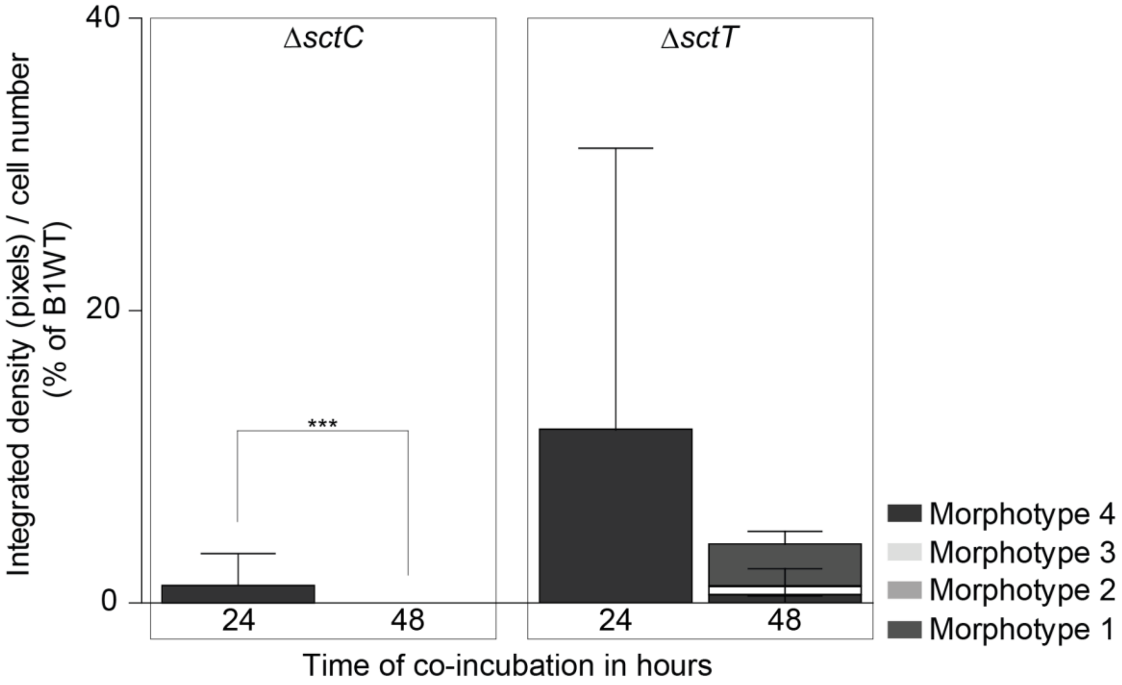

**Supplementary Figure 7. Rhizoxin-deficient *Burkholderia rhizoxinica* mutant strains re-infect *Rhizopus microsporus* similar to wild-type *B. rhizoxinica*. a,** Endosymbiont-free *R. microsporus* ATCC62417/S was co-incubated with *B. rhizoxinica* rhizoxin mutant strains (Δ*rhiG*::Kan$^{r}$) for 48 hours. Bacterial cells were stained with SYTO9 prior to co-incubation. Following fluorescence microscopy at 485/498 nm (SYTO9). The integrated density (product of area and mean grey value) per inoculated cell number was calculated for all morphotypes combined (Total) and for individual morphotypes (Morphotypes) using Fiji[4], and then plotted as percent of the positive control (*R. microsporus* ATCC62417/S co-incubated with wild-type *B. rhizoxinica*; % of B1WT). **b,** The same images were used to calculate the number of septa per area (in µm) and the number of septa per cell number using Fiji, and then plotted as percent of the positive control (*R. microsporus* ATCC62417/S co-incubated with wild-type *B. rhizoxinica*; % of B1WT). N = 3 biological replicates (16 technical replicates) ± one SEM. One-way ANOVA with Tukey's multiple comparison test (***$p<0.05$, Supplementary Table 13) **c,** Endosymbiont-free *R. microsporus* and *B. rhizoxinica* (Δ*rhiG*::Kan$^{r}$) were stained with SYTO9 and co-incubated in a bacterial-fungal interaction device. Reinfection was monitored over time using fluorescence microscopy. Scale bar: 10 µm.

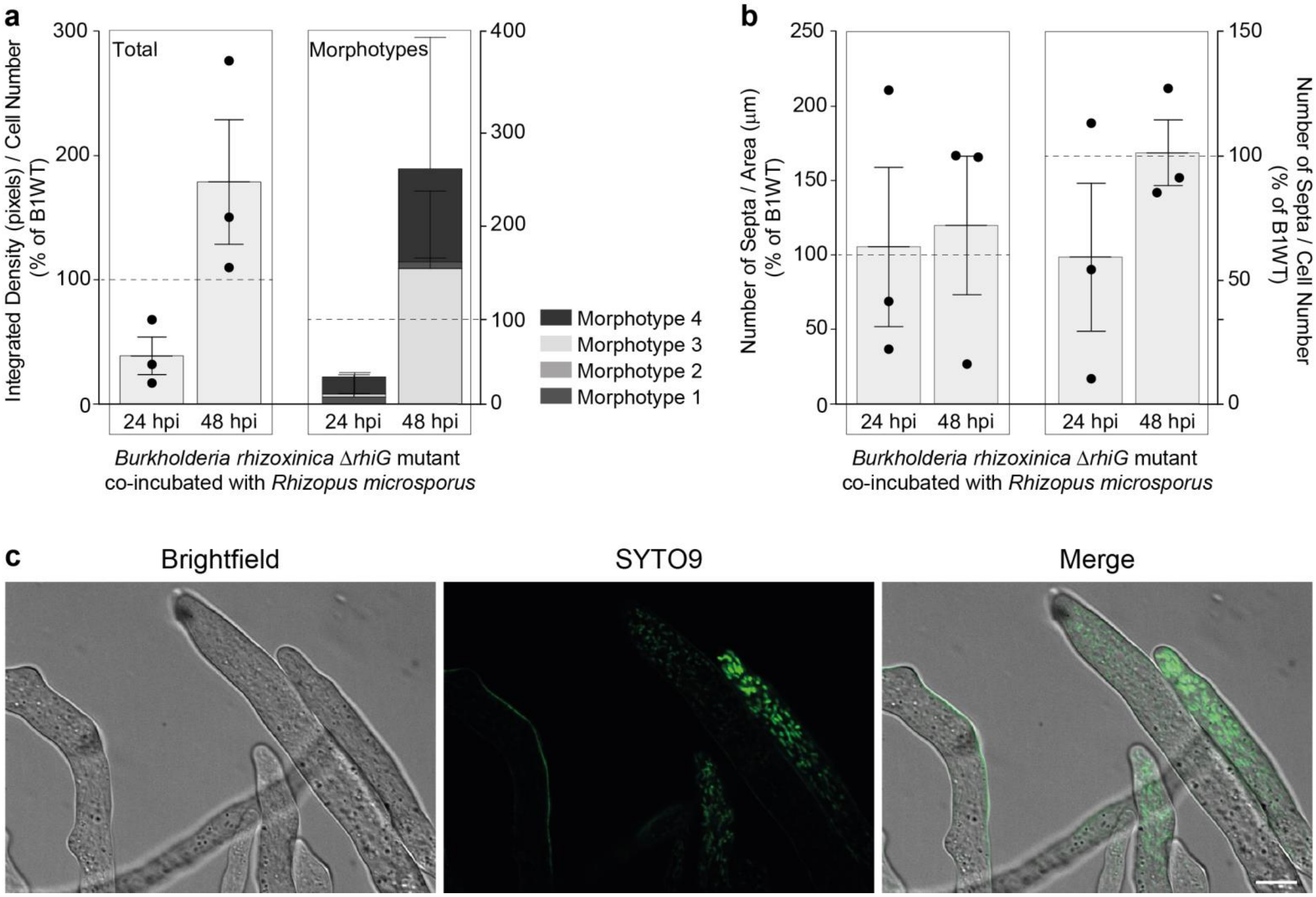

**Supplementary Figure 8. The majority of axenic wild-type *B. rhizoxinica* (B1 WT) and *Burkholderia* transcription-activator like effector (BAT) mutant strains are alive.** B1 WT and BAT mutant strains (*Δbat1*::Apra$^r$, *Δbat2*::Kan$^r$. *Δbat3*::Kan$^r$) were visualised with LIVE/DEAD BacLight fluorescent dyes. Green fluorescence (SYTO9) indicates that bacteria are alive, while red fluorescence (propidium iodide) is indicative of dead bacteria. Scale bar: 10 µm.

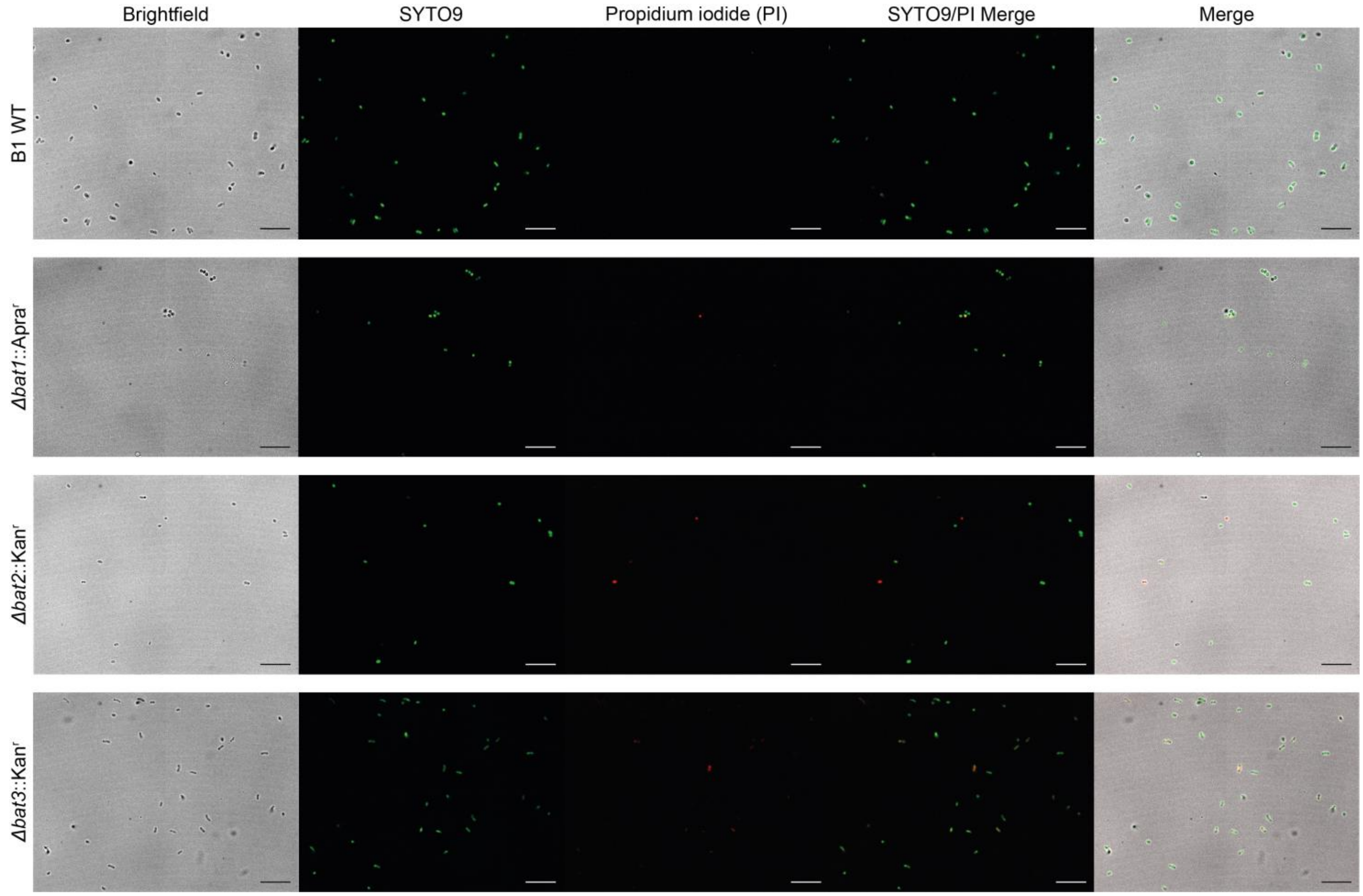
